# FiberBrowser: a unified framework for single-molecule genomics

**DOI:** 10.64898/2026.10.29.762797

**Authors:** Thomas W. Tullius, Christopher Catalano, Jane Ranchalis, Jie Hu, Mitchell R. Vollger, Andrew B. Stergachis, Michael S. Levine

**Author notes:** These authors jointly supervised this work (Andrew B. Stergachis, Michael S. Levine).

## Abstract

Single-molecule footprinting methods are rapidly transforming chromatin biology. They resolve nucleosome positioning, regulatory-factor occupancy, and accessibility on individual DNA fibers, revealing combinatorial heterogeneity that bulk measurements average away. Emerging footprinting methods leverage diverse enzyme systems and sequencing platforms, necessitating tools that can accommodate these systems. We present FiberBrowser, a unified platform that converts these diverse datasets into a common per-molecule representation of nucleosomes, accessible DNA, and subnucleosomal footprints for interactive and scripted analyses. Its inference engine, FiberHMM, weighs each observation against the assay’s informative sites and sequence-specific modification probabilities, separating missing information from protection in four assays. FiberBrowser clusters, sorts and compares these calls across thousands of molecules in seconds, linking chromatin architecture to haplotype, CpG methylation and non-reference structural variants on the same molecules. Because the representation is independent of chemistry, models also transfer between assays. A classifier trained on Fiber-seq footprints identifies 12 scDAF-seq cells as lymphoblastoid without retraining, whereas a model trained on raw modification patterns fails to transfer. FiberBrowser thus provides a unified, expandable analytical framework for single-molecule chromatin footprinting.

## Introduction

Single-molecule footprinting resolves nucleosome positioning, polymerase and transcription factor occupancy, and accessibility together on individual chromatin fibers, revealing combinatorial states that bulk short-read epigenomic methods average away^1,2^. Accessible DNA can be labeled by adenine methyltransferases (Fiber-seq, SAMOSA and others^3–6)^, cytosine methyltransferases (SMF, NOMe-seq and others^7–12)^ or cytosine deaminases (DAF-seq, FOODIE and others^13–16)^, and the marks are read by short-read, PacBio or Oxford Nanopore (ONT) sequencing.

These assays read the same chromatin states, but their data differ. Each enzyme targets different bases with its own sequence preferences, and methyltransferase assays sequenced on PacBio without amplification report both strands, whereas ONT and amplified deaminase assays report only one^17,18^. Resolution therefore depends on how densely the target bases occur and on whether both strands are read. For example, some of the cytosine deaminases show strong dinucleotide preferences^19^, and ONT sequencing of an adenine methyltransferase-marked sample will have half the resolution of the same sample sequenced with PacBio HiFi. Even within accessible DNA, enzymes modify only a context-dependent fraction of eligible bases. These differences matter because a footprint is inferred from the absence of modification, so an unmodified stretch can mean either that a protein protected the DNA or that the assay had few chances to report a modification there. A caller that ignores this distinction reports false footprints where informative bases are sparse and misses short footprints that dense data support. Accurate footprinting therefore requires weighing every observation against where the assay could have reported a modification and how likely it was to do so.

Analysis tools have largely been developed alongside individual assays and for specific use cases. Fibertools (ft) calls m6A and manipulates, annotates and summarizes Fiber-seq data, and its simple heuristic-based nucleosome caller underlies Fiber-seq inference of regulatory elements (FIRE)^20,21^. IDLI classifies nucleosomal and subnucleosomal footprints in SAMOSA data^4,22^, and SingleMoleculeFootprinting and FootprintCharter call footprints from CpG and GpC methylation at selected loci^10,23–25^. Analyses accompanying FOODIE^14^, FootTrack for cFOOT-seq^16^ and HiddenFoot^26^ score protection at predefined intervals, in aggregate signal or at specified binding sites. Most of these tools are tied to one enzyme and platform, and several require motif or locus annotations in advance, which limits their use genome-wide and across diverse chromatin types. What remains missing is a factor-agnostic footprint caller that returns nucleosome and footprint calls on individual molecules across chemistries, with an explicit model of what each assay can observe.

Visualization presents a second gap. General-purpose genome browsers such as IGV and the UCSC Genome Browser can display base modifications on individual long reads^27,28^, but they are organized around aggregate signal tracks. Two regulatory elements accessible on half of molecules give the same aggregate profile whether they open together or mutually exclusively. Distinguishing these cases requires grouping molecules by accessibility or footprint pattern, relating the groups to haplotype, CpG methylation or structural variation on the same molecules, and comparing them across samples. General-purpose browsers offer little support for these analyses, do not display chromatin within sequence absent from the reference, and perform poorly with deep targeted datasets of thousands of multi-kilobase reads at one locus. Dedicated packages plot methylation or occupancy on individual molecules^29–32^, but they are largely restricted to single assays and static views of selected loci.

Here we address both gaps. First, we generalize FiberHMM, our previously introduced footprint caller for Hia5 Fiber-seq^33^, into a framework that calls nucleosomes, accessible patches and subnucleosomal footprints for four chemistry–platform combinations. It models the informative bases of each assay, including the deamination that SsDddA introduces at specific, rotationally phased positions within nucleosomal DNA. For single-cell DAF-seq generated with SsDddA^15,34^, FiberHMM reconstructs both strands of individual fibers so that the assay can be calibrated directly in chromatin, and it infers CpG island methylation on the same molecules. Second, we build FiberBrowser, a genome browser and analysis platform that treats each read as an individual chromatin molecule. It reads FiberHMM’s annotations directly and combines interactive clustering of molecular states, allele-specific footprinting and inspection of non-reference structural variants with reproducible graphical and Python workflows, so that footprint calls can be queried and compared in real time across molecules, haplotypes and samples.

## Results

### FiberHMM calls nucleosomes and footprints across single-molecule assays

FiberHMM (fiberhmm-call) identifies nucleosomes, accessible modification-sensitive patches (MSPs) and subnucleosomal footprints on each sequencing read (i.e., molecule) of an aligned BAM, enabling a per-molecule representation of chromatin features (Fig. 1a). These per-molecule features, as well as their evidence scores, are then encoded directly in the aligned BAM using Molecular Annotation (MA) tags (https://github.com/fiberseq/Molecular-annotation-spec; Supplementary Table 1), enabling the genetic and epigenetic information of each molecule to be co-encoded within a widely adopted file format. These BAM files with MA tags can then be directly read by the FiberBrowser interface as well as other tools such as Fiber-seq inference of regulatory elements (FIRE)^21^ and IGV (version 3.0 and later), without format conversion.

**Figure 1.**
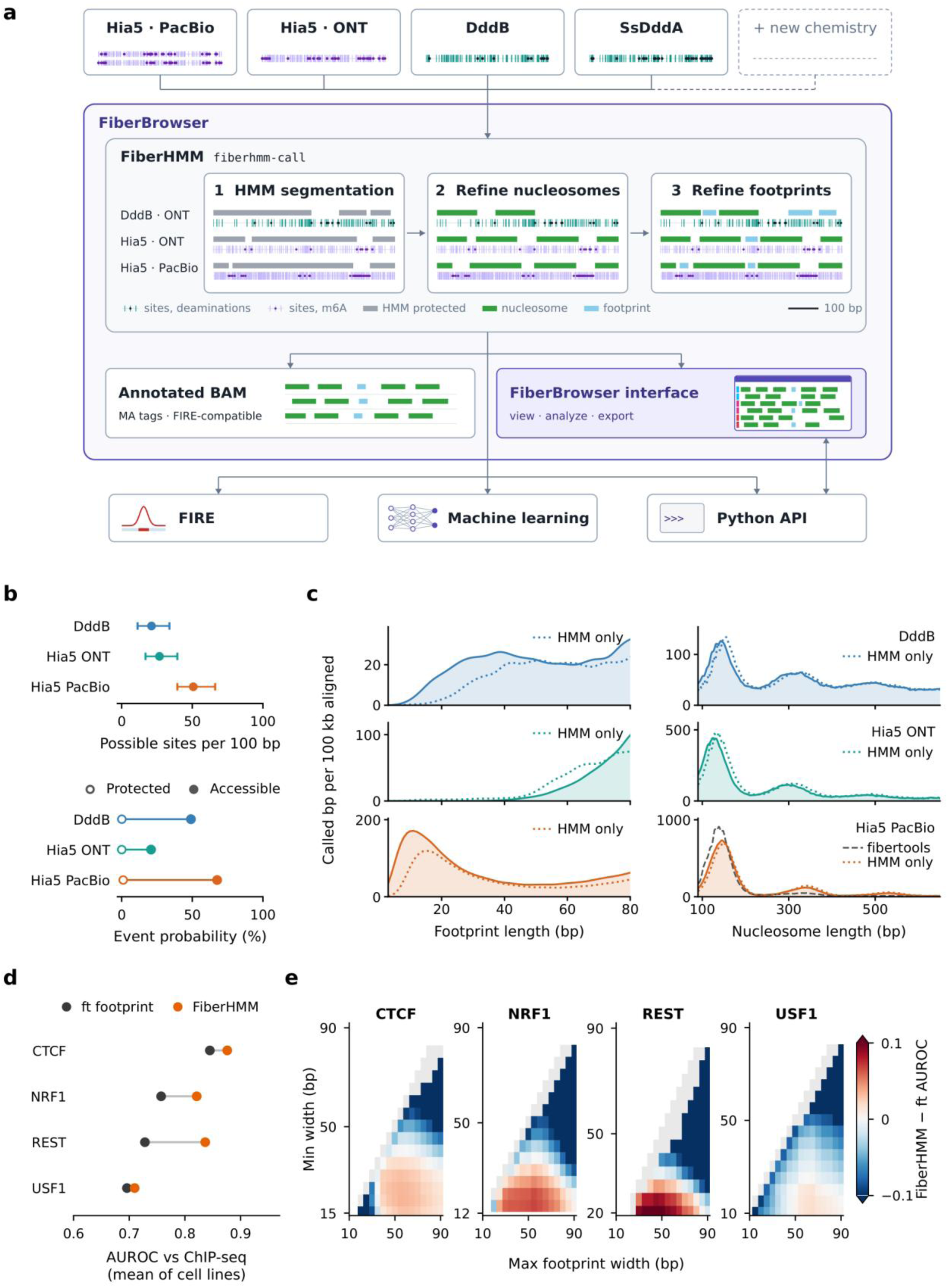
FiberHMM calls nucleosomes and footprints across single-molecule chromatin assays. a, The FiberBrowser framework. Reads from each supported chemistry (Hia5 PacBio, Hia5 ONT, DddB and SsDddA; new chemistries are added with a new preset) are processed by FiberHMM (fiberhmm-call) in three steps, shown on one real 600-bp window for each of the three illustrated assays (DddB ONT at *hb* and Hia5 ONT in *Drosophila* embryos; Hia5 PacBio in K562; scale bar, 100 bp): HMM segmentation of each molecule into protected (gray) and accessible DNA, nucleosome refinement at linkers (green) and footprint refinement at LLR 5 (sky blue). Ticks, enzyme informative sites; black dots, deaminations; purple dots, m6A. Calls are written to an output BAM as Molecular Annotation (MA) tags alongside the raw observations. They are read by the FiberBrowser interface (Fig. 3), which runs the same engine and is driven by the Python API, and by downstream analyses including FIRE and machine learning (Fig. 5). b, Median possible sites per 100 bp (10th–90th percentiles) and mean event probabilities in protected and accessible DNA across the informative sites of the plotted molecules. c, Call-length distributions at LLR 5 for footprint-scale (3–80 bp) and nucleosome-scale (90–650 bp) calls, as called bp per 100 kb aligned per 1-bp bin (y-axes scaled per assay; 81–89 bp not shown). DddB, 86,591 molecules from 22 *hb*, *ind*, *scyl* and *chrb* amplicon libraries; Hia5 ONT, 11,162 *Drosophila* yw embryo molecules (2–4 h); Hia5 PacBio, 3,000 K562 molecules. Dotted curves show HMM-only calls on the same DddB, Hia5 ONT and Hia5 PacBio molecules; dashed gray, fibertools ft add-nucleosomes on the matched PacBio molecules. SsDddA, whose nucleosome calls require the rotational refinement, and whose footprint emissions come from the duplex calibration introduced in Fig. 2, is characterized in Fig. 2 and Supplementary Fig. 1b,d, Mean AUROC against cell-matched ChIP-seq for ft footprint (gray) and fiberhmm-footprint with a ChIP-calibrated width window (orange), each window learned in the other cell lines of the same factor (CTCF, five cell lines; NRF1, REST and USF1, four). e, FiberHMM minus ft footprint AUROC for each [minimum, maximum] footprint-width window, mean of cell lines; y axes start at the motif length. Gray, windows with too few positives to rank motifs (FiberHMM positives in <1% of observations or at <20% of motifs); white, minimum above maximum. Per-cell-line values are in Supplementary Fig. 4.

FiberHMM represents each molecule by the sites where the assay can report a modification and the modifications observed there. The assays differ markedly in both quantities (Fig. 1b and Supplementary Fig. 1a,b). Hia5 PacBio Fiber-seq reports about 50 possible sites per 100 bp as it is capable of identifying modified adenines on both strands, roughly twice the resolution of methods that only mark a specific base on one strand, such as targeted cytidine deaminase-based methods or Fiber-seq sequenced using simplex ONT. Specifically, in Fiber-seq sequenced using simplex ONT, stretches of DNA longer than 10 bp can appear unmodified simply due to an absence of adenine bases. Thus, processing of these data must take into account multiple features that are common to all assays, as well as features unique to a specific assay (Fig. 1a). To accomplish this, FiberHMM employs a two-state hidden Markov model (HMM) that segments each molecule into protected domains and accessible MSPs. Two log-likelihood-ratio (LLR) refinement steps then score each candidate call against the probability that its informative sites would have been modified if it were accessible. Refinement splits long HMM-protected domains into individual nucleosomes and calls short footprints between them where the data support protection (Fig. 1a).

To select a default LLR threshold, we simulated protected spans of 8–60 bp in motif-centered genomic sequences, with matched fully accessible controls, from each assay’s accessible- and protected-state probabilities (Supplementary Fig. 2a,b,d). The threshold trades sensitivity against specificity, because lower values recover more short footprints but also call more footprints in accessible DNA (Supplementary Fig. 2a,b). We use LLR 5, the lowest integer threshold at which calls in accessible controls fell below 2% for every chemistry, including SsDddA. Users can change it with fiberhmm-call--min-llr. In held-out PacBio-sequenced, Hia5-treated naked DNA, calls at LLR 5 covered 3.6% of fixed windows, mostly short calls on under-labeled molecules (Supplementary Fig. 3a); the DddB and Hia5 ONT naked-DNA controls behaved similarly (Supplementary Fig. 3b). At this threshold, recovery follows each assay’s information density. Hia5 PacBio recovered 91% of 10-bp targets, DddB recovered 86% of 40-bp targets, and single-strand Hia5 ONT recovered 26% even at 50 bp.

The same pattern appears in endogenous calls (Fig. 1c). Hia5 PacBio yields abundant footprints below 20 bp and a sharp mononucleosome peak near 145 bp which closely matches the fibertools nucleosome caller on the same molecules. DddB and Hia5 ONT in *Drosophila* embryos yield fewer short footprints and broader nucleosome peaks, including more dinucleosome-sized calls where linker evidence is sparse. In DddB and Hia5 PacBio, refinement recovers short footprints that the HMM merged into protected domains, whereas in sparse Hia5 ONT data it removes short HMM segments that lack support. FIRE run on FiberHMM segmentation reproduces FIRE on the fibertools segmentation of the same reads, reaching 95–98% directional concordance among the most confident calls (Supplementary Fig. 1e,f).

To benchmark the performance of FiberHMM for calling transcription factor footprints, we first built fiberhmm-footprint, and directly compared the footprint calls to those of fibertools footprint (ft footprint). Similar to ft footprint, with fiberhmm-footprint a molecule counts as bound at a given motif when: (1) an MSP on that molecule spans the complete motif; (2) a footprint call covers the complete motif; and (3) the motif carries no m6A. Because FiberHMM calls footprints with defined edges, fiberhmm-footprint can also restrict the width of the footprint that counts as a binding event. Without a width restriction the two callers agree. A label-free cap of the 95th percentile of footprint widths at motifs improved how well the per-motif bound fraction separated ChIP-supported (ENCODE^35^) from unsupported motifs for CTCF, NRF1 and REST in every cell line tested (Supplementary Fig. 4a). Where ChIP-seq is available, the width window can instead be calibrated to maximize prediction accuracy while also reporting each factor’s characteristic footprint size (Fig. 1d,e, Supplementary Fig. 4, Supplementary Note 1 and Supplementary Tables 2 and 3).

### Rotational refinement resolves SsDddA nucleosomes

Hia5 and DddB fit this framework directly, with emission probabilities estimated from naked-DNA and untreated controls. SsDddA, used in targeted DAF-seq and single-cell DAF-seq (scDAF-seq), required two extensions. First, SsDddA deaminates not only accessible DNA but also specific positions within nucleosomes. These deaminations within presumed nucleosomes oscillate with a 10.3-bp period around inferred nucleosome dyads (Fig. 2a), matching the helical repeat of DNA on the histone surface and the rotational pattern of nuclease cleavage of nucleosomal DNA^36,37^. FiberHMM uses these internal deaminations to identify nucleosome cores using relaxed emission probabilities, keeping deaminations that fit the rotational phase inside the particle (fiberhmm-call--enzyme ddda). Truncated calls are extended to mononucleosome size while the accessible flanks are preserved (Fig. 2b).

**Figure 2.**
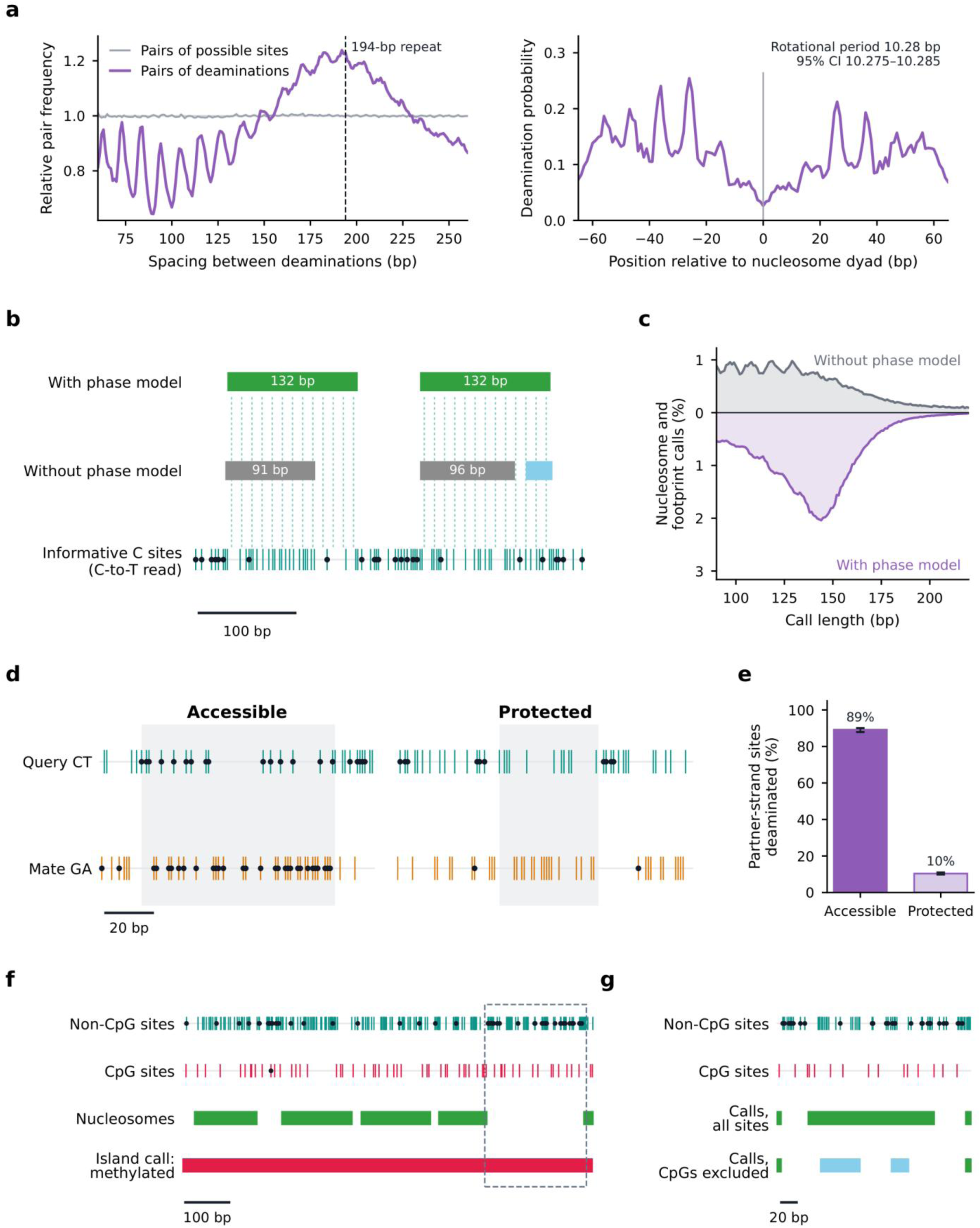
SsDddA rotational inference, duplex validation and CpG-island methylation. a, Deamination-to-deamination spacing (left) and deamination probability relative to inferred nucleosome dyads (right), with deamination pairs and possible-site pairs each normalized to their own 121-bp moving average (left) and deaminations per possible site around dyads placed by the phase model (right). Dashed guide, 194-bp repeat; rotational period 10.28 bp (95% CI 10.275–10.285, molecules resampled). b, One scDAF-seq molecule called without and with the phase model; ticks, the informative C sites of this C-to-T read after CpG masking and keep-one thinning, identical in both calls; sky blue, footprint call; dashed lines mark the 10.28-bp period centered on each phase-model call, and widths are labeled for calls of at least 35 bp. c, Length distributions of nucleosome and footprint calls for 20,935 molecules without and with the phase model (2.78 and 4.79 calls per mapped kb, an increase that mainly reflects merged calls of 250 bp or more, outside the plotted range, being resolved into individual particles; Supplementary Fig. 5a,b). d, Physical duplexes with an accessible and a protected query window (shading, query window; teal/amber ticks, CT/GA sites; black dots, deaminations). Strand pairing is described in Online Methods. e, Partner-strand deamination in windows called accessible or protected on the query strand across 12 libraries and both transfer directions; error bars, library-cluster 95% CIs. f, Whole-island methylation inference for one HG002 molecule across an 890-bp CpG island (1/20 CpGs and 43/76 non-CpGs deaminated; methylated posterior 0.99997; matched HG002 LCL long-read methylation 86.4% across 59 sites). g, The dashed window in f (molecule positions 1,250–1,470), called with all sites (Calls, all sites), where unconverted methylated CpGs read as protection and the window is called as one 144-bp nucleosome, and with CpGs excluded (Calls, CpGs excluded), which gives two footprints of 46 and 21 bp. Scale bars: b, f, 100 bp; d, g, 20 bp. Whole-island diagnostics, Supplementary Fig. 7a–d; held-out validation, Online Methods.

Across 20,935 molecules, the phase model increases the number of nucleosome and footprint calls per kilobase 1.7-fold (Fig. 2c and Supplementary Fig. 5a,b). Most of the gain comes from long merged calls being resolved into individual nucleosomes. Most nucleosomes carry at least one internal deamination, often on one side of the dyad. Because these deaminations fall at fixed rotational positions relative to the dyad, even a single event helps constrain the dyad position on an individual molecule. Furthermore, the internal deamination rate varies with nucleosome position across the *NAPA* and *UBA1* amplicons (Supplementary Fig. 5c–e), suggesting that some nucleosomes are more prone than others to internal SsDddA deamination. DddB showed little internal deamination at the enzyme concentrations used, so its nucleosomes are called with the standard refiner. Because adjacent same-strand SsDddA targets convert together, FiberHMM counts each such run once (Supplementary Fig. 5f–h).

### Strand duplexes calibrate SsDddA footprint calling

Unlike other stranded assays, scDAF-seq can reliably capture both strands of the same original DNA template. As most protein occupancy events are anticipated to occupy both strands at a given binding element, scDAF-seq data thereby enable the calibration of footprint detection on each strand, using the opposite strand as a comparator. This is required because SsDddA has no usable naked-DNA control: fully accessible DNA accumulates too many uridines to amplify. To test this, we calibrated accessible- and protected-state probabilities in chromatin using physical duplexes from scDAF-seq data, in which the separately amplified C-to-T and G-to-A strands of one fiber are both sequenced (Fig. 2d). Because the two strands are deaminated and read independently, a window called accessible or protected on one strand provides a fixed region in which to measure deamination on its partner. Strands can be readily paired using heterozygous variants that distinguish DNA arising from either of the two haplotypes within a single cell. However, without chaining reads together, sequence alone could not decide the mate for many duplexes.

To address this, we developed a tool within FiberHMM that pairs the strands by their nucleosome patterns, using deamination-insensitive sequence differences where they exist (fiberhmm-pair; Supplementary Fig. 6a–c), enabling pattern-based pairing to recover duplexes that haplotype and sequence cannot assign. Where the two homologs differ in sequence, so that sequence can identify the mate independently, pattern-based pairing agreed with it in 96% of assessable cases (Supplementary Fig. 6c). In windows called accessible on one strand, 89% of informative sites on the partner strand were deaminated, compared with 10% in windows called protected (Fig. 2e), directly establishing empirical accessible- and protected-state deamination rates. The contrast is consistent across 12 libraries and whichever strand serves as the query, and Hia5-defined accessible regions give similar rates in densely deaminated SsDddA neighborhoods (Supplementary Fig. 1c,d). Calling both strands together also fills each strand’s gaps with its partner’s informative sites, more than doubling recovery of short footprints using scDAF-seq (Supplementary Fig. 6d).

### SsDddA deamination reports CpG-island methylation

SsDddA deamination is mildly to moderately reduced by CpG methylation^15,16^, raising the possibility that SsDddA-based footprinting could be used to identify the methylation state of CpG islands. Specifically, methylated CpGs preferentially remain undeaminated even when they are not protein bound. Although this effect is too weak to call methylation status at single CpGs, we reasoned that it could classify entire CpG islands as methylated or unmethylated directly from SsDddA-based DAF-seq data. FiberHMM first segments each molecule with all CpGs masked and then compares CpG with non-CpG deamination within the called MSPs of each island (fiberhmm-tag-m5c). For each molecule, the deamination rate at non-CpG cytosines within those MSPs sets an accessible-state baseline, and the CpG outcomes are scored against calibrated methylated and unmethylated emission models with equal prior odds. Across one 890-bp island, for example, 43 of 76 non-CpG cytosines were deaminated but only 1 of 20 CpGs, classifying the island as methylated, in agreement with independent long-read CpG methylation data (Fig. 2f,g). Genome-wide, unmethylated island calls agreed with long-read CpG methylation at 99.9% precision (Online Methods).

For footprint calling, FiberHMM therefore treats CpGs as methylated by default and uses their deamination evidence only within confidently unmethylated islands, which prevents undeaminated methylated CpGs from being read as protection. Methylated islands carry more nucleosomes and therefore fewer and shorter MSPs, but this does not confound the comparison. MSPs are called with all CpGs masked, from the far more numerous non-CpG cytosines. CpG deamination is then compared with that of the non-CpG cytosines in the same confident MSPs of the same molecule. Accordingly, CpG-specific loss persisted at matched local accessibility (Supplementary Fig. 7e–g) and across MSP lengths and nucleosome geometries (Online Methods).

### FiberBrowser turns single-molecule visualization into real-time population analysis

Conventional genome browsers are built around aggregate tracks that obscure single-molecule heterogeneity, and they display molecules without analyzing them. We therefore built FiberBrowser, an interactive visual and computational platform organized around individual chromatin fibers rather than population averages (Fig. 3a). Each dataset opens as a panel, synchronized with the others or navigated independently, in which every molecule is drawn as a row of its FiberHMM calls, with nucleosomes in green, footprints in sky blue and accessible DNA left pale (Fig. 3a, labels 2, 4 and 7). Shared annotation tracks and metaprofiles of nucleosome and footprint occupancy sit above the molecules (Fig. 3a, labels 5 and 6), and raw modifications, CpG methylation and other layers can be overlaid on the same rows. FiberBrowser’s toolbar and analysis sidebar sort, filter and cluster molecules and scan for peaks and variants, and cluster assignments are drawn beside the molecules (Fig. 3a, labels 3 and 8–10). Sessions are opened, saved and exported from the file controls (Fig. 3a, label 1).

**Figure 3.**
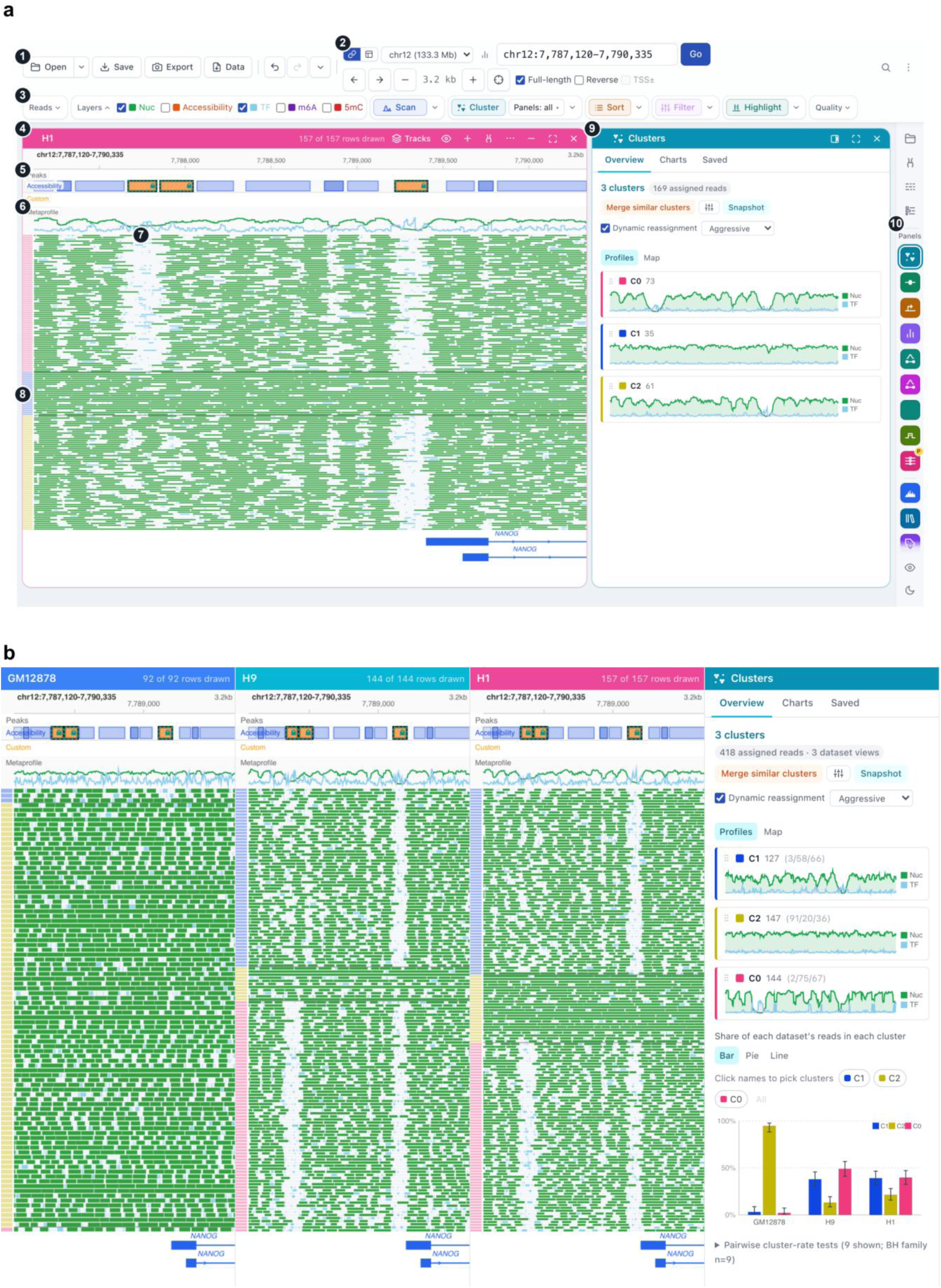
FiberBrowser turns single-molecule visualization into real-time population analysis. a, The FiberBrowser interface with one H1 Fiber-seq dataset at *NANOG* (chr12:7,787,120–7,790,335). Numbered labels mark (1) file and session controls, (2) locus and panel navigation, (3) the toolbar of molecule operations (layers, scanning, clustering, sorting and filtering), (4) the dataset panel header, (5) peak and custom-feature tracks, (6) the nucleosome (green) and footprint (sky blue) metaprofile, (7) molecule rows drawn from their FiberHMM calls (green, nucleosomes; sky blue, footprints; pale, accessible DNA), (8) the cluster-membership strip, (9) the analysis sidebar, here the Clusters panel, and (10) the rail that opens analysis panels. b, Joint clustering of lymphoblastoid (GM12878) and embryonic stem cell (H9 and H1) Fiber-seq on accessibility at the *NANOG* promoter and a nearby enhancer (chr12:7,787,120–7,790,335). Left, per-dataset molecular views with shared cluster assignments (green, nucleosomes; colored strips, cluster membership) and metaprofiles. Right, the interface’s summaries of the three clusters (C0–C2), with molecule counts (GM12878/H9/H1 in parentheses), nucleosome (green) and footprint (sky blue) metaprofiles, and cluster proportions per dataset (clustering used all 418 molecules in the region; 393 fully spanning molecules are drawn); error bars, 95% Wilson intervals with reads as units (FiberBrowser documentation).

To combine responsive display with quantitative analysis, FiberBrowser separates the rendering of the visualized reads from background computation. The view draws a sample of molecules for speed, while sorting, filtering, clustering and co-occupancy analyses use every eligible molecule in the region. Molecules from several datasets can be jointly clustered or sorted by their accessibility or footprint patterns over selected elements, showing at once how chromatin states are distributed across samples. At *NANOG*, joint clustering of GM12878, H9 and H1 Fiber-seq on accessibility at the promoter and a nearby enhancer separates three states (Fig. 3b). One state is closed at both elements and contains almost all lymphoblastoid GM12878 molecules. The two embryonic stem cell lines instead spread across all three states, with most of their molecules in the two accessible states, which differ in which elements are open. The cluster strip, per-cluster metaprofiles and composition chart display these states, and every summary can be traced back to the calls on individual molecules.

Every analysis is saved with the session, its results can be exported, and every function can be run through a Python API for scripted or headless use. The interface also includes a searchable data library with automatic quality control and tools for targeted assays, such as amplicon primer design (Supplementary Fig. 8). IGV desktop shows the first screen of a newly opened deep region sooner, but FiberBrowser redisplays regions 2.5–6.6-fold faster, draws all 50,000 molecules of a region in a similar time, and remains usable at depths where igv.js runs out of memory. Unlike IGV, FiberBrowser also analyzes every molecule in the region. At a 3,000-molecule amplicon, sorting, clustering and footprint co-occupancy of all molecules each take one to two seconds (Supplementary Fig. 9a,b; Supplementary Note 2 and Supplementary Tables 4–7).

### Dissecting allelic footprints, DNA methylation and non-reference structural variants on single fibers

A defining advantage of long-read single-molecule assays is that each read captures genetic variants, CpG methylation and chromatin footprints on the same DNA molecule. In FiberBrowser, molecules can be split by haplotype or allele, colored by methylation or grouped by a structural variant, then clustered and compared within seconds. At the imprinted *SNRPN* promoter in H9 embryonic stem cells, phasing molecules by heterozygous SNPs and overlaying CpG methylation shows the imprint directly on single molecules (Fig. 4a). Molecules arising from haplotype 1 (i.e., HP1) carry a broad, largely unmethylated accessible region across the differentially methylated region, whereas those arising from haplotype 2 (i.e., HP2) are densely methylated and packed with nucleosomes. Joint clustering placed 30 of 33 HP1 molecules but only 6 of 33 HP2 molecules in the accessible state, matching the parental imprint. The biallelic *RPLP0* promoter, by contrast, looked alike on both haplotypes.

**Figure 4.**
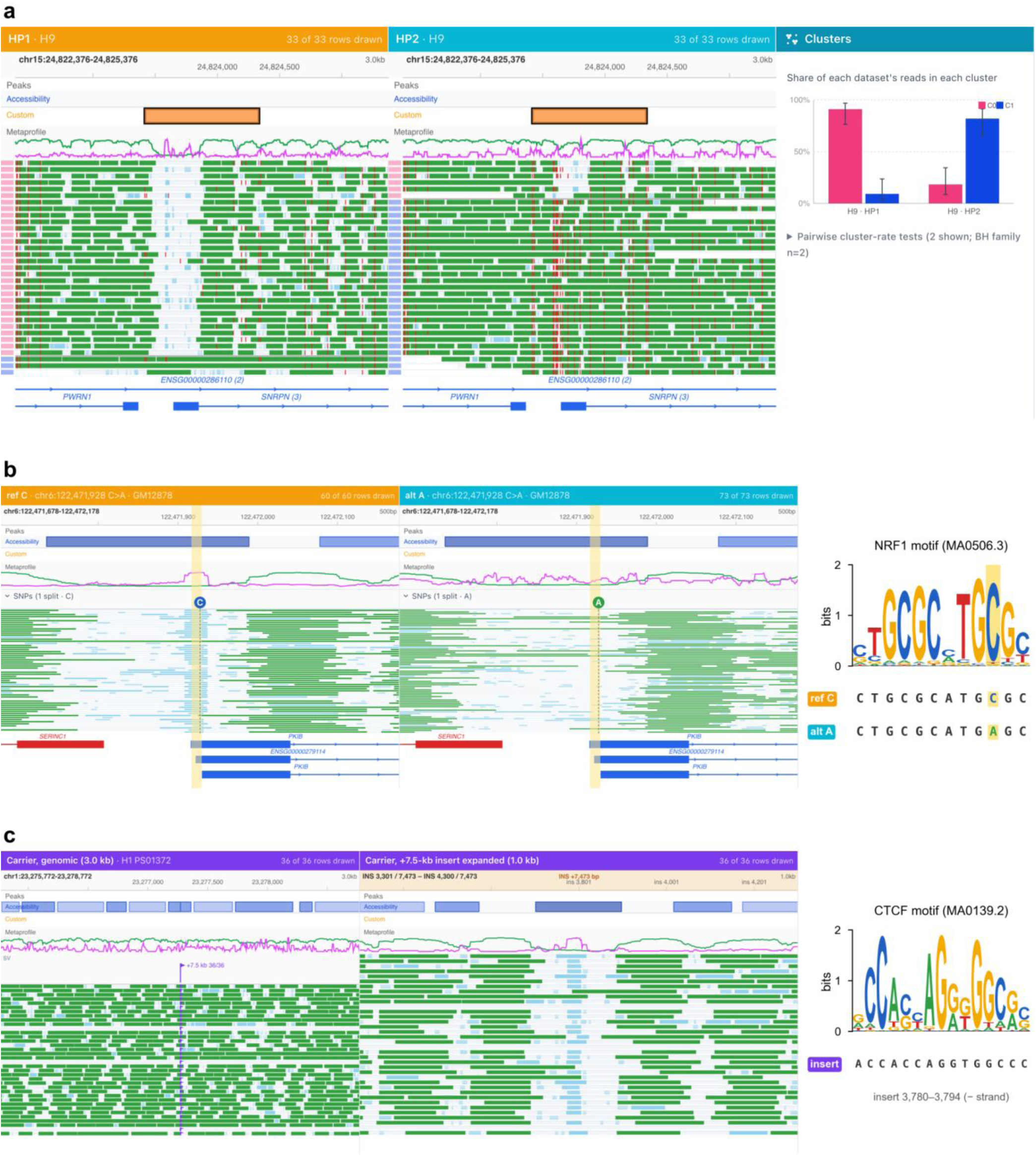
Dissecting allelic footprints, DNA methylation and non-reference structural variants on single fibers. a, Parent-of-origin chromatin at the imprinted *SNRPN* locus (chr15:24,822,376–24,825,376) in H9 Fiber-seq. Phased HP1 and HP2 molecules (33 each) are shown side by side with CpG methylation (red ticks, methylated CpGs) after joint k-means clustering (K = 2) over the 917-bp differentially methylated region (orange). Right, the interface’s per-haplotype cluster proportions; error bars, 95% Wilson intervals. Accessible cluster, 30/33 HP1 versus 6/33 HP2 molecules (two-sided Fisher’s exact P = 2.2 × 10⁻⁹; *RPLP0* in H1, P = 0.82). b, Allele-specific NRF1 occupancy at a heterozygous C>A variant at the bidirectional *SERINC1*–*PKIB* promoter (chr6:122,471,678–122,472,178) in GM12878 Fiber-seq. Molecules were split by the base at chr6:122,471,928 with the interface’s variant scan (reference C, 60 molecules spanning the motif, 59 with footprint calls; alternative A, 73 molecules; all drawn). The marker above each panel gives the variant base, the dashed line its position on the molecules, and yellow the NRF1 motif core. Right, the JASPAR NRF1 motif (MA0506.3) with the two alleles on the reference strand; the variant changes motif position 10 from C to A. Motif footprinted on 51/59 C and 4/73 A molecules (two-sided Fisher’s exact P = 4.0 × 10⁻²³ for the displayed molecules; top-ranked of 8,055 tests in the haplotype scan, 52/61 versus 4/76 molecules, Benjamini–Hochberg q = 1.5 × 10⁻¹⁹; Supplementary Table 8); replicated in both heterozygous H1 libraries (P = 8.2 × 10⁻¹¹ and 4.1 × 10⁻⁴). c, A non-reference insertion at chr1:23,277,272 in H1 Fiber-seq (7,471 bp in the reference alignment; 7,473-bp carrier consensus). Left, the 36 carrier molecules at the insertion site (3 kb; flag, insertion). Right, the same molecules with the insertion consensus expanded at insert positions 3,300–4,300 (1 kb), showing a nucleosome-free region at insert positions 3,723–3,879 with a footprint overlapping 3,771–3,804 on 28 of 36 carriers (78%; 26 covering the complete CTCF motif). The insert consensus partly matches MER41A and contains Alu fragments. Far right, the JASPAR CTCF motif (MA0139.2) with the inserted sequence at insert positions 3,780–3,794 (minus strand), inside the central footprint (FIMO P = 3.2 × 10⁻⁷, q = 0.005). Subnucleosomal footprints are sky blue on molecules; footprint metaprofiles are magenta.

Beyond the accessibility of whole elements, FiberBrowser resolves how single-nucleotide variants change individual binding events, even when promoter accessibility is unchanged. In GM12878, a heterozygous C>A variant falls in the core of an NRF1 motif at the bidirectional *SERINC1–PKIB* promoter. FiberBrowser’s variant scan finds the variant in the reads and splits the molecules by the base they carry (Fig. 4b). The promoter is open and largely unmethylated on both alleles, but only the reference allele is bound. A footprint covers the motif on 51 of 59 molecules carrying C and on 4 of 73 carrying A, while the rest of the promoter looks alike. The variant therefore abolishes occupancy at that NRF1 element without changing promoter accessibility or CpG methylation. Both H1 libraries heterozygous for the same variant show the same allelic footprint.

FiberBrowser also reveals chromatin inside structural variants that reference-based browsers cannot show. When molecules carry an insertion absent from the reference genome, standard browsers mark or clip the inserted sequence and hide its chromatin. FiberBrowser instead builds the inserted allele from the carrier molecules and draws their chromatin calls across it. In H1, a 7.5-kb insertion lies on one haplotype (Fig. 4c). On the reference genome, carrier molecules show regular nucleosome arrays on either side of the insertion. Expanding the inserted sequence reveals a 156-bp nucleosome-free region on most carrier molecules, with a footprint at its center on about 80% of them. The footprint contains a canonical CTCF motif, so the insertion brings a CTCF site, footprinted on most molecules, that analyses on the reference genome miss entirely.

### Footprint representations transfer cell-type classifiers across assays

Because FiberHMM translates diverse chemistries into a common representation of nucleosomes and footprints, models trained on one assay should also interpret another. We trained a convolutional neural network (CNN) on footprints and reference sequence in 5-kb promoter windows to distinguish five Hia5 Fiber-seq cell lines (GM12878, a lymphoblastoid cell line (LCL); H1; H9; K562; HepG2), combining promoter-level evidence into one prediction (Fig. 5a and Supplementary Fig. 10a–c). The CNN classified all 40 held-out Fiber-seq pseudo-cells correctly, each built from one or two molecules per promoter (Fig. 5b).

**Figure 5.**
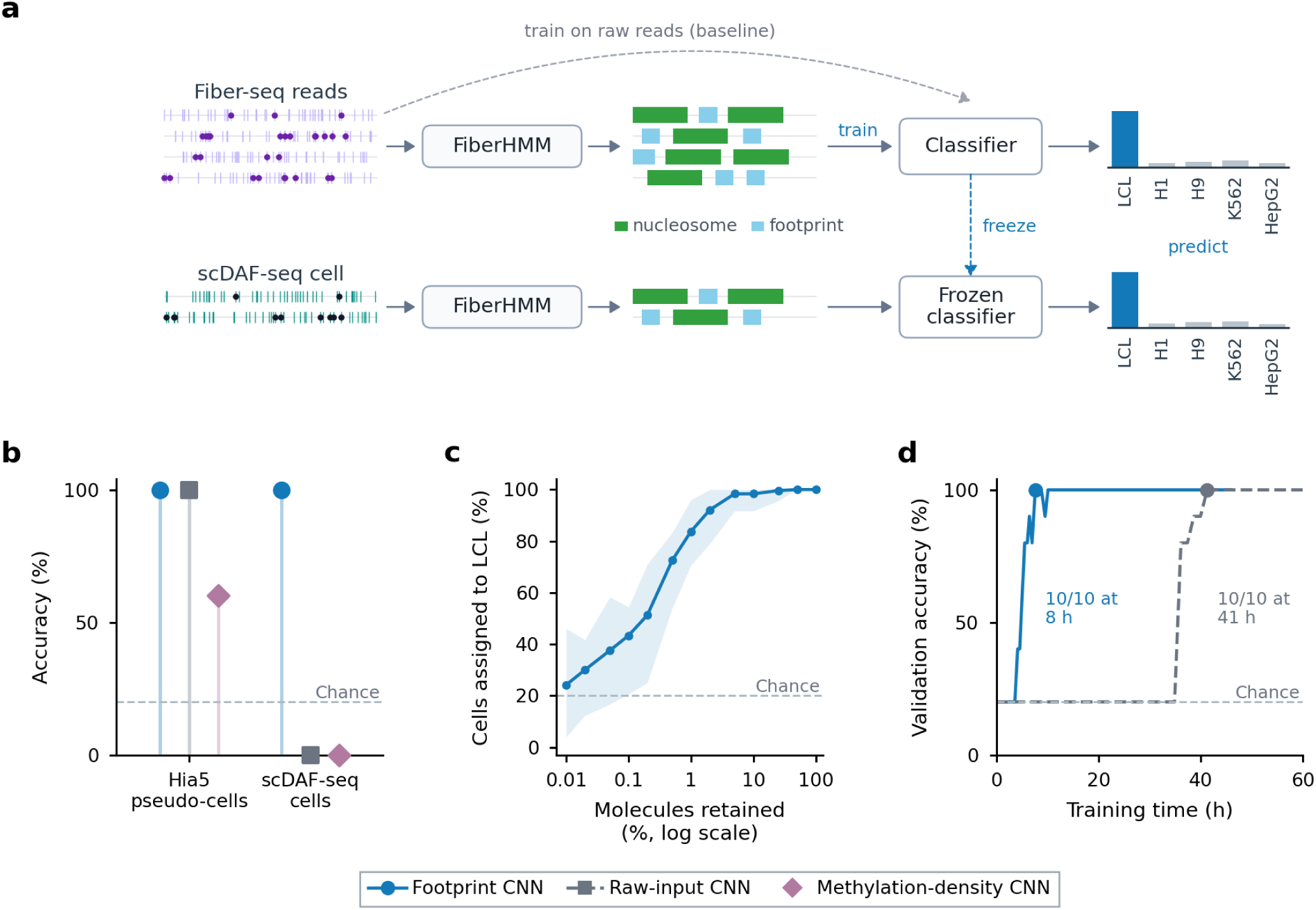
Footprint encoding supports cell identification across assays. a, Schematic. Fiber-seq reads are encoded by FiberHMM as nucleosome and footprint calls and used to train a promoter CNN that calls cell type (LCL, H1, H9, K562, HepG2); dashed arc, a CNN trained on raw reads instead. The trained classifier is frozen and applied to FiberHMM-encoded reads from single HG002 scDAF-seq cells; bar heights are illustrative. b, Full-input accuracy in Hia5 pseudo-cells (40/40 correct for the footprint and raw-input CNNs; 24/40 for the methylation-density CNN) and scDAF-seq cells (footprint CNN, 12/12; raw-input and methylation-density CNNs, 0/12). Blue, footprint CNN; gray, raw-input CNN; mauve, methylation-density CNN; dashed guide, five-class chance. c, LCL assignment across 13 molecule-retention levels (log-scaled axis) for the footprint CNN (240 predictions per level); shading, 2.5th–97.5th percentiles across footprint-CNN draws; dashed guide, five-class chance. d, Accuracy on the ten fixed Hia5 validation profiles against logged training and loading time; points mark the first evaluation at which each CNN classified all ten correctly (footprint CNN, 7.6 h; raw-input CNN, 41.3 h). Model architecture and training histories, Supplementary Fig. 10a–d; negative controls and the methylation-density CNN, Supplementary Fig. 11.

Applied without retraining to a different assay and enzyme, it identified all 12 HG002 scDAF-seq cells as LCL (Fig. 5b). Downsampling the scDAF-seq molecules showed how assignment depends on the number of molecules (Fig. 5c). Assignment stayed above 99% with 25% of molecules and above 50% with 0.2%. With one or two molecules per cell it fell to near chance (24%, versus 20%). Shuffling nucleosome and footprint calls within each molecule, assigning molecules to the wrong promoters, or supplying only the observation mask yielded LCL in 2 of 984 predictions (Supplementary Fig. 11a). The predictions therefore follow the positioned footprints in the molecules rather than coverage.

By contrast, a CNN with the same architecture trained on raw m6A patterns did not transfer and learned more slowly. It also classified all 40 Fiber-seq pseudo-cells correctly, but it assigned none of the scDAF-seq cells to LCL (Fig. 5b), and it first classified all ten held-out Hia5 validation profiles correctly after 41 h of training, compared with 8 h for the footprint CNN (Fig. 5d and Supplementary Fig. 10c,d). A CNN of the same architecture given the local methylation density instead of footprints classified 24 of the 40 pseudo-cells correctly and assigned none of the scDAF-seq cells to LCL (Fig. 5b and Supplementary Fig. 11b,c). Footprint inference therefore makes trained models portable across single-molecule assays.

## Discussion

Our study presents a unified framework for the analysis and interactive exploration of diverse single-molecule chromatin assays, FiberBrowser and its analytical engine, FiberHMM. Across distinct enzymes, chemistries and sequencing platforms, FiberHMM translates heterogeneous modification readouts into a common per-molecule representation of nucleosomes, accessible DNA and subnucleosomal footprints. FiberBrowser then allows thousands of molecules to be clustered, phased and compared in real time while retaining access to the observations underlying every call.

This approach preserves the resolution supported by each experiment. Because footprint evidence weighs observed modifications against the informative sites where the enzyme could act, FiberHMM distinguishes limited information from evidence of protection. Dense Hia5 PacBio Fiber-seq and duplex SsDddA scDAF-seq recover transcription factor footprints below 20 bp, whereas lower-resolution, single-stranded Hia5 ONT Fiber-seq is better suited to nucleosome architecture. Footprint discovery requires no motif annotations, factor identities or prespecified footprint widths, although assigning a protein to an inferred footprint still requires independent evidence such as motifs or ChIP. Existing tools serve complementary roles. Fibertools calls m6A and supports broad manipulation and analysis of Fiber-seq data, IDLI models nucleosomal substructures in SAMOSA, and HiddenFoot infers binding from supplied sequence and motif models. By contrast, FiberHMM returns per-molecule nucleosome and footprint calls for multiple chemistries in one BAM-native format, and its output can substitute into chemistry-specific pipelines, as shown for FIRE.

Combining this footprint caller with FiberBrowser reveals regulatory mechanisms that aggregate tracks and reference-bound browsers miss. Because long reads preserve genetic variants, CpG methylation and chromatin structure on the same molecules, splitting and clustering molecules links genotype directly to chromatin state. The examples here include the imprinted alleles at *SNRPN*, an NRF1 site lost from an open promoter through a single base change, and a CTCF site carried by a 7.5-kb non-reference insertion.

The transfer of cell-identity predictions from Fiber-seq to scDAF-seq shows that this shared representation retains information about chromatin state across chemistries. A classifier trained on Fiber-seq footprints recognizes lymphoblastoid identity in single-cell DAF-seq without retraining, whereas a model trained on raw modifications fails to transfer. Additional scDAF-seq cell types are needed to establish the specificity and generality of transfer. Deeply sampled reference maps in one assay could then help interpret sparse single-cell measurements from another, allowing assays with distinct advantages to contribute to a common analysis of chromatin state.

We designed FiberHMM and FiberBrowser to accommodate new single-molecule chemistries as they emerge. For conventional methyltransferases and deaminases, low-coverage treated naked-DNA and untreated controls can calibrate the observation model. SsDddA shows how rotational phase modeling, physical-duplex pairing and CpG island inference can extend the framework when naked-DNA controls cannot be amplified and enzyme activity depends on DNA methylation. Improvements in ONT m6A detection could be incorporated by updating the corresponding model parameters, and extension to EcoGII, the SMF methyltransferases and other enzymes awaits calibration and validation. FiberBrowser and FiberHMM are publicly available on GitHub, and we welcome collaborations to support additional single-molecule approaches.

## Online Methods

### Sample preparation and sequencing

#### Fly husbandry and embryo collection

Wild-type *yellow white* (*yw*) flies were maintained and embryos collected at 25 °C in petri-dish-sized collection cages. Pre-clearing collections were discarded earlier the same day, and timed collections were then aged to the required stage after egg laying^38^. The DddB amplicon libraries used half-hour collection windows from 1.5 to 4 h, with one pooled 2–4-h *ind* window; the whole-genome Hia5 libraries used embryos aged 2–4 h after egg laying. Embryos were dechorionated for 1 min 45 s in 50% bleach and rinsed^38^.

#### Nuclei isolation

Embryos were homogenized in 250 mM sucrose, 10 mM Tris–HCl (pH 8.0), 25 mM KCl, 5 mM MgCl₂ and 0.1% Triton X-100 with 40 strokes of a tight Dounce pestle. Nuclei were passed through a 40-µm filter, pelleted at 350*g* for 10 min at 4 °C and resuspended in 30 µl of 15 mM Tris–HCl (pH 8.0), 15 mM NaCl, 60 mM KCl, 1 mM EDTA, 0.5 mM EGTA and 0.5 mM spermidine. Each collection was reacted separately, and reactions were pooled before DNA purification or PCR^38^.

#### Hia5 Fiber-seq

Hia5 was purified and activity-calibrated as described^3^ and provided by the Stergachis laboratory. Nuclei were treated with 0.8 mM S-adenosylmethionine and 200 U Hia5 for 10 min at 25 °C, the reaction was stopped with SDS (1% final), and high-molecular-weight DNA was purified with the Wizard HMW DNA Extraction Kit (Promega, A2920)^38^.

#### PacBio Fiber-seq

The *Drosophila* 2–4-h library pooled independently collected and Hia5-treated embryo batches and has been described^38^. It was sequenced at the University of Washington on a PacBio Revio (SPRQ chemistry, instrument control software 13.3). HiFi reads with m6A calls were generated on the instrument with ccs 8.2.0 and jasmine 2.3.0, demultiplexed with lima 2.11.0 and aligned to dm6 with pbmm2 1.17.0. Reads overlapping the four amplicon loci were used here.

#### Hia5 ONT Fiber-seq and controls

A 2–4-h library pooling six independently collected and Hia5-treated embryo batches was prepared as above. For the accessible-state (naked-DNA) control, genomic DNA purified by the same procedure was treated with Hia5 under the standard reaction conditions and repurified. For the protected-state (untreated) control, nuclei were carried through the same workflow without Hia5. Libraries were prepared from unsheared DNA without size selection with the Ligation Sequencing Kit V14 (SQK-LSK114) and sequenced on R10.4.1 flow cells (FLO-PRO114M) on a PromethION 2 Solo. Reads were basecalled with dorado 2.0.1 (dna_r10.4.1_e8.2_400bps_sup@v5.2.0 with the 6mA@v1 modified-base model).

#### DddB footprinting

DddB was expressed and purified as described^38^ from a plasmid provided by X. S. Xie^14^. Nuclei were footprinted for 30 min at 25 °C in 50-µl reactions containing 2 µl DddB in DRB buffer (20 mM Tris–HCl pH 7.5, 20 mM NaCl, 2 mM DTT); all other steps matched the Fiber-seq workflow^38^.

#### Targeted amplicons

The *hb*, *ind*, *scyl* and *chrb* loci (9.5–17.8 kb) were amplified from the same DddB-treated embryo DNA with locus-specific primers (Supplementary Table 9) using KeyPo Master Mix (Vazyme), and sequenced by Plasmidsaurus with their standard Oxford Nanopore amplicon service (R10.4.1 flow cells, super-accuracy basecalling).

#### DddB naked-DNA controls

Genomic DNA purified from *yw* embryos as above was deaminated with DddB under the standard footprinting conditions and repurified. This DNA was then whole-genome amplified with the REPLI-g Mini Kit (Qiagen). The unamplified control was untreated genomic DNA, sequenced without deamination or amplification. Both libraries were sequenced by Plasmidsaurus on R10.4.1 flow cells and basecalled with dna_r10.4.1_e8.2_400bps_sup@v4.3.0.

#### Human cell lines

Hia5 Fiber-seq of H1, H9, K562 and HepG2 was performed as described^21^. H1 and H9 were obtained from the University of Washington Institute for Stem Cell and Regenerative Medicine, and K562 (CCL-243) and HepG2 (HB-8065) from ATCC. Libraries were sequenced on PacBio Revio (SPRQ chemistry), m6A was called on the instrument with jasmine 2.7.99, nucleosomes were annotated with fibertools 0.8.0 (ref. 20), and reads were aligned to hg38 with pbmm2 1.17.0.

### Software implementation and supported assays

FiberHMM 3.1, the analytical engine of FiberBrowser, provides command-line workflows for calling and duplex reconstruction. The FiberHMM command line and the FiberBrowser interface invoke the same engine, so interactive use and batch analysis share the fitted model and numerical implementation. Standard BAM outputs retain molecule identities, raw observations, calls, derived annotations and provenance.

FiberHMM is implemented in Python and processes aligned long-read BAM files^39^ through the command-line entry point fiberhmm-call. SsDddA, DddB and Hia5 are selected with assay presets, and Hia5 input is additionally designated as PacBio or ONT. Coordinate-sorted, indexed BAMs are processed in genomic chunks and merged without changing read names or source alignments; unindexed input can be streamed. Secondary, supplementary and unmapped records were excluded from inference in the reported analyses. Unless stated otherwise, footprint calls used FiberHMM 3.0.4–3.1.0; these releases share the HMM, refinement and footprint-calling algorithms and differ in the packaged DddB emission table (the refit described below, used for all DddB calls), the read-length filter for soft-clipped alignments and added commands such as fiberhmm-footprint. The interface analyses used FiberBrowser 3.0.0; the exact source commit and emission-table hash for each analysis are recorded in the accompanying provenance manifests.

Performance was measured on a desktop workstation. On a 4-Mb H1 Hia5 PacBio BAM containing 19,540 primary records, FiberHMM completed calling in 163 s with 218 MiB peak memory using eight compute threads. Browser display, navigation and analysis benchmarks are described in Supplementary Note 2 and Supplementary Fig. 9a,b.

### Molecular Annotation encoding and FIRE interoperability

FiberHMM writes, and the FiberBrowser interface reads, the Molecular Annotation specification developed by Mitchell R. Vollger (https://github.com/fiberseq/Molecular-annotation-spec). The MA:Z tag stores the original read length and 1-based start-length intervals in original molecule orientation, together with annotation type, biological strand and zero or more typed quality declarations. Q declares a linear 0–255 byte and P a Phred-scaled byte. AQ:B:C stores the declared bytes in MA interval order, and AN:Z stores optional names aligned to all MA intervals. Reference coordinates are derived through the alignment and do not replace molecule coordinates.

FiberHMM layers use nuc.Q or nuc.QQQ for nucleosome quality and optional edge qualities, msp. for accessible-patch coordinates and tf.QQQ for footprint TQ and left/right edge qualities, where TQ = min(255, round(10 × LLR)). Final MSP coordinates are also mirrored in the fibertools as/al arrays so that existing fibertools and FIRE versions can consume the same segmentation.

For FIRE interoperability, fibertools-rs 0.13.0 ft fire was run separately on fibertools and FiberHMM segmentations of the same 10,000-read GM12878 PacBio subset; 9,781 reads carried FIRE annotations in both outputs, and final FiberHMM MSP calls matched the as/al arrays consumed by FIRE. Positive FIRE calls followed the documented aq > 230 definition, and a directional call was shared when it overlapped a positive call from the other segmentation by at least 1 bp on the same read. At aq > 230, approximately 81% of fibertools calls and 79% of FiberHMM calls were shared. The concordance curve retained calls at each cumulative minimum aq; at aq ≥ 249 (aq/255 ≥ 0.976 estimated precision) the directional fractions were 98.0% and 94.8% (95% molecule-bootstrap intervals 97.6–98.3% and 94.3–95.4%). The 17,340 shared overlap components containing exactly one call from each segmentation had a median absolute width difference of 8 bp (interquartile range 4–21 bp; Supplementary Fig. 1e,f).

### Molecular observations and informative sites

FiberHMM represents a molecule by its mapped reference blocks, the sequence positions at which the selected enzyme could generate an observable event (informative sites) and the subset of those positions carrying an event. SsDddA and DddB informative sites were reconstructed on the relevant cytosine strand after alignment to the reference. Hia5 informative sites were adenines or thymines in the reference frame, with platform-specific modification tags supplying observed m6A. Aligned deletions and uncovered bases were treated as missing observations, not as evidence of protection.

Emission tables specify the probability of an event and of no event in the accessible and protected states for each encoded sequence context. Hia5 (PacBio and ONT) emissions and DddB protected-state emissions were derived from assay-matched controls; DddB accessible-state emissions were recalibrated in vivo, as described below. Hia5 PacBio emissions were counted in seven-base contexts from Hia5-treated and untreated *Drosophila* S2 genomic DNA, with m6A taken from the PacBio modification calls at ML ≥ 128; initial-state and transition probabilities are those of the original FiberHMM Hia5 PacBio model^33^.

Fully accessible naked-DNA SsDddA controls accumulate too many uridines to be amplified and sequenced efficiently, whereas nucleosomes and bound proteins keep conversion in chromatin within the recoverable range. The SsDddA nucleosome and footprint components therefore use separate tables: the nucleosome model accounts for internal rotational deamination, and the subnucleosomal table was derived from reconstructed physical duplexes as described below.

Adjacent same-strand SsDddA targets (CC on C-to-T reads, GG on G-to-A reads) do not convert independently. At confidently accessible *NAPA* sites, the second target of an isolated adjacent pair converted in 93% (CT) and 93% (GA) of molecules when the first did, and in 3% and 5% when it did not; targets 2–6 bp apart were not positively coupled (Supplementary Fig. 5f,g). Because the emission model treats each target as an independent observation, such a run would count one event more than once. FiberHMM therefore keeps only the 5′-most target of each run of adjacent same-strand SsDddA targets by default, on every informative-site set it evaluates: calling, footprint recall and physical-duplex merging (keep-one thinning,--daf-mask-runs; DddB informative-site sets are not thinned). This raised the precision of footprint calls read on the complementary strand of HG002 physical duplexes (Supplementary Fig. 5h).

### DddB emission calibration

DddB ONT emissions were first estimated from dm6-aligned *Drosophila* naked-DNA controls: a whole-genome-amplified (WGA), DddB-treated control and an unamplified, untreated control library. For each primary alignment, combined conversion was defined as (C→T + G→A)/(reference C + reference G). Analyses required mapping quality ≥20 and an aligned span ≥1 kb. The WGA control contained a prominent under-deaminated population. For a read with k conversions among n combined C/G informative sites, combined conversion was estimated as the Jeffreys-smoothed quantity (2k + 1)/(2n + 2). A two-component Gaussian mixture fitted after logit transformation to 395,658 eligible WGA molecules placed the equal-posterior boundary at 5.3% combined conversion (4.9–5.8% for unsmoothed logit- and log-rate fits), and WGA molecules with ≥20% combined conversion were retained as the accessible control.

For the unamplified control, the corresponding mixture fitted to all 564,675 eligible molecules separated a low-conversion component centered at 0.82% from a higher-mismatch component centered at 5.4%. Molecules with <1.75% combined conversion, below the 2.3% equal-posterior boundary (posterior probability of the low-conversion component 0.66 at 1.75%), were retained (49.0% of eligible molecules) as the protected-state background.

Up to 100,000 passing reads per control were used for emission estimation, excluding 10 bases from each read end. C→T observations were counted in C-centered seven-base contexts, and G→A observations were reverse-complemented into the same context space. The fixed central C leaves six variable flanking bases, giving 4,096 contexts. These control-derived probabilities replaced only the emission matrix; initial-state and transition probabilities were copied unchanged from the Hia5 PacBio model.

Comparison with in vivo data indicated that the naked-DNA table underestimates the accessible-state deamination rate in chromatin. The accessible-state emissions were therefore recalibrated to an in vivo level set by matched Hia5 PacBio Fiber-seq, keeping the sequence-context dependence of the naked-DNA table.

At four *Drosophila* amplicons (*chrb*, *hb*, *scyl* and *ind*), the overall accessible-state level was set from the modification burden of matched wild-type 2–4-h Hia5 PacBio Fiber-seq (11 SMRT Cell movies; m6A calls in these data all carry ML ≥ 250, so the result does not depend on the ML threshold). For each amplicon, Hia5 molecules (primary alignments with mapping quality ≥20 covering at least 0.5 kb of the amplicon interior, excluding 100 bp at each end) gave an implied accessible fraction f = (m − E_prot_)/(E_acc_ − E_prot_), where m is the number of m6A calls and E_acc_ and E_prot_ are the numbers expected if every informative site were accessible or protected under the Hia5 PacBio table. DddB deaminations observed over the same intervals in the 22 amplicon libraries (wild-type 1.5–4-h embryos; PCR duplicates collapsed) were divided by the number expected from the naked-DNA table at accessible fraction f. This ratio was 0.95 (hb), 1.21 (scyl), 1.28 (chrb) and 1.48 (ind), and the accessible-state level was set to the median, 1.245 times the naked-DNA table (95% bootstrap interval 1.23–1.26, resampling molecules within amplicons). An independent per-context fit, estimating each enzyme’s accessible-state rate against accessibility measured on a separate half of its own molecules, gave a consistent Hia5-calibrated level of 1.24-fold (95% locus-bootstrap interval 1.14–1.28; Supplementary Fig. 1a). This fit used five-base contexts at the *hb*, *ind*, *chrb* and *scyl* amplicons (DddB: the 22 amplicon libraries, wild-type 1.5–4-h embryos, called with the naked-DNA table; Hia5: PacBio 2–4 h, 11 movies), modeled each context as P(event) = a·p_acc_ + (1 − a)·p_prot_ with a the fraction of molecules from the other half whose MSP covers the site outside called footprints, and gave protected-state rates of 0.0046 (DddB) and approximately 0 (Hia5); excluding the 3 DddB contexts (1.1% of sites) fitted at the upper bound gave 1.23-fold. Deamination inside called nucleosomes was low and showed no rotational periodicity, so protected-state emissions were left unchanged.

The resulting accessible-state emissions are logit p_acc_(c) = logit p_acc,naked_(c) + α, with α = 0.488 chosen so that the informative-site-weighted mean accessible-state probability across the DddB amplicons is 1.245 times the naked-DNA value (0.409 to 0.509), and capped at 0.95 (1 of 4,096 contexts). Per-context values changed 1.02-to 1.58-fold (median 1.29). Initial-state and transition probabilities were unchanged. This in vivo-calibrated table is the default DddB model.

### Hia5 ONT emission calibration

Hia5 ONT emissions were estimated from two control libraries of *Drosophila yw* 2–4-h embryos described above: naked DNA treated with Hia5 for the accessible state and untreated (mock-treated) nuclei for the protected state. Reads were aligned to dm6 with minimap2^40^ (map-ont). Primary alignments with mapping quality ≥20 and an aligned span ≥1 kb were used, and m6A calls required ML ≥248, the threshold used for Hia5 ONT calling.

The naked-DNA control contained a low-methylation population. A two-component Gaussian mixture fitted to the logit-transformed, Jeffreys-smoothed per-read fraction of basecalled adenines called m6A placed the boundary for ≥0.95 posterior assignment to the methylated component at 11.4%, and the 49,961 of 62,033 eligible reads above it were retained. All 842,434 eligible untreated reads were used. Events and non-events were counted in adenine-centered seven-base contexts in the sequenced-strand orientation, excluding 10 bases at each read end (4,096 contexts per state; 1.97 × 10⁹ protected-state positions). Initial-state and transition probabilities were copied from the Hia5 PacBio model.

### First-pass hidden Markov segmentation

The first stage is the two-state FiberHMM introduced previously^33^. The states represent accessible and protected sequence, and emissions are selected by assay and local sequence context. The model is run along the query bases of each molecule, with emissions for informative and non-target positions as in the original model, and calls are projected to reference coordinates through the alignment; the refinement steps below score informative sites only. Contiguous accessible segments define modification-sensitive patches (MSPs), and protected segments provide the initial nucleosome-scale topology. The generalized method retains the published HMM parameters and topology and adds observation presets, post-HMM refinement and a common output format.

### Nucleosome and subnucleosomal-footprint likelihood refinement

The subnucleosomal step is referred to as footprint refinement.

Post-HMM inference is divided into nucleosome and subnucleosomal-footprint tasks. For a candidate segment with *observations x_i_ and sequence contexts c_i_*, FiberHMM accumulates the evidence

LLR = Σ_i_ log[P(x_i_ | protected, c_i_) / P(x_i_ | accessible, c_i_)].

For Hia5 and DddB, the two refiners use this context-specific additive score in opposite directions. Within each HMM-protected segment, the sign-inverted score identifies local runs of accessible evidence that support linker cuts; the protected-state score is then applied to each remaining fragment to define conservative nucleosome edges and a call quality score. The resulting accessible sequence is merged with the original HMM-accessible state to rederive MSPs. The footprint refiner then selects disjoint segments within the scan domains (MSPs and short HMM-protected candidates) that maximize the sum of segment LLRs minus a fixed penalty per selected segment, subject to a minimum informative-site requirement; this multi-segment selection is exact. SsDddA replaces the accessible-cut step with the rotational refinement described below.

The default footprint threshold is LLR 5 for SsDddA, DddB, Hia5 PacBio and Hia5 ONT, with at least three informative sites. Each call stores TQ = min(255, round(10 × LLR)), so the default TQ floor is 50 in every preset. TQ can be used for downstream filtering, but changing the per-call penalty requires fresh decoding because it can split, join or newly detect calls. TQ is an evidence score, not a posterior binding probability or false-discovery rate.

For the threshold benchmark in Supplementary Fig. 2a,b,d, 320 motif-centered genomic contexts were selected, the motif-forward and reverse-complement informative-site sets were evaluated independently, and protected spans of 8, 10, 15, 20, 25, 30, 40, 50 and 60 bp were programmed. Sixteen protected draws and sixteen matched fully accessible controls were generated for every context, orientation and size, giving 92,160 protected and 92,160 accessible-control trials per chemistry and threshold. No feasibility or minimum-information filter was applied before evaluation. Each threshold from LLR 2 through 8 was rerun, and a trial was recovered when a returned subnucleosomal call covered the programmed center.

SsDddA used the HMM, rotational nucleosome refiner and footprint refiner without keep-one thinning, because simulated targets are drawn independently, with the two strand-specific informative-site sets called independently; the other assays used their default calling settings. The endpoint measures center-covering footprint recovery rather than boundary accuracy. Uncertainty bands are 95% intervals from resampling genomic context blocks. The selected common setting, LLR 5, is the first integer threshold at which simulated accessible-control calls were below 2% in all four chemistries. Empirical false calls were measured in the Hia5 PacBio naked-DNA control library (*Drosophila* S2 genomic DNA treated with Hia5), the library used to fit the accessible-state emissions. The 46,678 molecules used for emission fitting were excluded, and 20,000 of the remaining primary alignments (MAPQ ≥20, aligned length ≥1 kb) were sampled with a fixed seed and called with the default Hia5 PacBio settings at LLR 5. One 800-bp window was fixed per molecule, and a false call was scored when a footprint covered the window center, the endpoint used for the simulated accessible controls. The model expectation was obtained by drawing observations from the accessible-state emissions at each molecule’s own informative sites and calling them in the same way (mean of three repeats). Per-molecule labeling was the fraction of informative A/T sites called as m6A. To separate variation in labeling between molecules from effects within molecules, the simulation was repeated with each molecule’s accessible-state probabilities scaled to its observed labeling, which gave 1.5% (95% CI 1.4–1.5%). Real calls averaged 2.9 footprints per kb of query sequence, 69% of them shorter than 15 bp. Intervals are percentiles of 5,000 molecule bootstrap resamples (Supplementary Fig. 3a). The same analysis was applied to the DddB and Hia5 ONT naked-DNA controls (Supplementary Fig. 3b). For DddB, 20,000 eligible WGA molecules with ≥20% combined conversion were sampled with the same seed and called with the default DddB settings at LLR 5, with simulated deaminations placed at informative sites on each molecule’s deaminated strand. For Hia5 ONT, the naked-DNA control was aligned to dm6 with minimap2 (map-ont), and 20,000 eligible molecules above the 11.4% boundary were sampled and called with the default Hia5 ONT settings. Per-molecule labeling was the fraction of informative sites called deaminated or m6A, and calls were scored, simulated and bootstrapped as for Hia5 PacBio. Unlike the Hia5 PacBio molecules, these molecules overlap the emission-fitting data, because every molecule passing these filters was eligible for fitting.

### Rotational SsDddA nucleosome refinement

SsDddA nucleosome refinement begins from permissive HMM-protected cores. Candidate extensions are evaluated against a phase model, distributed with FiberHMM, describing the probability of internal deamination as a function of distance and rotational phase from the protected core. The model allows a periodic series to skip one or more expected events and permits modest variation in spacing. A proposed edge is supported when accessible evidence in the flank is inconsistent with continuation of the periodic series; boundary uncertainty is retained when competing extensions have similar support.

The radial deamination profile of the phase model was estimated from pooled *NAPA* and *UBA1* amplicon molecules and applied unchanged to the genome-wide scDAF-seq libraries, which were not used for fitting; *NAPA* and *UBA1* diagnostics therefore overlap the fitting data. The molecule-matched ablation disabled only the phase model, with code, input molecules and all other parameters identical (Supplementary Fig. 5a,b). Internal-event counts, sidedness and caller-independent recurrent-site maps were evaluated as separate diagnostics (Supplementary Fig. 5c–e). DddB nucleosomes showed no detectable rotational phase, and the standard likelihood refiner remains the default.

Nucleosome density in the phase-model ablation was calculated by dividing the number of nucleosome and footprint calls by the sum of CIGAR-aligned bases over the same primary alignments. Length categories were defined before counting as less than 120 bp, 120–200 bp, 201–249 bp and at least 250 bp. The two call sets contained identical molecule names and 64,048,017 mapped bases.

The molecule-matched ablation included 20,935 scDAF-seq molecules. In it, 120–200-bp calls increased from 1.07 to 3.67 per mapped kb, calls of at least 250 bp became uncommon, and total calls increased from 2.78 to 4.79 per mapped kb. Event sidedness was evaluated against an equal-side finite-count null rather than treating asymmetry among sparsely modified particles as evidence of a preferred side.

### SsDddA event burden at recurrent nucleosome sites

Recurrent nucleosome dyads at *NAPA* and *UBA1* were nominated independently in two deterministic molecule halves. Peaks retained in both halves defined fixed 120-bp windows, and each molecule was evaluated only at centers learned from the opposite half. Every primary alignment spanning at least 95% of a window contributed its observed SsDddA events and non-CpG informative sites. Sequence- and radial informative-site weights were fitted separately by conversion strand in the opposite molecule half, and each fiber’s total internal-event count was allocated among covered positions in proportion to these weights. Burden ratios were calculated in the pooled data and separately for CT and GA channels; the figure reports the pooled estimate with molecule-bootstrap 95% intervals. Colors indicate enrichment or depletion at family-wise corrected P < 0.05, with nonsignificant positions gray. A secondary analysis restricted the calculation to windows containing a rotationally refined nucleosome call within 75 bp of the cross-fitted center.

### Endogenous call-length distributions

Figure 1c uses all primary mapped bases and all final nucleosome and footprint calls from 22 passing DddB libraries spanning the *hb*, *ind*, *scyl* and *chrb Drosophila* amplicons after FiberHMM calling (HMM, nucleosome and footprint refinement) at LLR 5 (86,591 molecules; 542,152,784 aligned bp; 225,556 footprint calls; 762,478 nucleosome calls). The Hia5 PacBio distribution uses 3,000 K562 molecules and the Hia5 ONT distribution uses 11,162 *Drosophila* yw embryo molecules (2–4 h; four 50-kb windows), both at the same LLR 5 footprint threshold. Curves report called base pairs per 100 kb aligned per one-base length bin and are smoothed for display only (Gaussian kernel, s.d. 2 bp). HMM-only distributions were obtained before nucleosome and footprint refinement on the same DddB, Hia5 ONT and Hia5 PacBio molecules, with the original SNP masks and read filters; skipped molecules remained in the aligned-base denominator and contributed no calls.

### Real-molecule stage illustrations

For the Fig. 1 stage illustrations, raw observations were passed through the FiberHMM HMM and refinement steps at LLR 5, and intermediate nucleosome boundaries were captured immediately before footprint *refinement.* Candidate windows were screened in 160 *hb* DddB molecules, 1,500 *Drosophila* embryo Hia5 ONT molecules and 160 K562 PacBio molecules, and three 600-bp windows were selected to illustrate a nucleosome split or boundary change and a supported subnucleosomal call. Each stage shows the same informative sites and observations in read coordinates. All three assays use the same refinement settings. Read identifiers, windows, calls and model hashes accompany the plotting data.

### Motif footprint scoring

Motif occupancy was scored with fiberhmm-footprint (FiberHMM 3.1.0), which reads FiberHMM calls from a BAM file or from fiberhmm-extract bigBed files and motif intervals from a BED or bigBed file. A molecule is eligible for a motif when an MSP spans the complete motif, and bound when one footprint call (LLR ≥5) contains the motif within its outer edges, no m6A (ML ≥125) falls within the motif and the conservative call width lies within [--min-width,--max-width]. Outputs are a table in the ft footprint format and optional per-motif MA layers for FiberBrowser. The default window (0–74 bp) is label-free: 74 bp is the 95th percentile of conservative widths of footprint calls overlapping the benchmark motifs on MSP-eligible molecules. fiberhmm-footprint calibrate learns a window from ChIP-seq labels by grid search over minimum and maximum widths in 5-bp steps, maximizing the mean per-cell-line AUROC; with--holdout cell it reports leave-one-cell-line-out AUROC, together with the windows within 0.005 AUROC of the optimum and a settings file that applies the window to other data. For the benchmark, both modes were run with--msp-bam set to the BAM read by ft footprint, so scoring, calibration and ft footprint share the same eligible molecules. Motif panels, labels and statistics are described in Supplementary Note 1.

### DAF duplicate handling and variant screening

Amplified DAF reads were grouped into same-strand amplification families using alignment endpoints and deamination fingerprints. Clustering was constrained to compatible endpoints and required a minimum number of informative events; reads that could not be fingerprinted remained labeled as such. The default caller retained all source reads; population analyses used one deterministic representative per family. Potential opposite-conversion variants were screened only when recurrent evidence was observed in both directions; screening was disabled automatically below its coverage requirement (defaults: alternative fraction ≥0.20, depth ≥5 and at least five supporting fibers), and masked sites were removed from the observations without refitting emission or transition parameters. Amplification families required endpoints within 50 bp, deamination-fingerprint Jaccard similarity ≥0.95 and at least ten deaminations.

### Reconstruction of physical scDAF-seq duplexes

Duplexes were reconstructed independently within each scDAF-seq library after collapsing same-strand amplification families (see DAF duplicate handling and variant screening). Reads were classified by their dominant SsDddA conversion channel as CT or GA, and candidate mates were restricted to overlapping reads from opposite channels. Each strand was called independently before pairing. Reference-coordinate nucleosome dyads from the broad call were placed on a 10-bp grid and convolved with a 30-bp Gaussian kernel, producing a nucleosome-scale fingerprint that retains nucleosome order and approximate position while absorbing strand-specific boundary variation.

Candidate pairs shared at least 1.5 kb of genomic span and contained at least four nucleosomes from each strand within that span. Similarity was the maximum normalized cross-correlation between the two nucleosome-density signals across register shifts of up to 60 bp. An assignment required reciprocal selection, correlation above the empirical unrelated-pair floor and a sufficient advantage over the next candidate in both directions.

Haplotype and deamination-insensitive sequence provided independent constraints. On haplotype-resolved alignments, candidates were restricted to the same maternal or paternal contig. Reads were also compared at shared reference A/T positions, which SsDddA does not alter: a strong reciprocal sequence preference could resolve a pair directly, and gross sequence discordance vetoed a nucleosome-pattern match. Local two-by-two ambiguities were resolved when one diagonal was uniquely supported; unresolved components remained explicitly unpaired. The pairing route, partner, correlation, sequence difference and assignment margin were recorded as BAM tags.

Accepted strands were projected into a common forward reference frame, with CT and GA events, informative-site sets and aligned coverage kept as separate channels; deletions and uncovered positions remained missing observations. Pair identities used for SsDddA calibration were fixed, using only the broad nucleosome architecture, before any query window was nominated or any model fitted.

The standalone fiberhmm-pair tool scores candidates with a fixed footprint-based score, combined by default with deamination-insensitive sequence evidence (hybrid mode); the genome-wide validation below used sequence-free scoring, and the calibration set above used sequence-assisted reconstruction. Candidates require at least 1,500 bp of overlap and four nucleosome dyads on each read. A fixed linear score combines lag-tolerant dyad correlation, dyad-anchor agreement, overlap and dyad-count features, and 20-bp-binned non-CpG protection correlations, including a residual relative to the local overlap-component background. A/T agreement, haplotype and footprint LLRs do not enter this score; the reference is used to enumerate enzyme informative sites. A selected edge must be reciprocal-best and exceed each competing edge, or a zero-score null, by a default margin of 1.0; other candidates remain unresolved. BAM provenance selects the calibration appropriate to the nucleosome-calling model recorded in the file. Pair scores, margins, partner identities and model identifiers are retained in BAM tags.

For the genome-wide pairing validation (Supplementary Fig. 6a–c), the fixed footprint score was evaluated across all primary chromosomes in twelve libraries. Connected components overlapping the chromosome 1 pairing-model fitting interval were excluded, and the prespecified chromosome 1 validation interval was retained separately. A/T features were withheld from footprint scoring, and default thresholds were not retuned. Independent sequence relationships classified selected pairs as matching the sequence-supported mate, selecting a different mate, or unresolved. Agreement used only sequence-assessable selections; unresolved relationships were neither successes nor errors. Recovery used all independently sequence-selected pairs, including footprint-unscorable pairs. Confidence intervals used 1,000 Poisson bootstrap resamples of connected components, preserving competing edges. Among the 511,214 selected test pairs, 503,763 (98.54%) had no decisive independent A/T mate assignment, and 76.97% had no homolog comparison. A sequence label was assigned only when both reads had a competing candidate. Where the two homologs are identical in sequence, such a label cannot distinguish the mate from the other homolog’s strand, so agreement was evaluated at loci where both reads mapped uniquely to one HG002 haplotype (diploid remapping): 5,297 of 5,516 selections agreed (96.0%; 95% interval 95.5–96.5%). Same-strand amplification copies were absent from the consensus reads (none of 10.4 million pairing-eligible reads in copy clusters). These counts describe the availability of independent validation assignments; pairing itself used the fixed nucleosome and non-CpG protection score with sequence and haplotype withheld.

### Joint inference from both strands of a physical duplex

For joint inference, fiberhmm-pair (merge and recall stages) combines an accepted physical pair in forward-reference coordinates while retaining the CT and GA channels and their aligned coverage. Missing flanks and deletions remain unobserved. The HMM, rotational SsDddA nucleosome refiner and footprint refiner run once on the combined molecule using the two-channel observation encoder. The merged record retains both source identities and strand-specific deamination and coverage annotations. Original single-strand calls remain in the separate source BAMs.

Population analyses count a successfully merged duplex once and retain both source identities. Tagged but unmerged pairs are rejected rather than counted as independent molecules. Merge failures retain the unchanged source records by default; an explicit pairs-only export restricts output to successfully merged molecules.

### Physical-duplex calibration and validation of SsDddA footprint emissions

Calibration used physical duplexes from the first 40 Mb of chromosome 1 in each of the 12 scDAF-seq libraries (23,388 duplexes), excluding CpG sites and the terminal 10 bp of each read. One strand was designated the query and the other, its partner strand (mate), was used only for evaluation. Query-accessible observations were selected inside the first-pass accessible state from local neighborhoods containing at least ten non-CpG informative sites with complete query deamination. Query-protected windows were maximal runs of at least nine consecutive missed non-CpG informative sites, bracketed by events and spanning no more than 40 bp; the first and last query misses fixed the window. Mate observations neither created a candidate nor moved its edges. Both CT-to-GA and GA-to-CT directions were analyzed.

Accessible- and protected-state mate outcomes were fitted separately with additive logistic models using the six flanking bases, with model structure and regularization selected by leave-one-library-out, state-balanced log loss (ridge penalty 1 in the validation fits below). The resulting context-specific probabilities were serialized into the shared FiberHMM model structure and used only by the footprint LLR; the first-pass SsDddA HMM and phase-aware nucleosome model were unchanged.

For the held-out partner-strand validation reported in Supplementary Fig. 2c, 541,821 accepted duplexes across twelve libraries were evaluated, yielding 386,864 query-nominated windows. Eight libraries supplied emission fitting, two were reserved for validation and two for testing; training-library diagnostics excluded the scored library from fitting. Protected and accessible query nominations were fixed before evaluating the mate, with at least three mate informative sites and CpGs excluded. The default footprint decoder was rerun at 14 penalties from 0 to 20, rather than filtering scores from a different penalty or using only a maximum fixed-window score. Standard emissions were compared with exact six-base emissions fitted on training libraries. Because the standard table was fitted on chr1:1–40 Mb in all twelve libraries, test duplexes overlapping this region were excluded, so that both models are evaluated on data neither was fitted to; this left 79,543 protected and 34,241 accessible test windows on 42,650 duplexes. Confidence intervals used 1,000 Poisson bootstrap resamples clustered by physical duplex, preserving repeated windows and both directions. At LLR 5, standard and expanded emissions gave protected-window recovery of 91.54% (95% CI 91.35–91.72%) and 93.37% (93.21–93.54%), respectively, and accessible-window call fractions of 4.17% (3.95–4.39%) and 4.31% (4.09–4.53%). Query labels are operational controls, not ground truth for factor binding.

### Alternative SsDddA accessible-rate estimation using Hia5 MSPs

An independent sensitivity analysis used Hia5 accessible segments to nominate SsDddA measurement regions at *NAPA* and *UBA1* (Supplementary Fig. 1c). Hia5 coverage was at least ten, MSP consistency at least 0.7, and recurrent footprint protection was excluded at recurrence 0.05 with at least three supporting molecules. Additional SsDddA protected-run masks required two, three or four consecutive misses, with maximum gap between informative sites 12 bp and span 90 bp. The horizontal reference is the independently reported partner-strand estimate of 89%, rounded as in Fig. 2e.

The genome-wide check processed all 6,022 nonoverlapping 1-Mb cores across 46 HG002 diploid contigs and all twelve scDAF-seq libraries, with Hia5 coverage at least five and MSP support at least 80% (Supplementary Fig. 1d). Recurrent Hia5 footprints were excluded. Both haplotypes of each chromosome remained in the same training, validation or test partition.

Each focal SsDddA observation was withheld when selecting accessible neighborhoods: at least six other C/G informative sites within 50 bp had to meet the indicated event-fraction threshold. Confidence intervals resample chromosome groups 2,000 times, retaining both haplotypes together. Context models were fitted on training chromosomes and selected using validation chromosomes; exact six-base models were selected for both strands under the independent Hia5 MSP mask.

Hia5-MSP-based estimates test how SsDddA deamination rates depend on the choice of measurement region (Supplementary Fig. 1c,d). At *NAPA*, masking runs of at least two, three or four missed deaminations gave rates of 90.5%, 84.2% and 80.6%; at *UBA1*, the corresponding rates were 94.0%, 88.4% and 82.6%. Because SsDddA outcomes contribute to the mask, these are selection-dependent rates rather than unbiased conversion probabilities; the independent partner-strand estimate is approximately 89%.

In the genome-wide HG002 analysis, independent Hia5 MSP selection alone gave test-set event rates of 34.7% and 34.5% for CT and GA. Additional focal-site-excluded neighbor selection raised rates to 86.3–93.3%, depending on the threshold (Supplementary Fig. 1d).

### Complete-CpG-island methylation inference

Expanded whole-island validation used per-CpG methylation from HG002 LCL PacBio Fiber-seq (PS00166), a population reference, and all 12 matched scDAF-seq libraries. Methylation was called with pb-CpG-tools v3.0.0 (default settings) after pbmm2 alignment to the HG002 v1.1a assembly and lifted to the HG002 v1.1 diploid reference with per-20-kb offsets; 60,502,765 of 60,511,357 records (99.99%) were placed on a v1.1 CpG after offset placement, and the remainder were discarded. Islands required at least ten reference CpGs and mean methylation at most 0.10 or at least 0.90 for the extreme-state comparison. Entire chromosome groups, with both haplotypes assigned together, defined fixed training, validation and test sets.

CpG islands were inferred from the indexed reference using 200-bp windows stepped every 10 bp. Windows with GC fraction at least 0.50 and CpG observed/expected at least 0.60^41^ were retained, where observed/expected was the CpG count multiplied by window length and divided by the product of C and G counts; overlapping qualifying windows were merged. Users may instead supply a merged, nonoverlapping BED catalog. The supplied or inferred complete island remains the reporting unit in either case.

Initial MSPs were called with every CpG observation masked in both the first-pass HMM and the refinement steps. For each molecule-island overlap, only CpG and non-CpG SsDddA observations whose query positions lay within an initial FiberHMM MSP were retained. The non-CpG rate across the complete overlap supplied a molecule-specific accessible baseline, adjusted for the calibrated SsDddA 5′ sequence factors. CpG outcomes were summed under the calibrated methylated and unmethylated emission models with equal prior odds. Calls required at least 15 CpGs and 10 non-CpGs and were methylated at posterior at least 0.99, unmethylated at posterior at most 0.01 and otherwise uninformative. No transition or boundary was inferred within an island.

The default fiberhmm-tag-m5c workflow infers islands from the reference, writes the exact catalog on request and records confident methylated and unmethylated islands as MA:ddda_mcg and MA:ddda_ucg spans, respectively. An external--cpg-islands BED overrides the catalog without changing call resolution. The optional audit table retains all three molecule-island states and their evidence counts. SsDddA footprint recall enables CpG masking by default (--use-m5c--cpg-mask-policy unmethylated-only), retaining CpG observations only within confidently unmethylated MA:ddda_ucg spans and excluding all other CpGs from the informative-site set in both the first-pass HMM and refinement. Excluded sites contribute no likelihood, do not count toward minimum evidence, and cannot define footprint boundaries or boundary uncertainty; non-CpG observations retain their ordinary emissions. An alternative setting (--cpg-mask-policy methylated-only) instead excludes CpGs only within MA:ddda_mcg spans.

In a 1-kb regional CpG benchmark against the same reference across 46 haplotype contigs, the truth-window-weighted mean extreme-state AUROC was 0.9969 (contig range 0.990–0.999), and mean methylated- and unmethylated-call precision was 90.9% and 80.5% (Supplementary Fig. 7a). This benchmark ranks strongly methylated against strongly unmethylated reference windows. The whole-island classifier, which pools MSP evidence across a complete sequence-defined island without fitting internal boundaries, was evaluated separately. Calls required at least 15 informative CpGs; Supplementary Fig. 7c shows agreement and called fraction across evidence floors on validation chromosomes.

In the held-out whole-island evaluation, 26,179 overlaps with truth-unmethylated islands met the requirement of at least 15 informative CpGs and ten non-CpGs. Of these, 23,816 received a confident call (91.0%), and 23,746 agreed with the reference state (99.7%; 95% island-cluster bootstrap interval, 99.6–99.8%). Among 3,080 evidence-eligible overlaps with truth-methylated islands, 2,351 received a confident call (76.3%), and 2,322 agreed (98.8%; 95% interval, 97.9–99.4%). Because individual fibers need not match a population reference, the implied false-positive and false-negative rates (0.3% and 1.2%) are upper bounds on molecular calling error. Of confident predictions, 2,322 of 2,392 methylated (97.1%) and 23,746 of 23,775 unmethylated (99.9%) agreed with the reference. Footprint calling retains CpGs only in confidently unmethylated islands, so the unmethylated precision determines its accuracy; the lower methylated precision reflects the excess of unmethylated islands in the test set. Intervals were obtained from 1,000 Poisson island-cluster bootstrap resamples, retaining all molecules from each island together.

CpG specificity at matched local accessibility was assessed on the 3,860 evidence-eligible molecule–island overlaps (766 methylated-reference and 3,094 unmethylated-reference; one read that FiberHMM 3.0.4 flags as a PCR duplicate was excluded at extraction) from the test chromosome groups at mapping quality ≥30 across all 12 libraries, using raw aligned observations without restriction to an HMM state. For each focal CpG, local accessibility was the conversion of its ten nearest non-CpG observations within 100 bp, with observations required on both sides; focal non-CpG controls used the same rule with the focal site excluded, so a focal outcome never contributed to its own accessibility estimate. The primary analysis used focal sites with at least eight of ten neighbors converted. CpG and non-CpG conversion were standardized across shared strata of library, strand, 5′ base, local non-CpG activity and non-CpG activity on the same molecule outside the island (at least 20 observations required), retaining 179,332 of 211,823 site observations in 348 strata across 1,172 islands. The difference-in-differences was the methylated-minus-unmethylated change in CpG conversion minus the corresponding change in non-CpG conversion, with 95% intervals from 2,000 island-level bootstrap resamples. The primary estimate was −54.7 percentage points (95% interval −57.9 to −52.2; Supplementary Fig. 7e). Sensitivity analyses varied the neighbor requirement (6/10 and 10/10), radius (50 and 200 bp), mapping quality (≥60), strand, library and omitted test chromosome. Two held-out analyses assigned island state from a deterministic 80% of CpGs and measured conversion at the withheld 20%, using either initial MSPs or raw non-CpG neighbors alone (Supplementary Fig. 7e–g).

Geometry controls used 3,859 of these overlaps, with 148,203 CpG and 435,923 non-CpG initial-MSP observations. Methylated-reference islands were less accessible than unmethylated-reference islands (median island MSP fraction 0.27 versus 0.49; median linker 48 versus 62 bp), which these stratified controls address. The difference-in-differences was computed by the length of the MSP containing each observation (<100, 100–149, 150–199, 200–299, 300–499, 500–999 and ≥1,000 bp) and by tertiles of island MSP fraction, overlap-weighted MSP length, and median linker length, nucleosome repeat length and nucleosome-covered fraction within ±1 kb. Island state labels were shuffled 10,000 times within the 22 occupied strata formed by linker, MSP-fraction and MSP-length tertiles, and each shuffle was scored by the difference-in-differences computed within each stratum and averaged across strata (weights n_M_·n_U_/(n_M_ + n_U_)). The overall difference-in-differences across all initial-MSP observations was −40.9 percentage points (95% CI −42.4 to −39.3) and remained between −39 and −43 percentage points in every geometry tertile. The stratum-matched estimate was −40.2 percentage points, against a shuffle null centered on zero (mean −0.3; 95% range −3.3 to 2.7; P = 1 × 10⁻⁴, the smallest attainable with 10,000 shuffles).

### FiberBrowser interface

The FiberBrowser interface is a deployable application for inspecting and comparing single-molecule datasets. Indexed retrieval and display sampling limit rendering work, while sorting, filtering, clustering and co-occupancy use all eligible molecules and peak scanning uses the displayed molecules. Footprint inference is delegated to the shared FiberHMM engine, with stage progress, checkpoint reuse and report export exposed in the interface. Reversible curation, saved sessions and annotated-BAM export preserve an auditable path from the underlying observations to displayed results.

The FiberBrowser interface is implemented as a Python server with a browser-based front end and a Python client sharing one API; it is distributed with a Dockerfile and example datasets. Indexed BAM and bigBed molecule datasets can be displayed with BED, bigBed, bigWig^42^, GTF/GFF, FASTA and Hi-C reference tracks. Molecular Annotation (MA, AQ and AN), standard modification (MM and ML) and supported legacy tags are decoded into molecule objects carrying mapped blocks, informative sites, events, calls, scores and provenance. The molecules analyzed for the specified region are held independently of the subset sampled for rendering.

The genome panel supports multiple FASTA and annotation sets with distinct contig names, and searchable BED3, BED6 and BED12 annotations, including exon blocks. Individual windows can navigate independently using local gene or coordinate search. Clustering selected windows fits one shared model to common window-relative accessibility features, retaining assignments and source identities across chromosomes or haplotypes. Users supply comparable windows; the procedure does not infer coordinate correspondence or perform liftover.

Population operations run on the server-side working set: molecules are clustered by accessibility pattern, summarized as cluster metaprofiles and co-accessibility maps, and compared across datasets with source identity retained. Sequence and motif views, coverage-based autozoom and primer design support locus inspection. In the footprint panel, chemistry and platform are read from the established FIBERHMM-CHEMISTRY:v1: BAM header comments, with supported legacy metadata also recognized. Missing metadata disables inference and links to Dataset info, where the user can declare it and optionally persist canonical comments to the BAM. Conflicting metadata cannot be silently overridden. A compatible FiberHMM installation is required.

At individual regulatory elements, footprint inspection links call size, position and evidence to the underlying sequence; motif overlays and sequence export help interpret a footprint after it has been detected, without requiring a factor identity to define the original call. User-defined call labels and selected molecule groups carry into profiles, clustering and co-occurrence analyses, and every operation leaves the source data unchanged, except an optional, explicit write of chemistry metadata to the BAM header.

### Haplotype and inserted-sequence examples

Haplotype examples used haplotagged (HP) PacBio Fiber-seq alignments re-called with FiberHMM 3.0. FiberBrowser joint k-means clustering (K = 2) used accessibility over the *SNRPN* differentially methylated region core (chr15:24,823,417–24,824,334) or the *RPLP0* promoter core (chr12:120,200,910–120,201,310; H1 Fiber-seq; accessible cluster 41/54 HP1 and 46/58 HP2 molecules) in both haplotype panels; per-haplotype cluster counts were compared by two-sided Fisher’s exact test. CpG methylation used the PacBio 5mC calls in the input BAM, with a CpG counted as methylated at ML ≥ 128. Allelic footprints were screened at phased heterozygous SNVs within JASPAR 2024 motif cores (log2 odds against a uniform background with 0.1 total pseudocount per position and exact score P values; tested when one allele scored P < 10⁻⁴ and the other lost at least 3 bits). A molecule spanning the motif was counted as bound when a single FiberHMM footprint covered the complete motif core, and haplotypes were compared by two-sided Fisher’s exact test with Benjamini–Hochberg correction across all 8,055 tests in GM12878 PacBio Fiber-seq (Supplementary Table 8). Replication used the same motif coordinates in the H1, H9, K562 and HepG2 libraries heterozygous for the variant. For display, molecules were split by the base at the variant with the FiberBrowser variant scan (minor-allele fraction at least 0.1, depth at least 20, both strands). Insertions of at least 100 bp supported by at least eight molecules (MAPQ ≥ 30) were identified from CIGAR operations, and the inserted sequence was taken as the consensus of carrier molecules, which FiberBrowser expanded as a separate allele for display. Insert accessibility was measured from FiberHMM MSPs in insert coordinates, and a nucleosome-free region was counted on a molecule when an MSP of at least 150 bp covered it. The footprint sequence was scanned with JASPAR 2024 CORE vertebrate matrices^43^ (pseudocount 0.8; background A and T 0.3, C and G 0.2), with P values from the exact score distribution.

### Promoter CNN cell-identity transfer

A five-class promoter classifier was trained on Hia5 Fiber-seq from GM12878 (LCL), H1, H9, K562 and HepG2 and applied to 12 individual HG002 scDAF-seq cells^15,44^. Validation used independent H1, H9 and K562 libraries and parent-molecule-disjoint 80:20 training–validation splits for GM12878 and HepG2. scDAF-seq molecules were reconstructed as physical duplexes and processed with the nucleosome and footprint callers described above, as implemented in FiberHMM 2.16.8 at LLR 7 with the original within-cell duplex assignments; the Hia5 inputs were called with FiberHMM 2.13.4–2.16.3. The classifier was not retrained for later FiberHMM releases.

Single-molecule inputs were 5-kb hg38 promoter windows oriented by transcription-start-site strand; the training panel contained 101,318 supported windows. Each observation was required to span both window ends, cover at least 4,950 bases and contain no unobserved gap longer than 30 bp. At 1-bp resolution, the input encoded nucleosome and subnucleosomal protection, soft encodings of nucleosome-free gap length, reference A/C/G/T sequence gated by footprints or regulatory gaps, and observation validity. The same reference sequence was used for both assays.

The residual CNN^45,46^ used 32 channels and nine kernel-9 convolutional blocks with dilations 1, 2, 4, 8, 16, 32, 64, 64 and 128, giving a receptive field spanning the 5-kb input. Mean and maximum features across 25 positional bins, together with feature fractions, were projected to a 32-dimensional local representation, and learned locus-specific weights related these representations to the five reference classes. Contributions were averaged within promoters, within overlapping promoter groups and across observed groups to obtain one prediction per cell.

Synthetic reference cells were assembled by sampling one or two independent molecules per promoter, with promoter coverage varied during training. Training minimized class-balanced cross-entropy with AdamW (learning rate 5 × 10^−4^; weight decay 0.01). Checkpoints were selected solely by mean cross-entropy across ten fixed Hia5 validation profiles, comprising one- and two-molecule profiles for each reference class; scDAF-seq predictions did not enter fitting or model selection. The selected footprint model was obtained after 1,540 optimizer updates. Its expanded reference evaluation used 160 fixed pseudo-cells: four deterministic draws for each class, one- or two-molecule sampling and promoter retention of 8%, 25%, 50% or 100%. Each scDAF-seq cell was classified separately from its eligible duplex observations without fitting any classifier parameter to scDAF-seq. Class scores are softmax-transformed logits, and the LCL margin is the LCL logit minus the largest competing-class logit.

### Raw-observation comparisons

The matched raw-input CNN replaced FiberHMM call channels with binary modification events, assay-specific observable-substrate masks, reference A/C/G/T sequence and observation validity. Raw events remained at their original genomic positions without smoothing, rate normalization, quality-score channels or inferred calls. Architecture, promoter-spanning molecules, reference splits, synthetic-cell sampling, batch size and initial optimizer settings matched the footprint CNN. The raw checkpoint was selected solely on held-out Hia5 reference data at update 3,780; training stopped at update 3,798 after reference accuracy saturated. Cumulative logged training and loading time to the selected checkpoint was 157.4 h for raw input and 45.9 h for footprint input. scDAF-seq was not used for training or checkpoint selection. Training histories for both CNNs are shown in Supplementary Fig. 10d.

The methylation-density control CNN used the architecture, promoter windows, reference splits, class map, synthetic-cell sampling and optimization of the footprint CNN, but its input replaced the FiberHMM call channels with the fraction of modified informative sites in a centered 51-bp running window, standardized with per-assay means and standard deviations (Hia5 values from the Hia5 training libraries; SsDddA values from pooled unlabeled scDAF-seq molecules). It was trained on Hia5 Fiber-seq only, checkpoints were selected by cross-entropy on the same ten fixed Hia5 validation profiles, and scDAF-seq labels were never used. Training ran without a stopping rule to update 1,537, with no improvement after update 700; the best checkpoint (update 700) is shown.

### Molecular sampling requirements for cell identification

Figure 5b compares the three CNNs at 100% promoter retention using 40 matched pseudo-cells (four draws for each of five classes at one- and two-molecule sampling). The fixed reference evaluation also included 8%, 25% and 50% retention, giving 160 held-out pseudo-cells: the footprint CNN classified 151 (94.4%) correctly, with all nine errors at 8% retention and all 120 pseudo-cells at 25–100% retention correct, and the raw-input CNN classified 158 (98.8%) correctly. The methylation-density CNN classified 24 of the 40 full-retention pseudo-cells (60%) and 61 of all 160 (38%) correctly.

Transfer negative controls applied the frozen footprint CNN to the same 12 scDAF-seq cells and eligible duplex observations with degraded inputs. Nucleosome and footprint calls were re-placed at random within each molecule, or calls and the gaps between them were reordered, preserving per-base call content; molecules were assigned to permuted promoter slots, keeping either their own reference sequence or that of the new slot; or only the observation-validity mask and reference sequence were supplied, with no calls or with every observed base called nucleosome. Stochastic controls were drawn 20 times per cell (240 predictions each), and deterministic controls were run once per cell. With empty input, the learned class prior was uniform (softmax 0.20 per class).

To measure prediction at sparse molecular depth, the fitted footprint CNN was held fixed and applied to 20 deterministic nested molecule subsets of each of the 12 HG002 scDAF-seq cells at 0.01%, 0.02%, 0.05%, 0.1%, 0.2%, 0.5%, 1%, 2%, 5%, 10%, 25%, 50% and 100% of eligible promoter-spanning molecules, retaining all promoter windows of a physical duplex together and aggregating contributions as for complete cells (240 predictions per depth). The footprint CNN assigned all 12 complete cells to LCL; at 5% and 1% of molecules, 236 and 201 of 240 predictions were lymphoblastoid, and at 0.01%, with one or two retained molecules per cell, 58 of 240 (Fig. 5c). The raw-input CNN, evaluated on matched selections at 1–100% input, assigned 22.5%, 5.4% and 0% of predictions to LCL at 1%, 25% and full input, compared with 99.6% and 100% for the footprint CNN at 25% and full input. Bands in Fig. 5c show the 2.5th–97.5th percentiles across draws. Model architecture, training and checkpoint selection are described above and in Supplementary Fig. 10a–d.

### Model training records and supplementary diagnostics

The promoter CNN architecture, classwise reference confusion and optimization history are shown in Supplementary Fig. 10a–c. The confusion matrix uses the 160 fixed reference pseudo-cells (32 per class). The history distinguishes individual training losses, their 28-update rolling mean and selection cross-entropy on ten fixed Hia5 profiles. The selected update was 1,540 and training stopped at update 1,680. Selection loss and expanded 160-profile evaluation loss describe different cohorts.

### Statistics and reproducibility

Resampling and validation respected the measurement unit. Controlled-recovery intervals resampled genomic context blocks; SsDddA emission models left out entire scDAF-seq libraries, with covariance or bootstrap resampling grouped by physical duplex or library as specified for each analysis. Motif footprint comparisons with ChIP-seq used site-level scores with 2,000 whole-genomic-block bootstrap samples for uncertainty and paired AUROC differences. Source tables retain the prespecified metrics and all effect directions, and commands, seeds, input and model hashes, cohort definitions and exclusions are recorded in machine-readable manifests. Software tests covering saved numerical replays, MA/AQ/AN BAM round trips, reverse-orientation modification handling, masked duplex recall and browser workflows are included in the repositories.

## Data availability

Hia5 PacBio Fiber-seq of the ENCODE cell lines generated for this study (H1, H9, K562 and HepG2; 28 runs) is available from the NCBI Sequence Read Archive under BioProject PRJNA1417779. The new Drosophila data will be deposited in the Sequence Read Archive before publication: *Drosophila* DddB *hb*, *ind*, *scyl* and *chrb* amplicons, the *Drosophila* yw 2–4-h embryo Hia5 ONT Fiber-seq dataset and its naked-DNA and untreated controls, the DddB naked-DNA controls (whole-genome-amplified and unamplified), and the Hia5 PacBio 2–4-h Fiber-seq reads at the four amplicon loci, a subset of the library described in ref. 38. Submitted BAM files retain base-modification (MM/ML) tags, which are kept in the original-format files but are not included in normalized SRA downloads. Hia5-treated and untreated Drosophila S2 genomic DNA PacBio controls are available from GEO (GSE249742).

Published scDAF-seq (BioProject PRJNA1203351; runs SRR33130336–SRR33130342, SRR33130350 and SRR33130359–SRR33130362) and *NAPA*/UBA1 datasets are from Swanson et al.^15^. Previously published GM12878 Fiber-seq data are described in the cited Fiber-seq studies^20,21^. CTCF ChIP-seq peak sets were obtained from ENCODE^35^ (K562, ENCFF430KTH; GM12878, ENCFF635MMB; H1, ENCFF414GZI; H9, ENCFF152GTF; HepG2, ENCFF008COY). NRF1, REST and USF1 ChIP-seq peak sets (conservative IDR peaks) were also obtained from ENCODE (NRF1: K562, ENCFF777PKJ; GM12878, ENCFF864EBU; H1, ENCFF780FSK; HepG2, ENCFF785BAI; REST: K562, ENCFF649HPP; GM12878, ENCFF348LKE; H1, ENCFF755NTJ; HepG2, ENCFF227COI; USF1: K562, ENCFF913XQO; GM12878, ENCFF688QPS; H1, ENCFF971AKN; HepG2, ENCFF048LKP). HG002 LCL Fiber-seq (PS00166) is publicly available as part of the HG002 FIRE dataset (s3://stergachis/public/FIRE/broadly-consented/)^21^; per-CpG methylation calls (pb-CpG-tools v3.0.0, HG002 v1.1a) were generated as in Tullius et al.^47^. Plotting tables, model configurations and analysis provenance accompany the source data.

## Code availability

FiberBrowser (interface and Python API) is released and available at https://github.com/fiberseq/FiberBrowser under the MIT license. Its analytical engine, FiberHMM, including the calling, pairing and CpG tools, is released and available at https://github.com/fiberseq/FiberHMM under the MIT license. The repositories provide installation instructions and workflow documentation. Software versions and model identifiers for individual analyses accompany the source data.

## Acknowledgements

We thank K. M. Munson and the University of Washington Northwest Genome Center for PacBio sequencing, and Bernard Kim and members of his laboratory in the Department of Ecology and Evolutionary Biology, Princeton University, for use of their PromethION. We thank the ∼60 scientists who tested FiberBrowser and FiberHMM, especially members of the Levine and Stergachis laboratories, Lucia Ichino, Lakshmi Mahendrawada and Artem Yaschenko. This work was supported by NIH grants R35 GM118147 (to M.S.L.) and 1DP5OD029630, UM1DA058220 and 1U01HG013744 (to A.B.S.). A.B.S. holds a Career Award for Medical Scientists from the Burroughs Wellcome Fund and is a Pew Biomedical Scholar.

## Author contributions

T.W.T. conceived the study, developed FiberHMM and FiberBrowser, generated data, performed all analyses and wrote the manuscript. M.R.V. developed the Molecular Annotation specification. C.C., J.R., J.H. and M.R.V. provided data. A.B.S. and M.S.L. supervised the study. All authors reviewed the manuscript.

## Supplementary Information

**Supplementary Figure 1.**
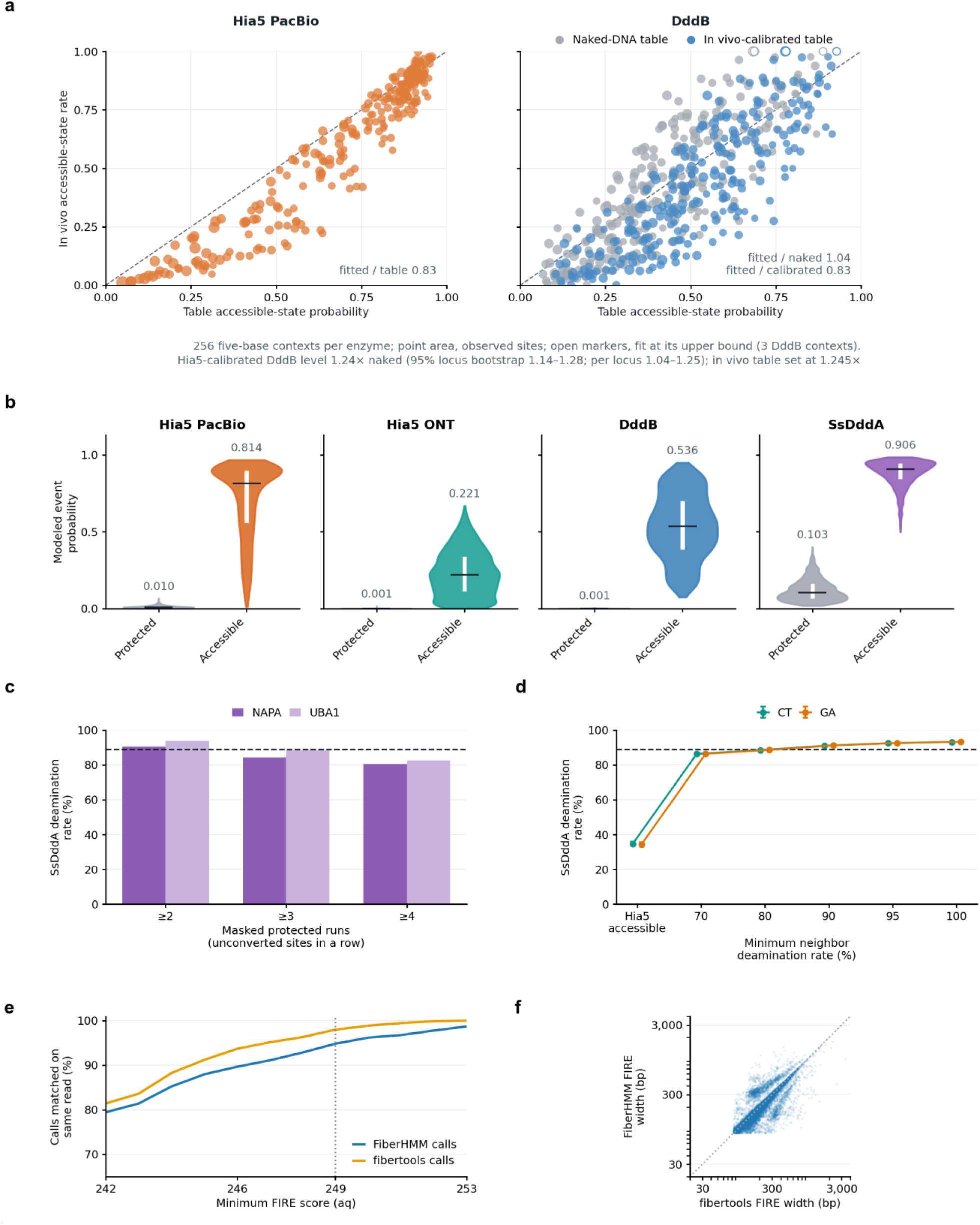
Emission calibration and FIRE interoperability. a, In vivo accessible-state calibration. In vivo accessible-state rates by five-base context against the emission tables, at the *hb*, *ind*, *chrb* and *scyl* amplicons (DddB, 22 amplicon libraries from wild-type 1.5–4-h embryos; Hia5 PacBio 2–4 h, 11 movies). For each enzyme, the accessible-state rate per context was fitted as P(event) = a·p_acc_ + (1 − a)·p_prot_, where a is the fraction of molecules from an independent half of the same enzyme’s data whose MSP covers the site outside called footprints. Hia5 (left) reached 0.83 of its table in this per-context fit; DddB (right) exceeded the naked-DNA table 1.04-fold (gray) and reached 0.83 of the in vivo-calibrated table (blue). Relative to Hia5, this places DddB at 1.24× the naked table (95% locus bootstrap 1.14–1.28; per locus 1.04–1.25), consistent with the 1.245× table level set from the Hia5 modification-burden comparison (Methods). 256 five-base contexts per enzyme; point area, observed sites; open markers, fits at the upper bound (3 DddB contexts). **b, Sequence-context emission distributions.** Modeled event probabilities for all 4,096 contexts of each chemistry, not weighted by how often each context occurs; gray, protected; colored, accessible; black line, median; white bar, interquartile range. **c, Hia5-accessible-region-based SsDddA rate sensitivity. *NAPA* and *UBA1* deamination rates after excluding runs of two, three or four unconverted SsDddA sites.** The dashed line marks the reported physical-duplex accessible rate of 89%. **d, Genome-wide Hia5-guided rate sensitivity.** Test-chromosome CT and GA event fractions under the independent Hia5-accessible mask alone and with additional focal-site-excluded SsDddA neighbor-rate thresholds. All 6,022 genomic blocks and twelve matched scDAF-seq libraries entered the predefined splits. Error bars are 95% chromosome-group bootstrap intervals, with both haplotypes resampled together. The dashed line marks the 89% duplex reference. **e, Same-read FIRE concordance versus minimum FIRE score (aq). Each curve is the fraction of one caller’s FIRE calls matched by a positive FIRE call from the other segmentation on the same read, among 9,781 common LCL (GM12878) reads. At aq ≥ 249, concordance is 98.0% for fibertools and 94.8% for FiberHMM.** Final MA:msp coordinates match FIRE’s as/al input exactly. **f, Width agreement of shared FIRE calls.** Paired widths for 17,340 one-to-one shared calls; the diagonal marks equality. Median absolute difference is 8 bp (interquartile range 4–21 bp). Axes are logarithmic.

**Supplementary Figure 2.**
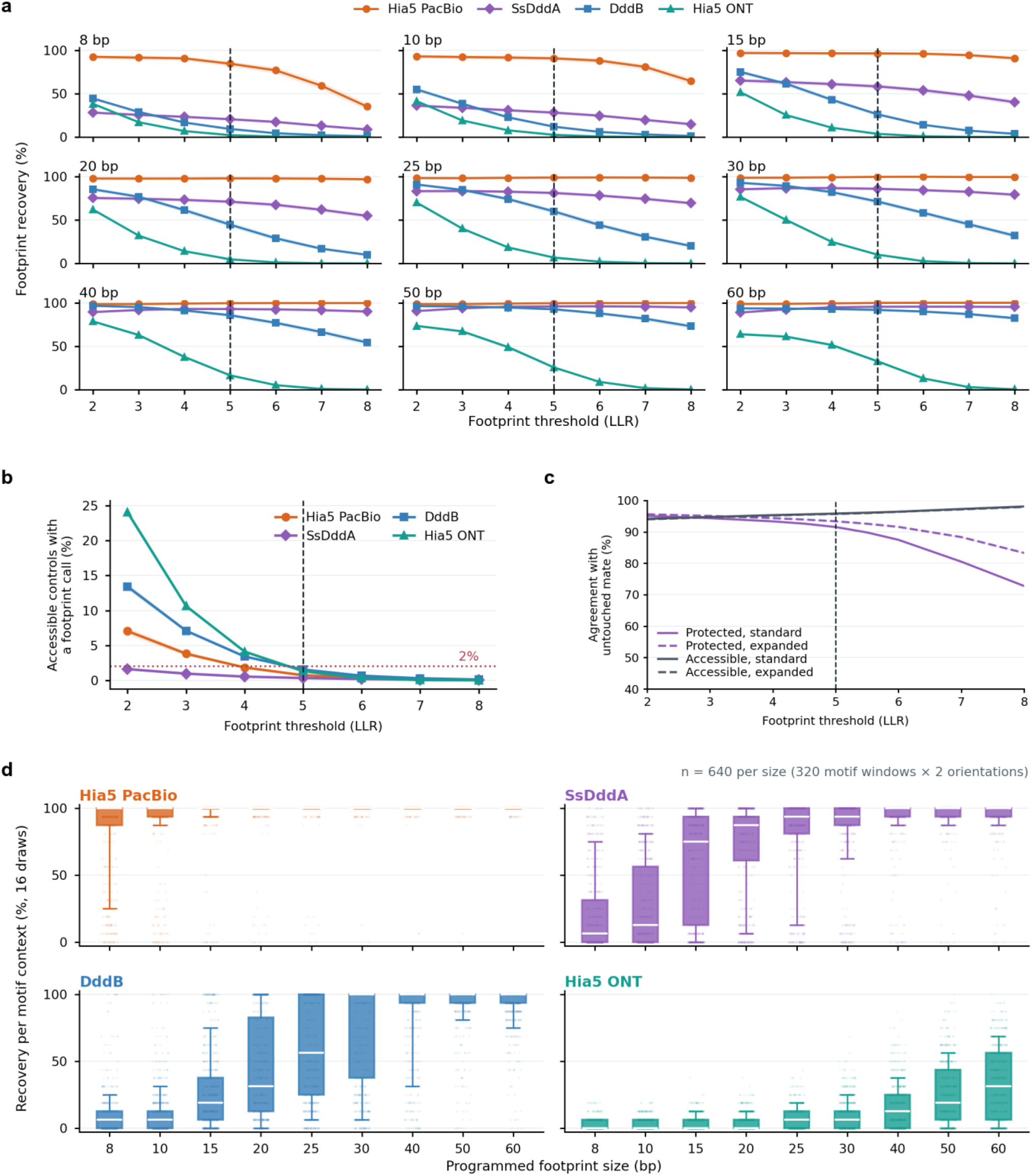
Footprint threshold and recovery benchmarks. **a, Protected-target LLR curves.** Footprint recovery (target center covered) across LLR 2–8 for each programmed footprint size from 8 to 60 bp. Each chemistry and threshold uses 320 genomic motif contexts, two orientations and 16 draws per size. Shading gives 95% genomic-block-bootstrap intervals, and the vertical guide marks the shared LLR 5 setting. **b, Pooled accessible-control curve.** Matched fully accessible windows are pooled across the nominal target-size strata because no protected target exists in these controls. The horizontal guide marks 2%; LLR 5 is the first shared integer setting below 2% for every chemistry. **c, Expanded partner-strand calibration.** Default-decoder protected-window recovery and accessible-window specificity on partner strands from two held-out libraries, comparing standard emissions (solid) with expanded exact six-base emissions (dashed); purple, protected-window recovery; gray, accessible-window specificity. The full cohort comprises 541,821 duplexes from twelve libraries with an eight/two/two training/validation/test split. Test duplexes overlapping chr1:1–40 Mb, the region used to fit the standard emissions, were excluded, leaving 79,543 protected and 34,241 accessible windows on 42,650 duplexes. Shading gives 95% physical-duplex-cluster bootstrap intervals, conditional on the two test libraries; the vertical guide marks LLR 5. Both query directions are pooled. **d, Context-level recovery distributions at LLR 5.** Per-context recovery of a subnucleosomal footprint covering the programmed center, the endpoint of a, across 16 draws for all four chemistries at LLR 5 and every programmed footprint size. Each size has 640 units (320 motif windows × 2 orientations), each recalled over 16 draws.

**Supplementary Figure 3.**
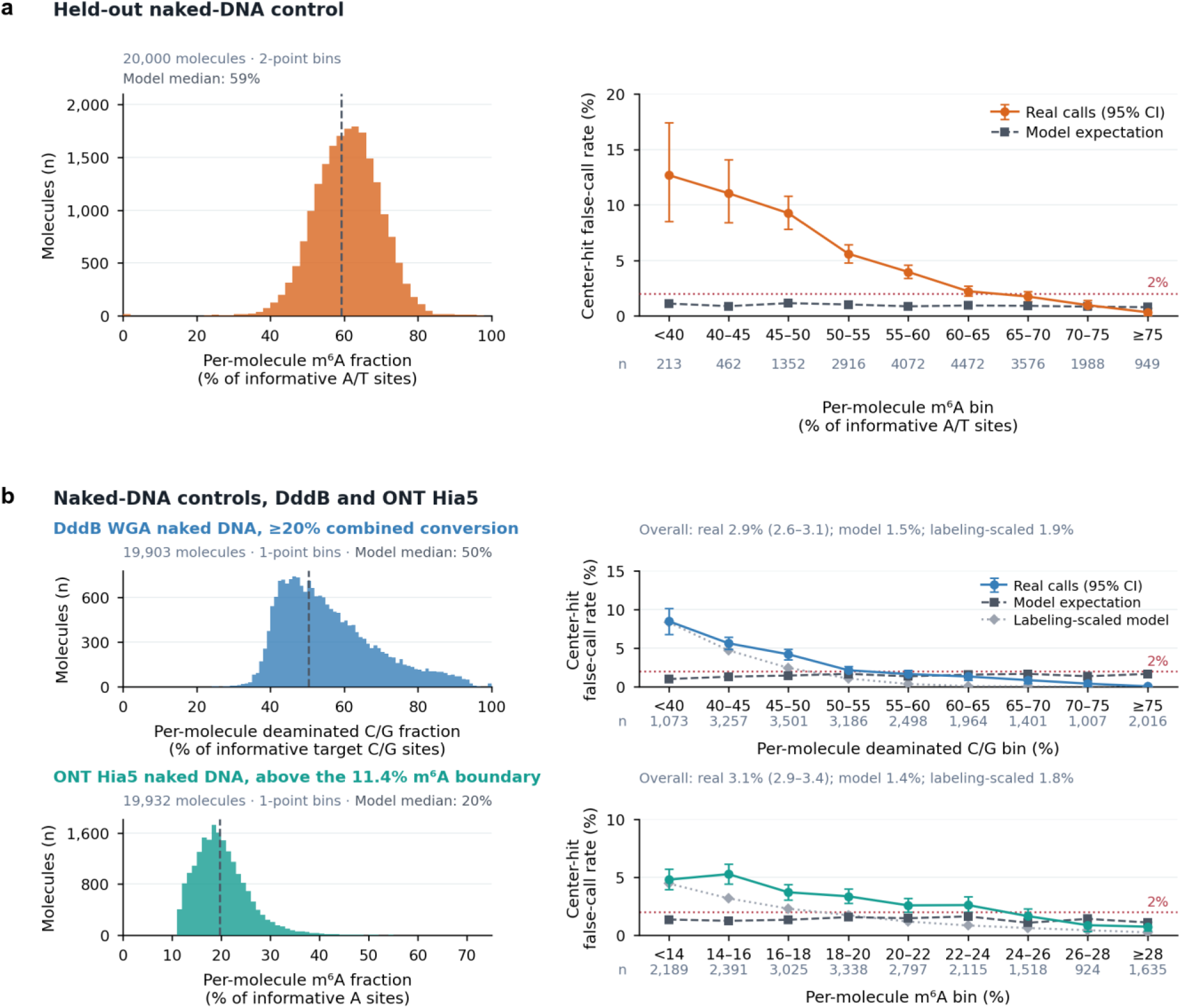
False footprint calls in naked-DNA controls. **a, Held-out naked-DNA control.** Left, per-molecule m6A fraction in 20,000 Hia5-treated naked-DNA molecules not used for emission fitting; dashed line, median of the model expectation. Right, fraction of fixed 800-bp windows with a footprint covering the center at LLR 5, by per-molecule m6A fraction, for real calls (95% molecule-bootstrap intervals) and for observations drawn from the accessible-state emissions at the same sites; n, molecules per bin. Overall rates are 3.6% and 0.9%. **b, Naked-DNA controls for DddB and Hia5 ONT, analyzed as in a.** Top, DddB WGA naked DNA with ≥20% combined conversion; bottom, Hia5 ONT naked DNA above the 11.4% m6A boundary (20,000 molecules sampled per set; 19,903 and 19,932 decoded by the caller). Left, per-molecule labeling (fraction of informative target C/G sites deaminated, or of informative A sites called m6A); dashed line, median of the model expectation. Right, fraction of fixed 800-bp windows with a footprint covering the center at LLR 5, by labeling bin, for real calls (95% molecule-bootstrap intervals), for draws from the accessible-state emissions at the same sites (mean of three) and for draws scaled to each molecule’s labeling (mean of ten); n, molecules per bin; dotted line, 2%. Overall rates are 2.9% (model 1.5%, labeling-scaled 1.9%) for DddB and 3.1% (1.4%, 1.8%) for Hia5 ONT. Unlike the held-out molecules in a, these molecules overlap the data used to fit the emissions.

**Supplementary Figure 4.**
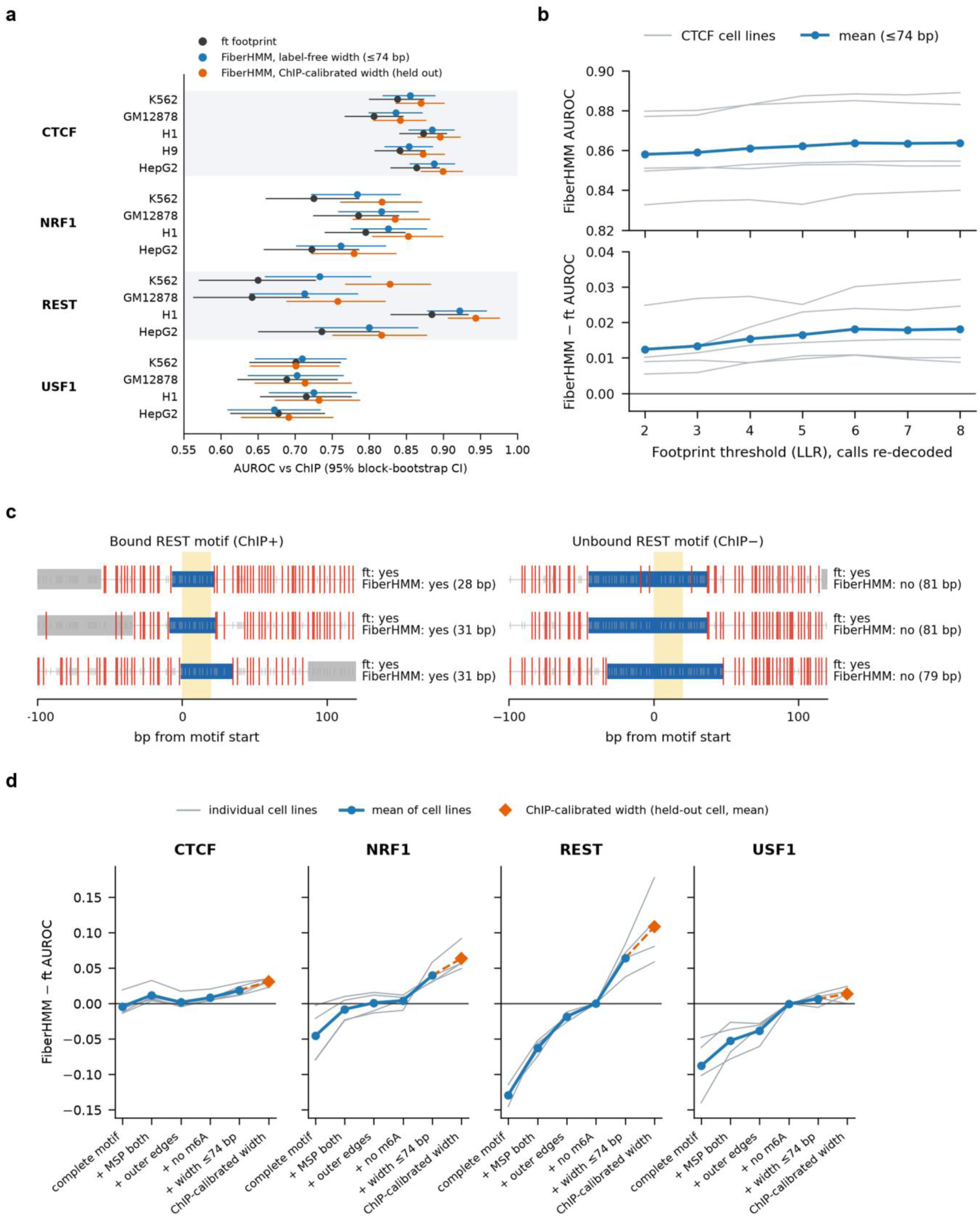
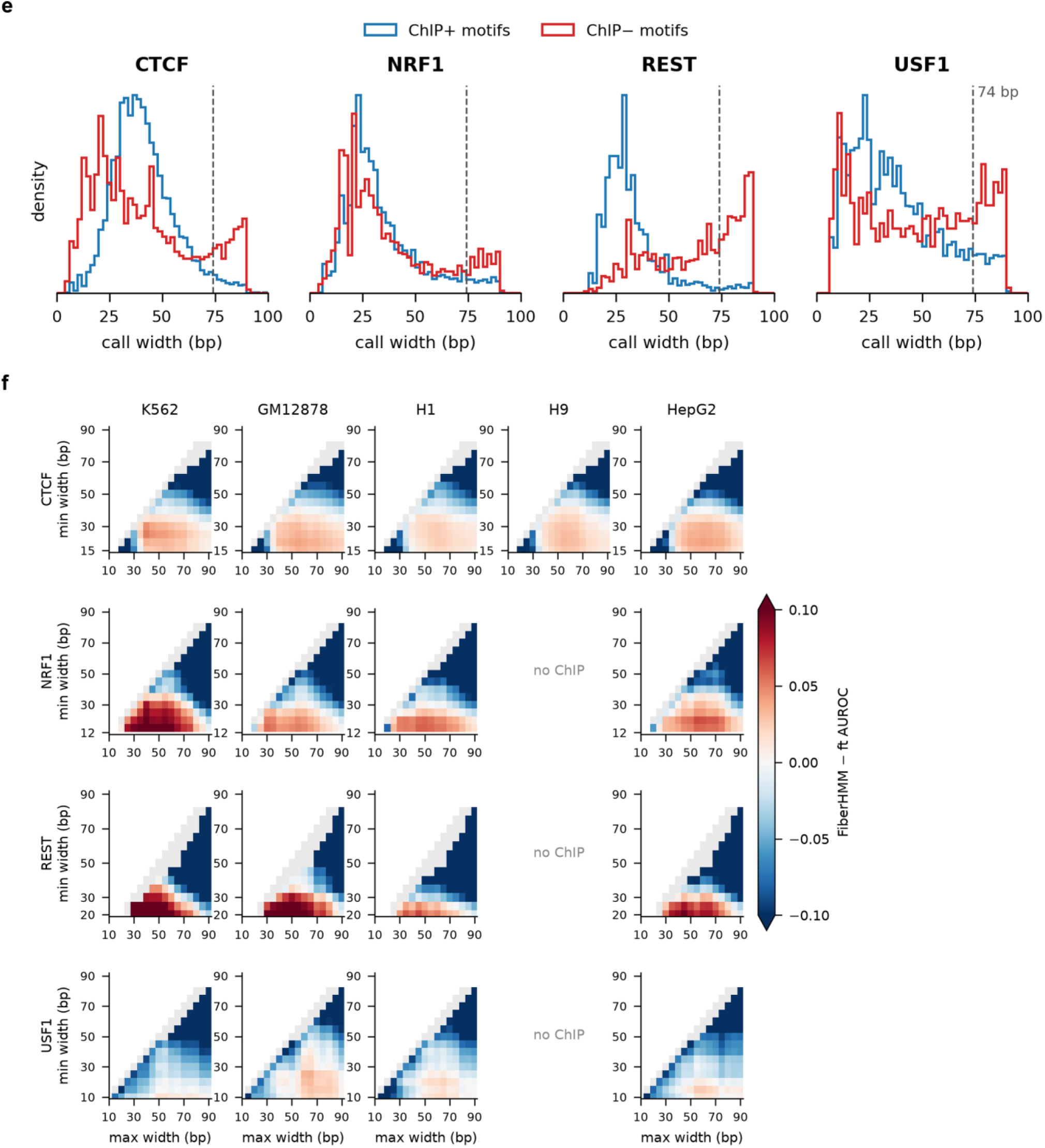
Motif footprint benchmark against ChIP-seq. **a, Per-cell-line AUROC.** AUROC (95% genomic-block-bootstrap interval) of per-motif bound fractions against cell-matched ChIP-seq for ft footprint (gray), fiberhmm-footprint with the label-free 74-bp maximum width (blue) and fiberhmm-footprint with a ChIP-calibrated window learned in the other cell lines of the same factor (orange). All callers use the same MSP-eligible molecule–motif observations. CTCF, 512 motifs in five cell lines; NRF1, REST and USF1, 254–256 motifs in four (H9 lacks ChIP-seq for these factors). **b, Footprint threshold.** CTCF FiberHMM AUROC (top) and FiberHMM − ft difference (bottom) with the label-free 74-bp maximum width, for calls re-decoded at LLR 2–8 on the fixed LLR 5 eligible observations; gray, individual cell lines; blue, mean of five cell lines (Supplementary Table 3). **c, Example molecules.** Three H1 molecules at a ChIP-supported (left) and an unsupported (right) REST motif (yellow). Blue, the motif-containing FiberHMM footprint call drawn at its outer edges; gray blocks, nucleosome calls; red ticks, m6A; thin gray ticks, unmethylated informative sites. ft footprint counts all six molecules as bound because the motif is unmethylated; fiberhmm-footprint counts the 28–31-bp calls at the bound motif and rejects the 79–81-bp protections at the unbound motif, which exceed the 74-bp maximum. Labels give conservative call widths. **d, Rule components.** Per-cell-line (gray) and mean (blue) FiberHMM − ft AUROC as scoring criteria are added in turn: a footprint containing the complete motif; the MSP requirement applied to both callers; containment within the call’s outer edges; no m6A in the motif; the label-free 74-bp maximum width; and the ChIP-calibrated window evaluated on held-out cell lines (orange). Without a width limit FiberHMM matches ft footprint (identically for REST and USF1); the width step provides the gain. **e, Footprint widths at motifs.** Conservative widths of motif-containing FiberHMM footprint calls at ChIP-supported (blue) and unsupported (red) motifs, pooled across cell lines (unmethylated motifs on MSP-eligible molecules). Dashed line, the label-free 74-bp maximum. ChIP labels are shown for display only; the cap was set without them. **f, Width-window surfaces.** FiberHMM − ft AUROC across [minimum, maximum] width windows for each factor and cell line, as in Fig. 1e. Gray, windows with FiberHMM positives in <1% of observations or at <20% of motifs (pooled across the factor’s cell lines); white, minimum above maximum; no ChIP, factor not profiled in that cell line. Color scale clipped at ±0.10.

**Supplementary Figure 5.**
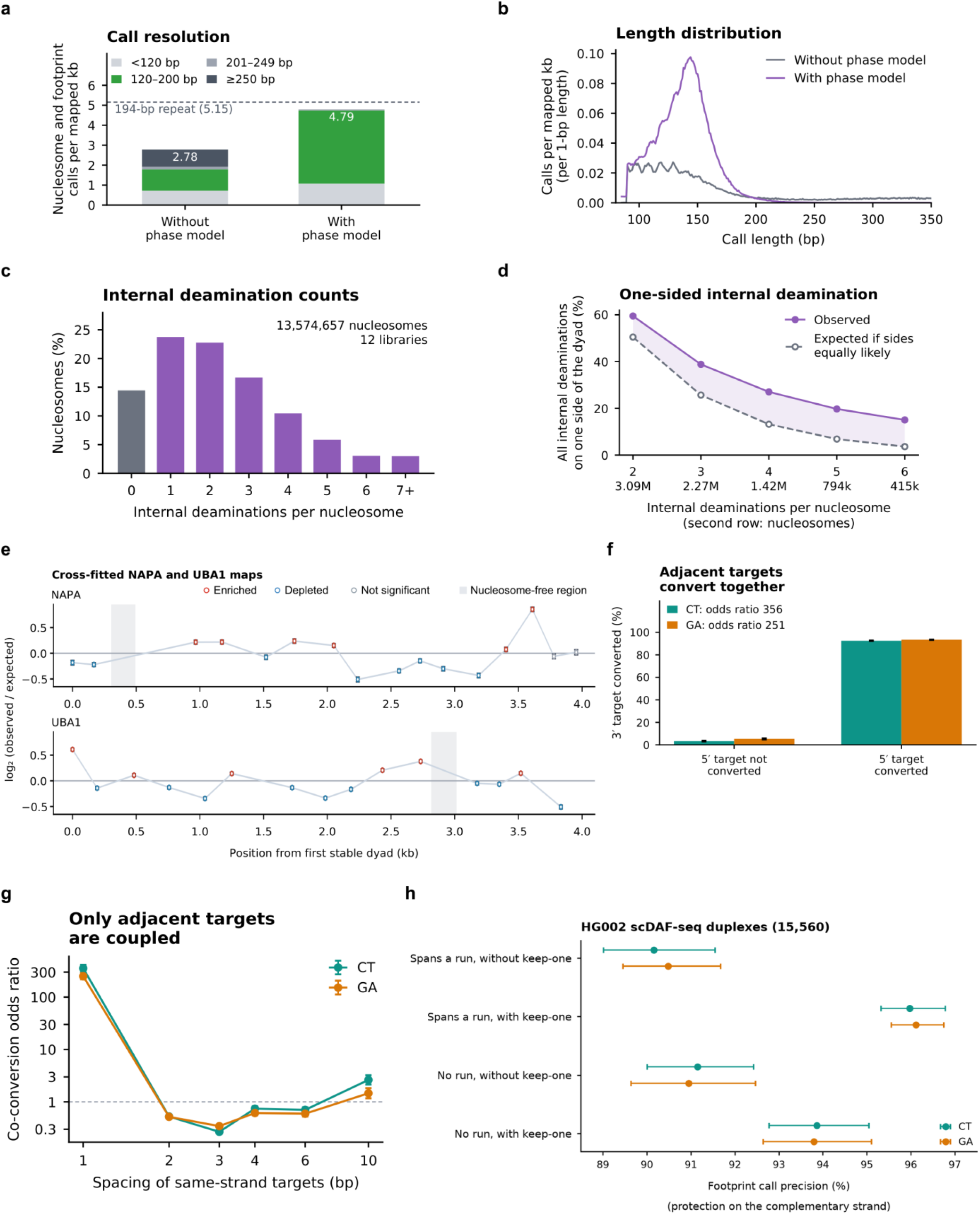
SsDddA nucleosome inference and adjacent-target dependence. **a, Nucleosome call resolution.** Nucleosome and footprint calls per mapped kb without or with the phase model, separated by span. The dashed line is 1,000/194 = 5.15 calls per kb, a spacing reference for a 194-bp repeat. **b, Length distribution with the phase model.** Length distributions of nucleosome and footprint calls from the same molecules without or with the phase model. **c, Internal event-count distribution.** Internal deamination counts among all 13,574,657 eligible nucleosomes from 12 libraries, including 1,961,459 with no internal event (14.4%). **d, Internal-event sidedness.** Nucleosomes with all internal deaminations on one side of the called dyad, compared with the expectation if both sides are equally likely, for two to six events; the second row of x labels gives the number of nucleosomes in each group. **e, Cross-fitted *NAPA* and *UBA1* maps.** Cross-fitted fixed-window event burden (recurrent centers nominated from called particles in the opposite molecule half; all spanning molecules evaluated) at recurrent nucleosome sites, shown as log2(observed/expected). Error bars are the molecule-bootstrap 95% intervals. Red and blue indicate significant enrichment and depletion, respectively, at family-wise corrected P < 0.05; gray indicates nonsignificant positions. Gray bands mark the annotated nucleosome-free regions (NFRs) at *NAPA* chr19:47,515,050–47,515,240 and *UBA1* chrX:47,193,580–47,193,780. Positions are measured from the first stable dyad in each amplicon. **f, Adjacent targets convert together.** Conversion of the 3′ target of isolated adjacent same-strand pairs (CC on CT reads, GG on GA reads), conditional on conversion of the 5′ target in the deaminated strand’s orientation, at confidently accessible sites of *NAPA* SsDddA molecules (inside an MSP and at least 10 bp from any footprint or nucleosome call; unthinned informative-site sets). Odds ratios 356 (CT; 42,871 pairs) and 251 (GA; 40,028 pairs). **g, Adjacent targets are by far the most strongly coupled.** Co-conversion odds ratio against the spacing of consecutive same-strand targets with no target between them. Shared local chromatin state would couple nearby targets at every short spacing; instead coupling is by far strongest for adjacent targets, and pairs 2–6 bp apart are anti-correlated (spacings tested: 1, 2, 3, 4, 6 and 10 bp; odds ratios at 10 bp, 2.6 for CT and 1.5 for GA). **h, Physical-duplex precision.** Mate-likelihood-estimated precision (%) of native SsDddA footprint calls in 15,560 sequence-assigned HG002 scDAF-seq physical duplexes, read as protection on the complementary strand of the same molecule, for calls that do or do not span same-strand runs, without and with keep-one. CpGs excluded; library-bootstrap 95% intervals.

**Supplementary Figure 6.**
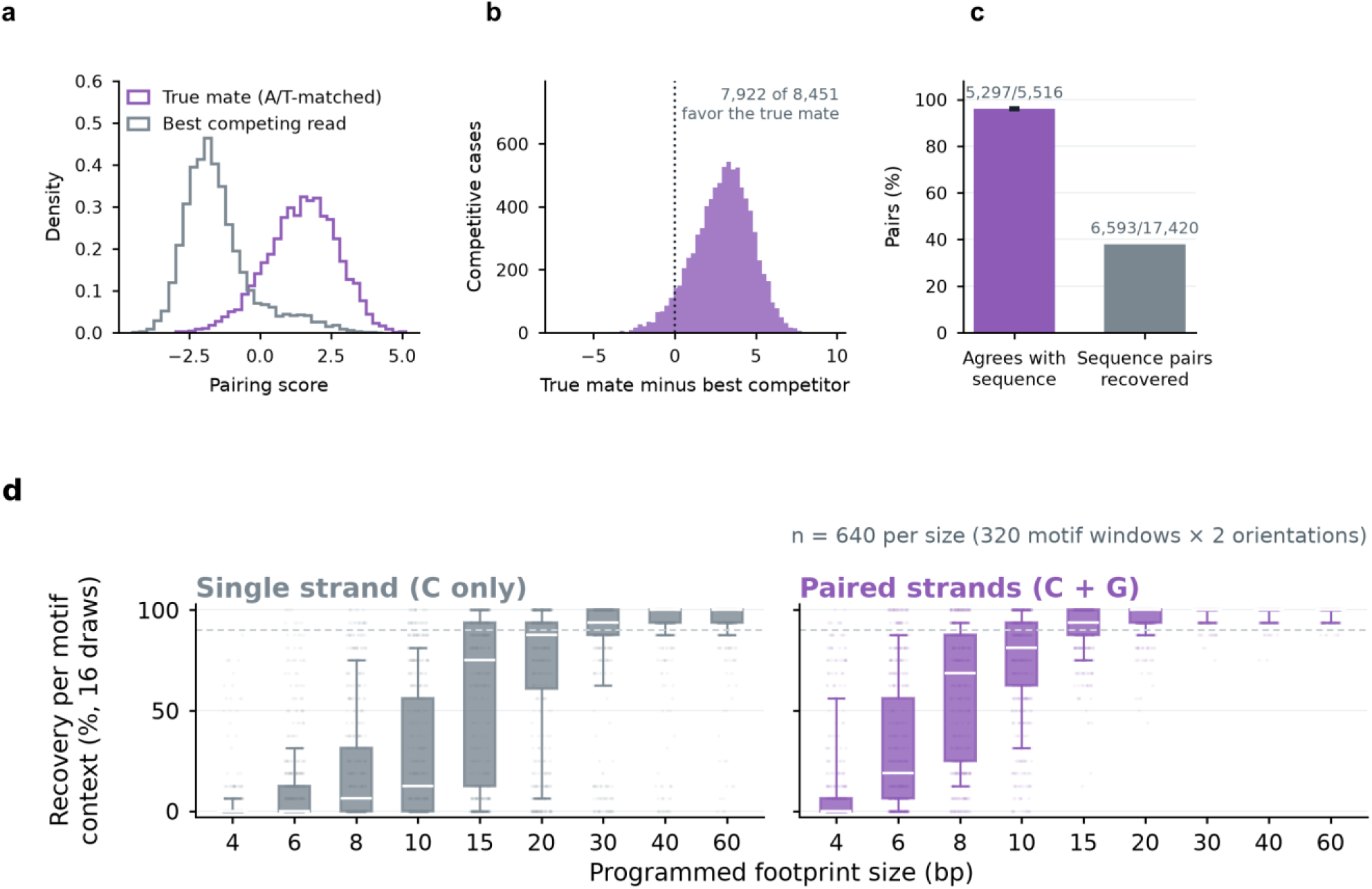
Physical-duplex pairing and resolution. Validation of footprint-only pairing against independent sequence-supported mate assignments. **a, Pairing score distributions.** Fixed footprint-only scores for independently A/T-selected mates and their best read-sharing competitors in 8,451 competitive cases where both reads map uniquely to the same HG002 haplotype, across the twelve-library evaluation cohort. A/T identity supplies only the evaluation label; pairing-model fitting regions are excluded. **b, Competitive score differences.** Mate score minus the best competing score in the same cases; 7,922/8,451 favor the independently selected mate. **c, Pairing agreement and recovery.** At loci where both reads map uniquely to one HG002 haplotype, so that sequence can distinguish the mate from the other homolog’s strand, 5,297 of 5,516 sequence-assessable footprint-selected pairs agree with the independent A/T relationship (96.0%; 95% connected-component bootstrap interval, 95.5–96.5%). Recovery is 6,593/17,420 among all independently A/T-selected pairs, including footprint-unscorable pairs. The full cohort contained 511,214 footprint-selected pairs: 503,763 (98.54%) lacked a decisive A/T mate assignment, and 76.97% lacked a homolog comparison. Sequence and haplotype were withheld from footprint pairing. **d, Resolution gained from complementary strands.** Context-level recovery distributions from 320 genomic motif windows in two orientations, with 16 draws per unit, called with the same default SsDddA pipeline as Supplementary Fig. 2a,d (HMM, rotational nucleosome refiner and footprint refiner) at LLR 5 with at least three informative sites; single-strand values are those of Supplementary Fig. 2a,d. Pairing combines the complete C+G informative-site set instead of calling only the C-bearing strand. Pooled recovery increases from 20.5% to 57.8% at 8 bp, 28.1% to 75.3% at 10 bp, and 58.2% to 92.1% at 15 bp; boxes show quartiles, whiskers the 10th–90th percentiles, and points all 640 units (320 motif windows × 2 orientations), each recalled over 16 draws.

**Supplementary Figure 7.**
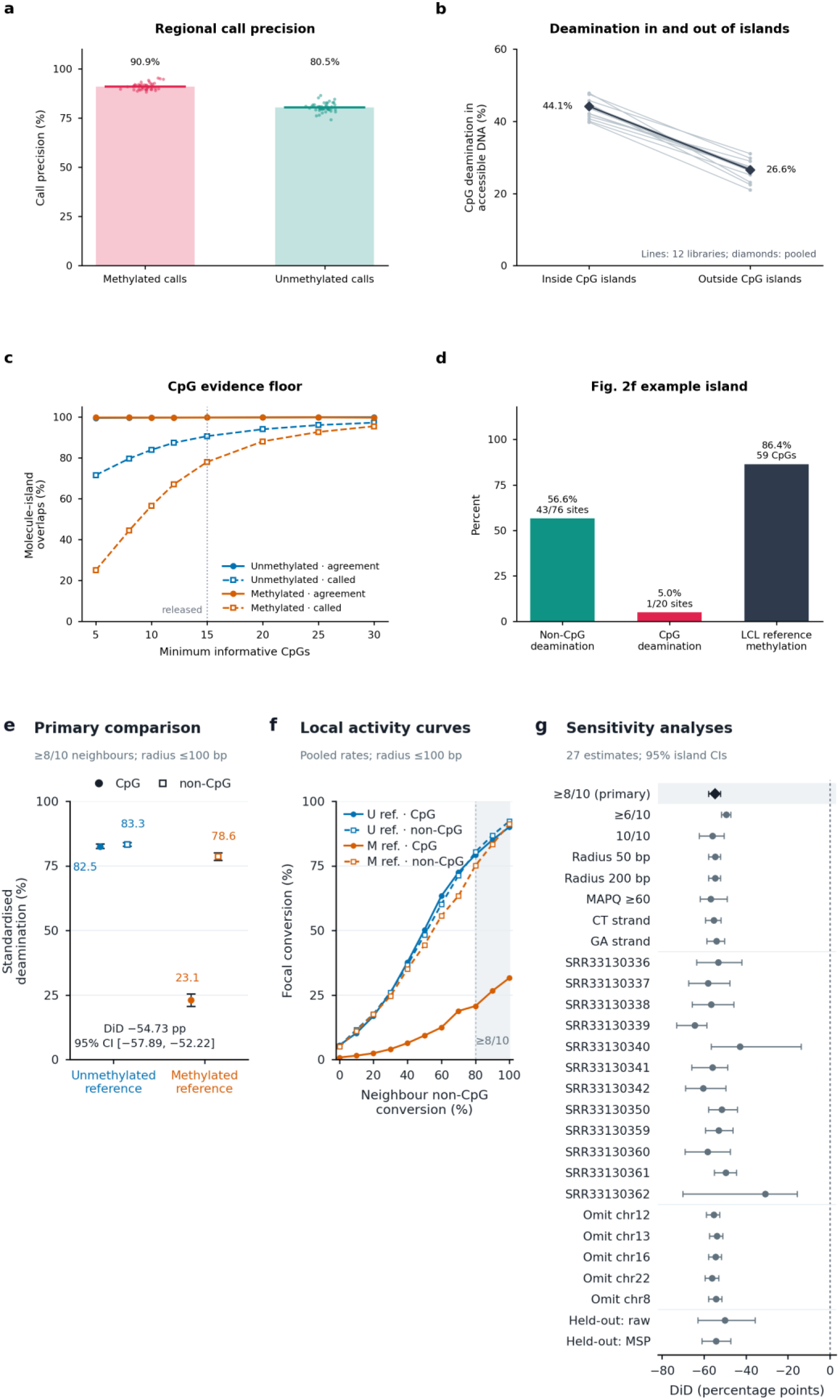
Complete-CpG-island methylation inference. **a, Regional call precision.** Methylated- and unmethylated-call precision in the 1-kb regional validation against the HG002 LCL reference, shown on a full 0–100% scale. Points are 46 haplotype-contig estimates; bars are means weighted by the number of truth windows (90.9% and 80.5%, respectively). **b, Deamination inside and outside islands.** CpG deamination in accessible DNA inside and outside inferred CpG islands across the validation libraries; gray lines, individual libraries; diamonds, pooled. **c, CpG evidence floor. Agreement with the HG002 LCL** reference among confident calls (solid) and fraction of evidence-eligible molecule–island overlaps called confidently (dashed) for unmethylated- and methylated-reference islands on validation chromosomes, as the minimum number of informative CpGs varies; posterior threshold 0.99 and at least 10 non-CpG observations. The dotted line marks the 15-CpG floor. **d,** Fig. 2f **example island.** Deamination of non-CpG (43/76) and CpG (1/20) sites in the Fig. 2f island and its matched HG002 LCL long-read methylation (86.4% across 59 CpGs). **e, CpG-specific loss at matched local accessibility.** Standardized CpG and non-CpG deamination in methylated- and unmethylated-reference islands for focal sites with at least eight of their ten nearest non-CpG neighbors within 100 bp converted. Difference-in-differences −54.73 percentage points (95% island-bootstrap interval −57.89 to −52.22); 179,332 matched site observations across 1,172 islands and 12 libraries. **f, Local activity curves.** Focal conversion against neighbor non-CpG conversion. Non-CpG curves overlap between reference states at every accessibility level, whereas CpG conversion is reduced only in methylated-reference islands. Shaded region, primary ≥8/10 tier; U, unmethylated reference; M, methylated reference. **g, Sensitivity analyses.** Difference-in-differences with 95% island-bootstrap intervals for alternative neighbor requirements, radii and mapping quality, each strand, each library, each omitted test chromosome and two held-out analyses in which island state was assigned from 80% of CpGs and conversion measured at the withheld 20%, using either initial MSPs or raw non-CpG neighbors alone.

**Supplementary Figure 8.**
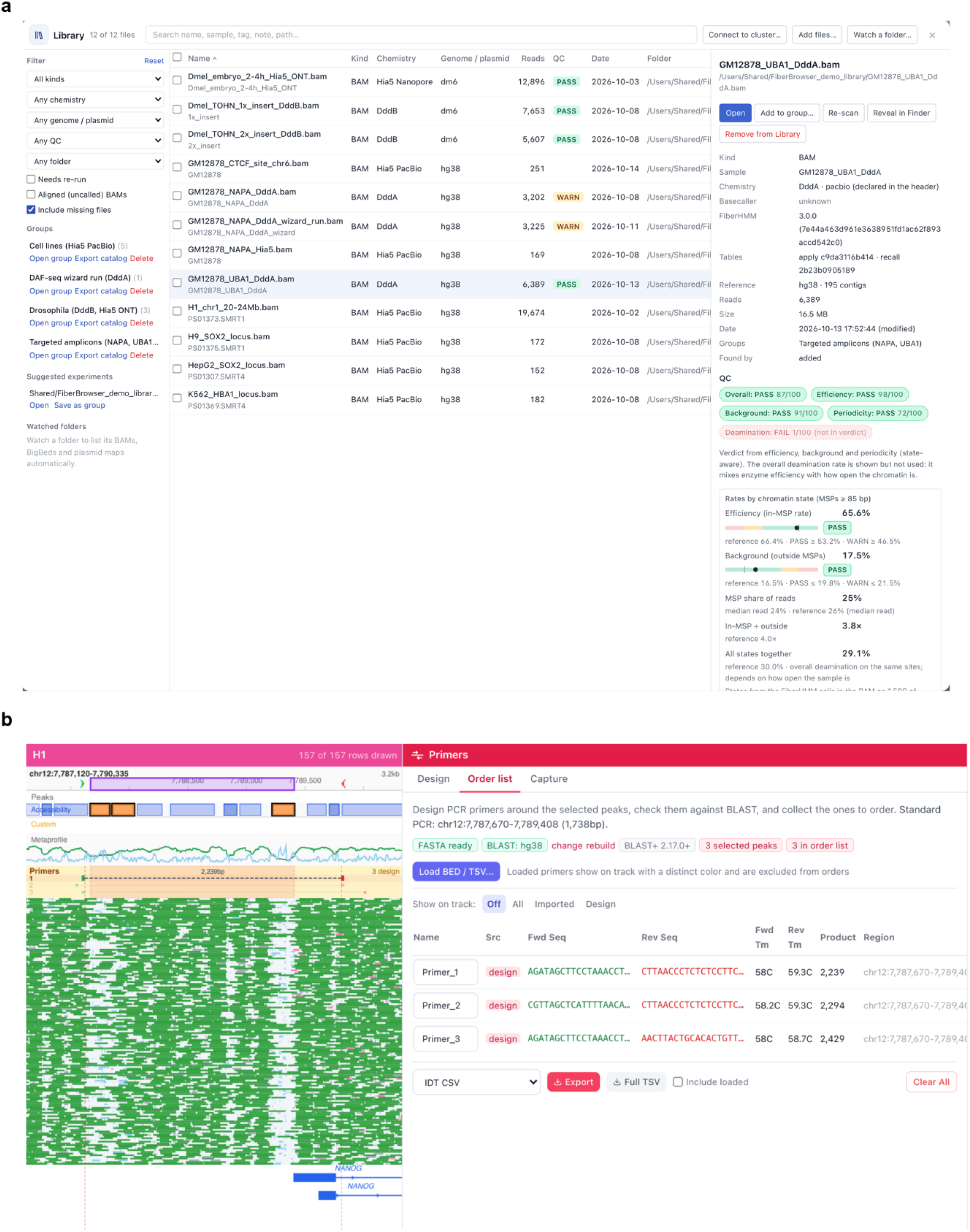
FiberBrowser data library and primer design. **a, Data library.** Twelve datasets from this study (Hia5 PacBio, Hia5 ONT, DddB and SsDddA, labeled DddA in the interface) organized into user-defined groups. Chemistry, genome and read counts are read from the BAM headers, and quality control (QC) compares each dataset with a chemistry-matched control; QC is shown only where such a control exists. The selected SsDddA amplicon dataset passes the efficiency, background and periodicity checks; its overall deamination rate is shown but excluded from the state-aware verdict because it mixes accessibility with labeling efficiency. b, Primer design at *NANOG* in H1. Candidate amplicons are displayed above the molecular view, alongside primer sequences, melting temperatures and product lengths in the exportable order list. The displayed primers have not been experimentally validated.

**Supplementary Figure 9.**
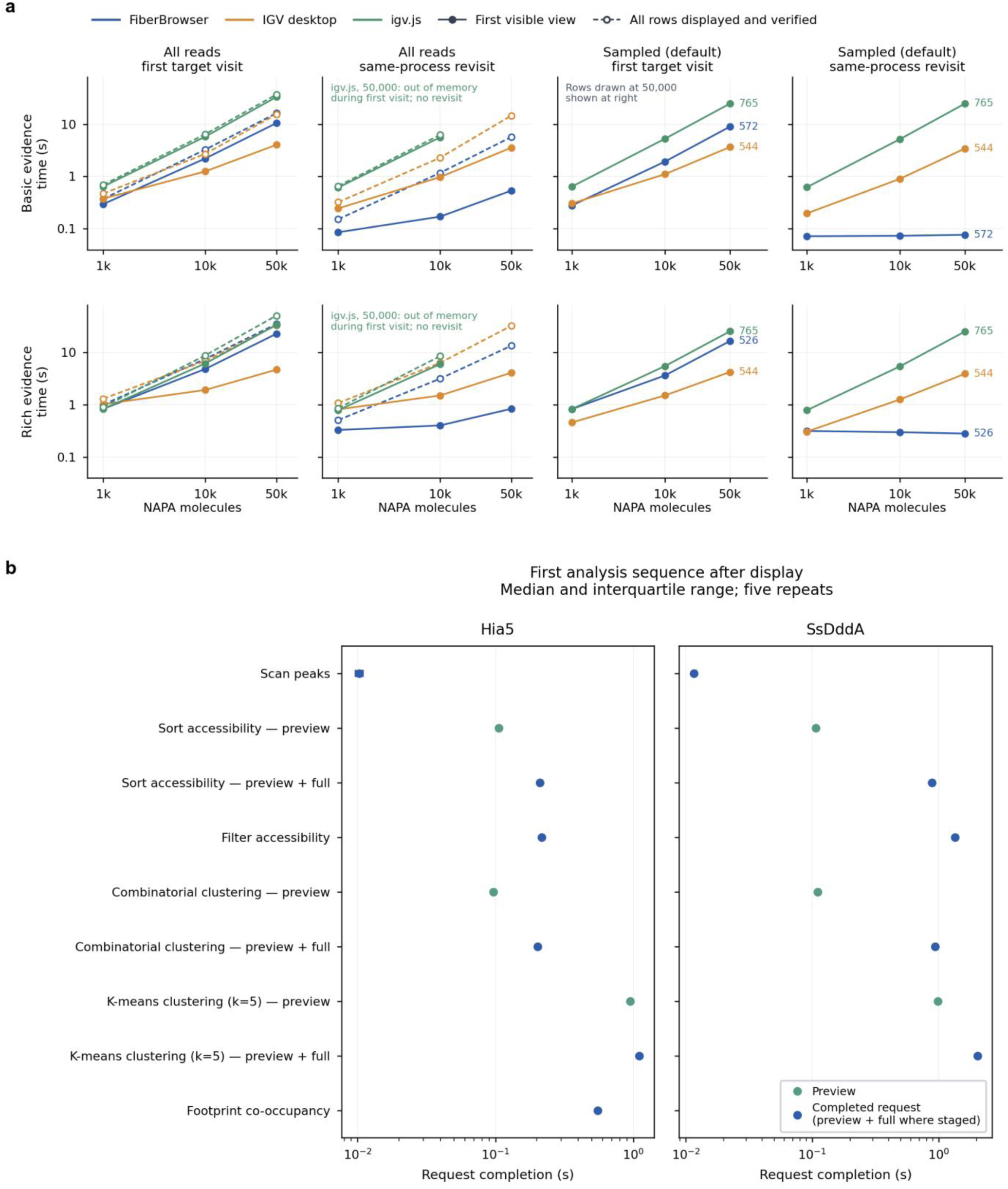
FiberBrowser display and analysis performance. **a, Display performance.** FiberBrowser, IGV desktop and igv.js at 1,000–50,000 NAPA molecules, with all reads displayed (left) or with each tool’s default sampling (right), on a first target visit or a same-process revisit. Solid lines, time to the first visible view (first bounded viewport image written). Dashed lines, time until every read has been displayed and verified across bounded image tiles (all-read trials only). Numbers at right give the rows drawn at 50,000 molecules with sampling. Points show the median and bars the interquartile range of five trials. Basic mode omits additional molecular evidence; rich mode displays FiberBrowser molecular annotations or IGV mismatch evidence. These are distinct annotation workloads. igv.js ran out of memory during the first all-read visit at 50,000 molecules, so it has no revisit. **b,** First sequence after verified display; median and interquartile range of five trials. Preview and preview-plus-full completion are separate measurements. The repeated-sequence plot and per-stage source tables are supplied separately. Teal, preview; blue, completed request (preview plus full refinement where staged).

**Supplementary Figure 10.**
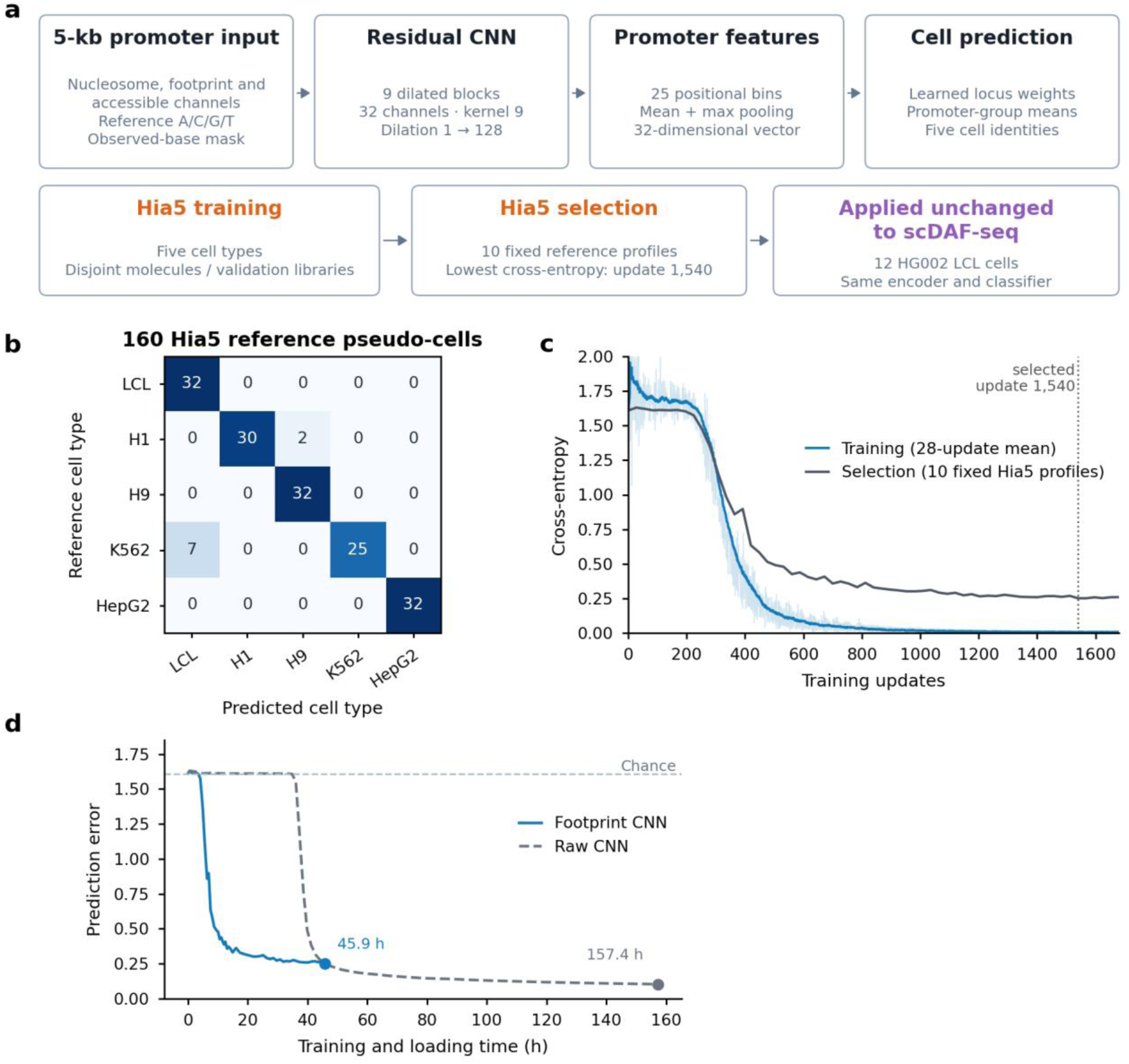
Promoter CNN architecture, training and selection. **a, Promoter CNN.** Input channels (nucleosome, footprint and accessible calls; reference sequence), dilated convolutional encoder, promoter aggregation and five-class prediction. Training and checkpoint selection use Hia5 only; HG002 scDAF-seq predictions use the frozen model. **b, Reference confusion matrix.** Counts among 160 fixed Hia5 reference pseudo-cells, 32 per cell type. Rows are reference identities and columns predictions; color denotes row percentage. All nine errors occur at 8% promoter retention; seven are K562 pseudo-cells assigned to LCL at near-tie scores (mean K562–LCL logit −0.13), whereas the learned class prior is uniform. **c, CNN optimization and selection.** Individual training cross-entropies (pale), their 28-update rolling mean (blue) and cross-entropy on ten fixed selection profiles (dark gray). The dotted line marks selected update 1,540. **d, CNN training histories.** Prediction error (validation cross-entropy) of the footprint and raw-input CNNs against cumulative logged training and loading time. Points mark the reference-selected checkpoints (footprint, update 1,540 at 45.9 h; raw, update 3,780 at 157.4 h); dashed guide, five-class chance. Times are to the selected checkpoints.

**Supplementary Figure 11.**
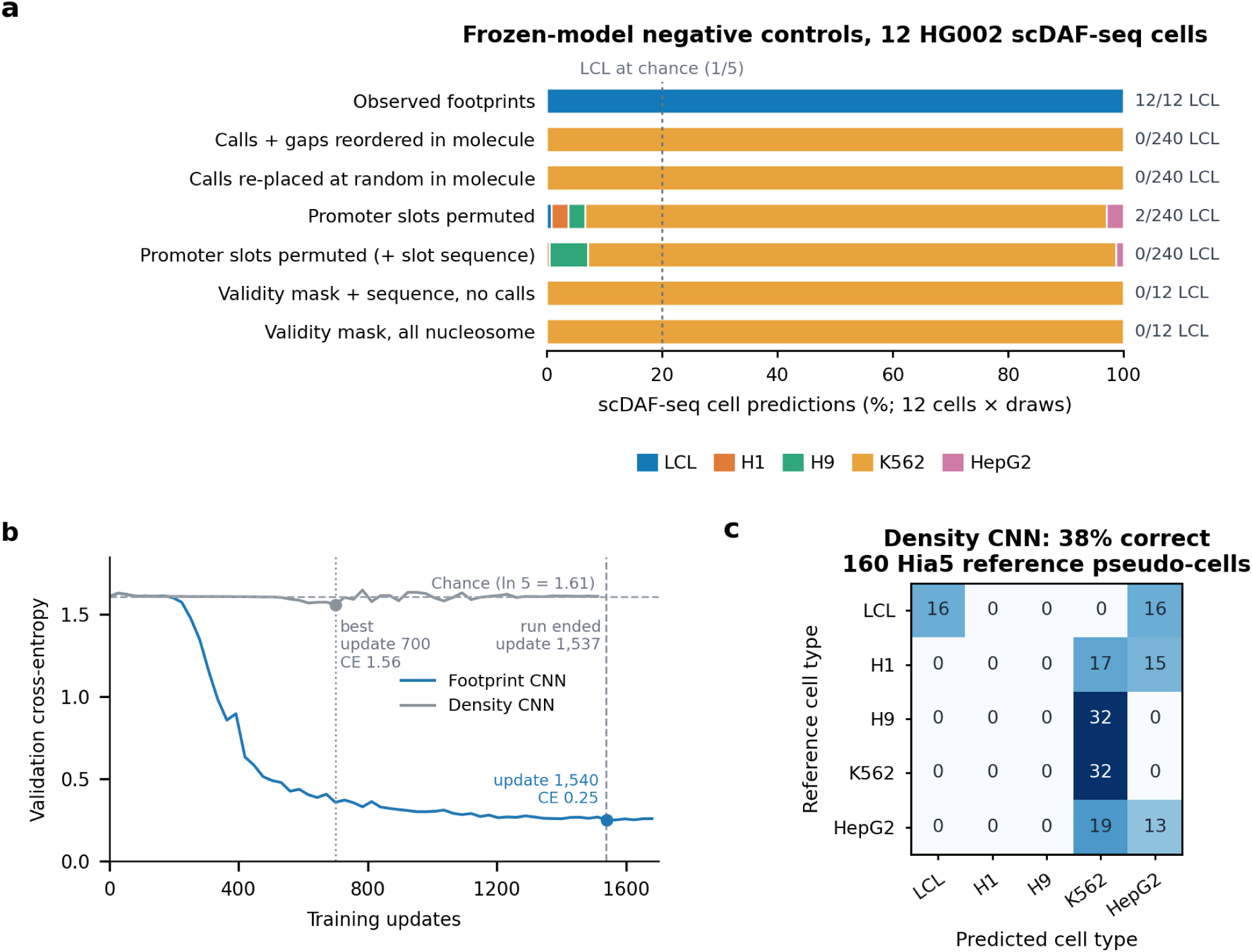
Promoter CNN controls. **a, Frozen-model negative controls on the 12 HG002 scDAF-seq cells.** Predicted-class composition for observed footprints and for inputs with calls shuffled within molecules, promoter slots permuted, or only the observation-validity mask supplied (20 draws per stochastic control; 240 predictions each). Labels give LCL assignments; the dashed line marks five-class chance. In a separate mixture control, pseudo-cells built from one molecule per promoter drawn from a randomly chosen reference line were weakly assigned to LCL at full coverage (20 of 20; LCL probability 0.28–0.36), far below the confidence of the scDAF-seq cells (0.77–0.98). **b, Methylation-density CNN training.** A CNN with the footprint CNN’s architecture and training recipe was given the fraction of modified informative sites in a 51-bp running window instead of FiberHMM calls, trained on Hia5 only and selected on the same ten fixed Hia5 validation profiles. Validation cross-entropy against training updates: the methylation-density CNN stayed near five-class chance (ln 5 = 1.61; dashed line), with its best checkpoint at update 700 (1.56; dotted line), whereas the footprint CNN reached 0.25 at update 1,540. Over the same updates its training loss fell from 1.7 to 0.7, so the density model fitted its training libraries without generalizing to held-out libraries. Training ran without a stopping rule to update 1,537 (dashed vertical line) with no further improvement. **c, Methylation-density CNN reference confusion.** Counts among the 160 Hia5 reference pseudo-cells, as in Supplementary Fig. 10b (38% correct, 61 of 160). The methylation-density CNN never predicted H1 or H9 and assigned most pseudo-cells to K562 (100 of 160) or HepG2 (44 of 160).

**Supplementary Table 1.** Molecular Annotation encoding written by FiberHMM and read by the FiberBrowser interface. The Molecular Annotation specification developed by Mitchell R. Vollger supplies the shared interval representation. Coordinates refer to the original molecule, including reverse-aligned reads.

| Tag or layer | Stored content | Interpretation |
|---|---|---|
| MA:Z | Original molecule length; 1-based start/length intervals; type, strand and quality declarations | Alignment supplies genomic projection. |
| AQ:B:C | Declared quality bytes in MA interval order | Q is a linear 0–255 byte; P is Phred-scaled. Meaning depends on the declared field. |
| AN:Z | Optional names aligned to MA intervals | Optional interval names; native annotations are unchanged. |
| nuc.Q or nuc.QQQ | Nucleosome quality; optional left/right edge qualities | Native nucleosome layer. |
| msp. | Accessible-patch intervals | Final coordinates are also written to as/al for fibertools/FIRE. |
| tf.QQQ | TQ, left-edge quality, right-edge quality | TQ = min(255, round(10 × LLR)); it is not a posterior probability. |
| ddda_mcg. | Confident methylated-island interval | Whole-island molecule-specific annotation; not per-CpG methylation. |
| ddda_ucg. | Confident unmethylated-island interval | High-precision allowlist for CpG footprint evidence in CpG-aware SsDddA footprint recall. |

See Online Methods, Molecular Annotation encoding and FIRE interoperability, and the cited MA specification.

**Supplementary Table 2.** Motif footprint AUROCs against ChIP-seq.

| Factor | Cell line | ft footprint | FiberHMM, ≤74 bp [− ft] | FiberHMM, ChIP-calibrated | Window (bp) | Calibrated − ft |
|---|---|---|---|---|---|---|
| CTCF | K562 | 0.838 (0.799–0.874) | 0.855 (0.818–0.889) [+0.017 (0.006–0.029)] | 0.870 (0.836–0.902) | 20–60 | 0.031 (0.017–0.045) |
|  | GM12878 | 0.807 (0.767–0.846) | 0.836 (0.799–0.872) [+0.030 (0.017–0.042)] | 0.842 (0.806–0.877) | 25–55 | 0.035 (0.015–0.056) |
|  | H1 | 0.873 (0.840–0.905) | 0.885 (0.853–0.915) [+0.012 (0.003–0.020)] | 0.896 (0.865–0.923) | 20–55 | 0.023 (0.009–0.036) |
|  | H9 | 0.842 (0.807–0.876) | 0.854 (0.820–0.886) [+0.013 (0.004–0.021)] | 0.873 (0.842–0.902) | 25–55 | 0.031 (0.014–0.049) |
|  | HepG2 | 0.864 (0.828–0.895) | 0.888 (0.855–0.916) [+0.024 (0.014–0.034)] | 0.900 (0.870–0.927) | 25–55 | 0.036 (0.018–0.053) |
| NRF1 | K562 | 0.725 (0.660–0.787) | 0.784 (0.722–0.842) [+0.058 (0.024–0.099)] | 0.817 (0.761–0.871) | 20–55 | 0.092 (0.047–0.143) |
|  | GM12878 | 0.785 (0.724–0.840) | 0.817 (0.758–0.867) [+0.031 (0.006–0.055)] | 0.835 (0.777–0.883) | 15–55 | 0.050 (0.018–0.082) |
|  | H1 | 0.795 (0.740–0.849) | 0.826 (0.775–0.878) [+0.030 (0.007–0.053)] | 0.853 (0.804–0.900) | 15–55 | 0.057 (0.028–0.088) |
|  | HepG2 | 0.723 (0.658–0.786) | 0.762 (0.701–0.823) [+0.039 (0.021–0.057)] | 0.780 (0.722–0.837) | 15–55 | 0.057 (0.033–0.082) |
| REST | K562 | 0.650 (0.570–0.728) | 0.734 (0.659–0.803) [+0.084 (0.053–0.115)] | 0.828 (0.767–0.883) | 10–45 | 0.178 (0.124–0.238) |
|  | GM12878 | 0.642 (0.562–0.720) | 0.713 (0.638–0.785) [+0.071 (0.033–0.110)] | 0.757 (0.688–0.822) | 15–45 | 0.116 (0.062–0.171) |
|  | H1 | 0.885 (0.829–0.934) | 0.922 (0.878–0.958) [+0.038 (0.019–0.058)] | 0.944 (0.906–0.976) | 0–40 | 0.059 (0.013–0.103) |
|  | HepG2 | 0.736 (0.650–0.815) | 0.800 (0.727–0.866) [+0.064 (0.039–0.092)] | 0.817 (0.750–0.878) | 10–35 | 0.081 (0.029–0.132) |
| USF1 | K562 | 0.701 (0.638–0.762) | 0.710 (0.646–0.769) [+0.008 (−0.019–0.036)] | 0.701 (0.639–0.760) | 15–65 | −0.000 (−0.042–0.043) |
|  | GM12878 | 0.689 (0.622–0.758) | 0.703 (0.636–0.766) [+0.014 (−0.011–0.040)] | 0.713 (0.645–0.777) | 10–65 | 0.024 (−0.015–0.064) |
|  | H1 | 0.715 (0.653–0.777) | 0.725 (0.665–0.784) [+0.010 (−0.018–0.037)] | 0.733 (0.672–0.788) | 15–65 | 0.017 (−0.025–0.058) |
|  | HepG2 | 0.678 (0.612–0.740) | 0.672 (0.608–0.734) [−0.005 (−0.032–0.020)] | 0.691 (0.627–0.751) | 10–65 | 0.014 (−0.021–0.049) |

AUROC (95% CI) of per-motif bound fractions against cell-matched ChIP-seq. All callers use the same MSP-eligible molecule–motif observations. FiberHMM, ≤74 bp: fiberhmm-footprint with the label-free 74-bp maximum width. ChIP-calibrated: the window learned on the other cell lines of the same factor (leave-one-cell-line-out) and applied to this cell line; Window, the learned [minimum, maximum] conservative call width. Bracketed values and Calibrated − ft, paired differences from ft footprint with 95% CI. ≤74 bp − ft intervals exclude zero for every CTCF, NRF1 and REST cell line and include zero for every USF1 cell line. Confidence intervals use paired genomic-block resampling (Supplementary Note 1). Values are rounded to three decimal places; differences were calculated before rounding.

**Supplementary Table 3.** CTCF motif footprint AUROCs across footprint thresholds.

| Cell line | LLR | FiberHMM ≤74 bp AUROC (95% CI) | FiberHMM − ft (paired 95% CI) | Bound fraction, ChIP+ / ChIP− |
|---|---|---|---|---|
| K562 | 2 | 0.850 (0.813–0.883) | 0.010 (−0.002–0.021) | 0.421 / 0.088 |
|  | 3 | 0.851 (0.814–0.885) | 0.011 (−0.001–0.022) | 0.420 / 0.086 |
|  | 4 | 0.853 (0.817–0.886) | 0.014 (0.001–0.025) | 0.417 / 0.082 |
|  | 5 | 0.854 (0.818–0.887) | 0.014 (0.001–0.027) | 0.411 / 0.077 |
|  | 6 | 0.854 (0.819–0.888) | 0.015 (0.001–0.028) | 0.408 / 0.076 |
|  | 7 | 0.855 (0.819–0.888) | 0.015 (−0.001–0.030) | 0.405 / 0.073 |
|  | 8 | 0.855 (0.818–0.889) | 0.015 (−0.001–0.030) | 0.403 / 0.070 |
| GM12878 | 2 | 0.833 (0.795–0.868) | 0.025 (0.011–0.039) | 0.372 / 0.065 |
|  | 3 | 0.835 (0.797–0.870) | 0.027 (0.012–0.041) | 0.371 / 0.062 |
|  | 4 | 0.835 (0.798–0.870) | 0.027 (0.013–0.042) | 0.368 / 0.060 |
|  | 5 | 0.833 (0.796–0.868) | 0.025 (0.009–0.040) | 0.362 / 0.057 |
|  | 6 | 0.838 (0.800–0.873) | 0.030 (0.013–0.046) | 0.360 / 0.053 |
|  | 7 | 0.839 (0.802–0.874) | 0.031 (0.014–0.048) | 0.358 / 0.050 |
|  | 8 | 0.840 (0.803–0.875) | 0.032 (0.014–0.050) | 0.356 / 0.047 |
| H1 | 2 | 0.880 (0.847–0.911) | 0.005 (−0.005–0.014) | 0.460 / 0.077 |
|  | 3 | 0.880 (0.847–0.912) | 0.006 (−0.005–0.015) | 0.459 / 0.075 |
|  | 4 | 0.883 (0.851–0.914) | 0.009 (−0.002–0.018) | 0.455 / 0.072 |
|  | 5 | 0.884 (0.852–0.914) | 0.010 (−0.002–0.020) | 0.449 / 0.070 |
|  | 6 | 0.885 (0.853–0.915) | 0.011 (−0.002–0.021) | 0.447 / 0.067 |
|  | 7 | 0.884 (0.852–0.914) | 0.010 (−0.005–0.022) | 0.444 / 0.065 |
|  | 8 | 0.883 (0.851–0.913) | 0.009 (−0.007–0.022) | 0.442 / 0.062 |
| H9 | 2 | 0.851 (0.816–0.884) | 0.009 (−0.001–0.018) | 0.416 / 0.067 |
|  | 3 | 0.851 (0.817–0.884) | 0.009 (−0.001–0.019) | 0.416 / 0.066 |
|  | 4 | 0.851 (0.816–0.883) | 0.009 (−0.002–0.019) | 0.412 / 0.063 |
|  | 5 | 0.853 (0.819–0.885) | 0.011 (−0.000–0.021) | 0.406 / 0.061 |
|  | 6 | 0.853 (0.819–0.885) | 0.011 (−0.001–0.023) | 0.405 / 0.057 |
|  | 7 | 0.852 (0.817–0.884) | 0.010 (−0.004–0.023) | 0.403 / 0.055 |
|  | 8 | 0.852 (0.818–0.885) | 0.010 (−0.004–0.024) | 0.401 / 0.052 |
| HepG2 | 2 | 0.877 (0.843–0.906) | 0.013 (0.001–0.023) | 0.443 / 0.070 |
|  | 3 | 0.878 (0.844–0.907) | 0.013 (0.001–0.024) | 0.442 / 0.067 |
|  | 4 | 0.883 (0.850–0.911) | 0.019 (0.006–0.031) | 0.439 / 0.063 |
|  | 5 | 0.887 (0.855–0.916) | 0.023 (0.010–0.035) | 0.433 / 0.060 |
|  | 6 | 0.888 (0.856–0.917) | 0.024 (0.010–0.037) | 0.431 / 0.058 |
|  | 7 | 0.888 (0.854–0.916) | 0.023 (0.009–0.038) | 0.429 / 0.055 |
|  | 8 | 0.889 (0.855–0.917) | 0.025 (0.010–0.039) | 0.427 / 0.052 |

CTCF calls were re-decoded at each threshold and scored by fiberhmm-footprint with the label-free 74-bp maximum width, with the LLR 5 eligible molecule–motif observations as a fixed denominator. ft footprint AUROCs do not depend on the FiberHMM threshold and are given in Supplementary Table 2. Bound fractions are pooled over eligible observations at ChIP-supported and unsupported motifs. Because the decoders were re-run at each threshold, LLR 5 values differ slightly from Supplementary Table 2. Confidence intervals use paired genomic-block resampling (Supplementary Note 1).

**Supplementary Table 4.** Application failures in the display benchmark.

| Tool / depth | Mode / evidence | Failed operation | Failure kind | Trials |
|---|---|---|---|---|
| igv.js / 50000 | all / basic | pan by half-width (first visit) | JavaScript heap out of memory | 5 |
| igv.js / 50000 | all / rich | pan by half-width (first visit) | JavaScript heap out of memory | 5 |

**Supplementary Table 5.** All-read display times: first visible view and all rows displayed and verified.

| Molecules | Tool / evidence | Visit | First visible view, median [IQR], s | All rows displayed and verified, median [IQR], s |
|---|---|---|---|---|
| 1000 | FiberBrowser / basic | cold | 0.297 [0.294–0.300] | 0.373 [0.370–0.376] |
| 10000 | FiberBrowser / basic | cold | 2.203 [2.203–2.210] | 3.253 [3.243–3.261] |
| 50000 | FiberBrowser / basic | cold | 10.444 [10.336–10.488] | 16.109 [15.733–16.259] |
| 1000 | IGV desktop / basic | cold | 0.378 [0.378–0.390] | 0.472 [0.467–0.478] |
| 10000 | IGV desktop / basic | cold | 1.248 [1.234–1.256] | 2.695 [2.693–2.724] |
| 50000 | IGV desktop / basic | cold | 4.019 [4.004–4.025] | 15.352 [15.186–15.355] |
| 1000 | igv.js / basic | cold | 0.643 [0.636–0.645] | 0.695 [0.688–0.696] |
| 10000 | igv.js / basic | cold | 5.744 [5.738–5.769] | 6.410 [6.398–6.430] |
| 50000 | igv.js / basic | cold | 32.964 [32.874–33.033] | 36.727 [36.635–36.789] |
| 1000 | FiberBrowser / basic | warm | 0.086 [0.078–0.086] | 0.153 [0.144–0.153] |
| 10000 | FiberBrowser / basic | warm | 0.171 [0.169–0.176] | 1.171 [1.170–1.172] |
| 50000 | FiberBrowser / basic | warm | 0.538 [0.537–0.543] | 5.679 [5.653–5.711] |
| 1000 | IGV desktop / basic | warm | 0.245 [0.245–0.249] | 0.323 [0.319–0.328] |
| 10000 | IGV desktop / basic | warm | 0.972 [0.961–0.981] | 2.280 [2.275–2.297] |
| 50000 | IGV desktop / basic | warm | 3.553 [3.544–3.557] | 14.536 [14.477–14.565] |
| 1000 | igv.js / basic | warm | 0.614 [0.614–0.615] | 0.658 [0.658–0.659] |
| 10000 | igv.js / basic | warm | 5.585 [5.570–5.591] | 6.230 [6.205–6.240] |
| 50000 | igv.js / basic | warm | Unreached / failed (0/5 completed) | Unreached / failed (0/5 completed) |
| 1000 | FiberBrowser / rich | cold | 0.830 [0.829–0.833] | 0.999 [0.995–1.008] |
| 10000 | FiberBrowser / rich | cold | 4.796 [4.789–4.845] | 7.457 [7.445–7.462] |
| 50000 | FiberBrowser / rich | cold | 22.543 [22.373–22.598] | 34.652 [34.605–34.776] |
| 1000 | IGV desktop / rich | cold | 1.022 [1.020–1.028] | 1.297 [1.296–1.302] |
| 10000 | IGV desktop / rich | cold | 1.904 [1.896–1.913] | 6.838 [6.760–6.852] |
| 50000 | IGV desktop / rich | cold | 4.652 [4.640–4.685] | 32.756 [32.722–32.896] |
| 1000 | igv.js / rich | cold | 0.828 [0.827–0.834] | 0.905 [0.902–0.912] |
| 10000 | igv.js / rich | cold | 6.106 [6.104–6.107] | 8.699 [8.686–8.712] |
| 50000 | igv.js / rich | cold | 33.406 [33.333–33.428] | 50.217 [50.111–50.231] |
| 1000 | FiberBrowser / rich | warm | 0.328 [0.328–0.342] | 0.507 [0.505–0.507] |
| 10000 | FiberBrowser / rich | warm | 0.401 [0.401–0.401] | 3.134 [3.079–3.136] |
| 50000 | FiberBrowser / rich | warm | 0.834 [0.825–0.839] | 13.352 [13.271–13.357] |
| 1000 | IGV desktop / rich | warm | 0.812 [0.798–0.820] | 1.073 [1.054–1.087] |
| 10000 | IGV desktop / rich | warm | 1.496 [1.489–1.514] | 6.293 [6.283–6.404] |
| 50000 | IGV desktop / rich | warm | 4.071 [4.029–4.079] | 32.011 [31.919–32.072] |
| 1000 | igv.js / rich | warm | 0.785 [0.782–0.797] | 0.857 [0.854–0.868] |
| 10000 | igv.js / rich | warm | 5.937 [5.924–5.985] | 8.436 [8.406–8.462] |
| 50000 | igv.js / rich | warm | Unreached / failed (0/5 completed) | Unreached / failed (0/5 completed) |

**Supplementary Table 5.**
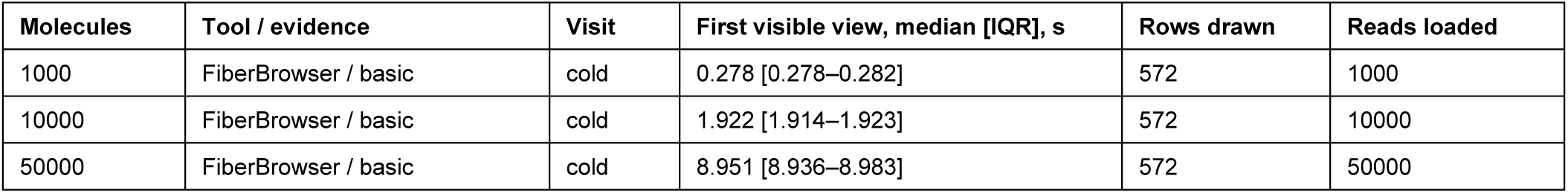
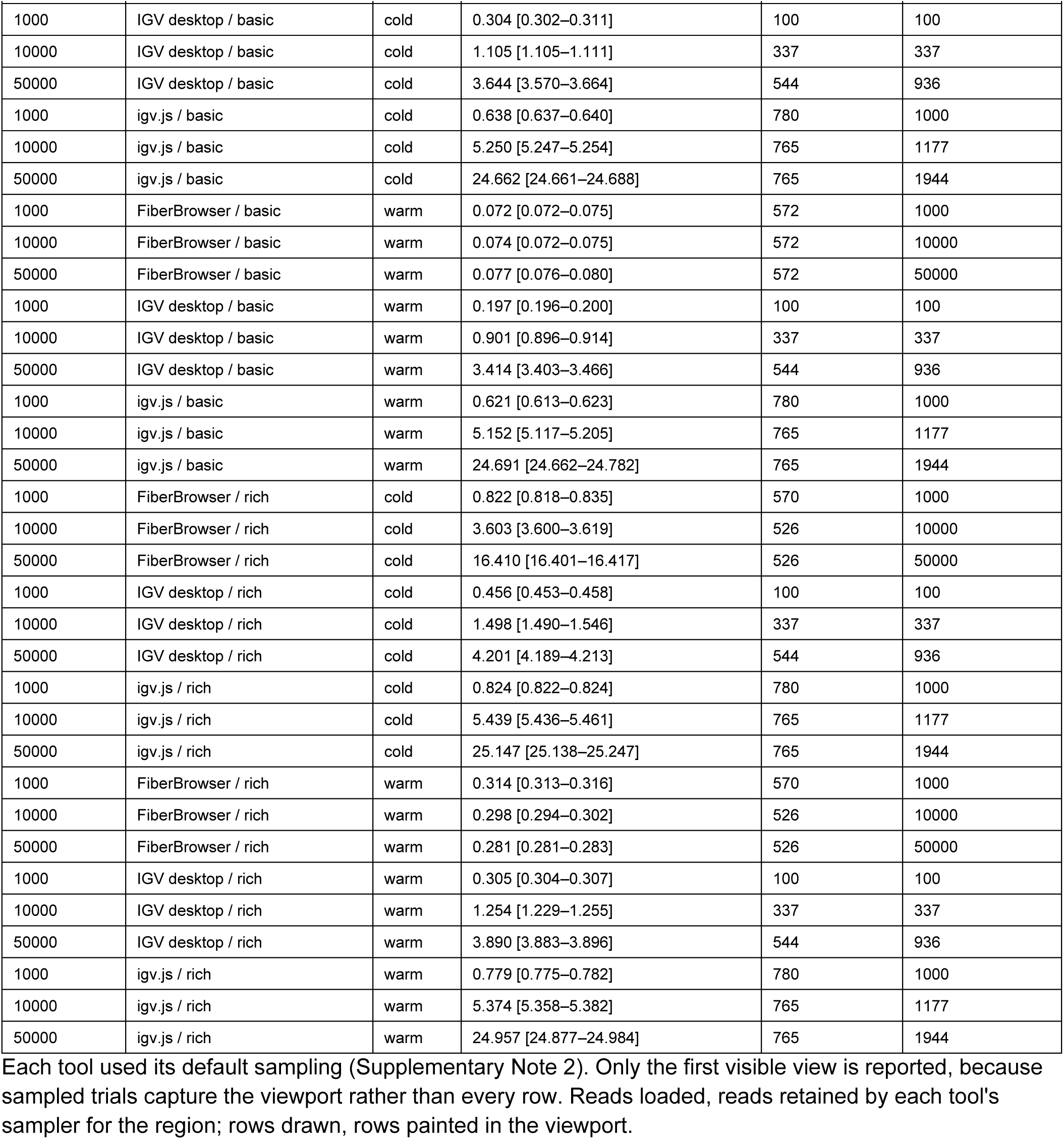
Default-sampling display times: first visible view, rows drawn and reads loaded.

**Supplementary Table 6.**
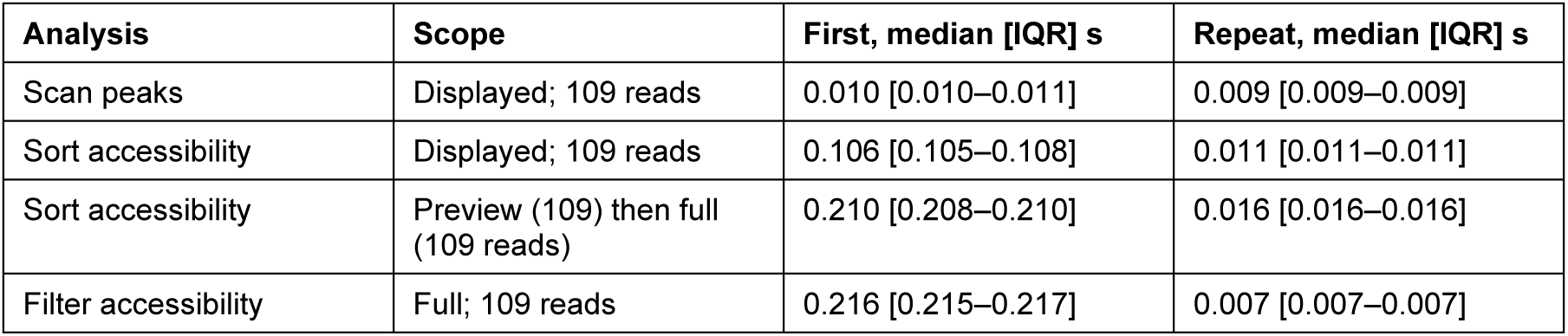
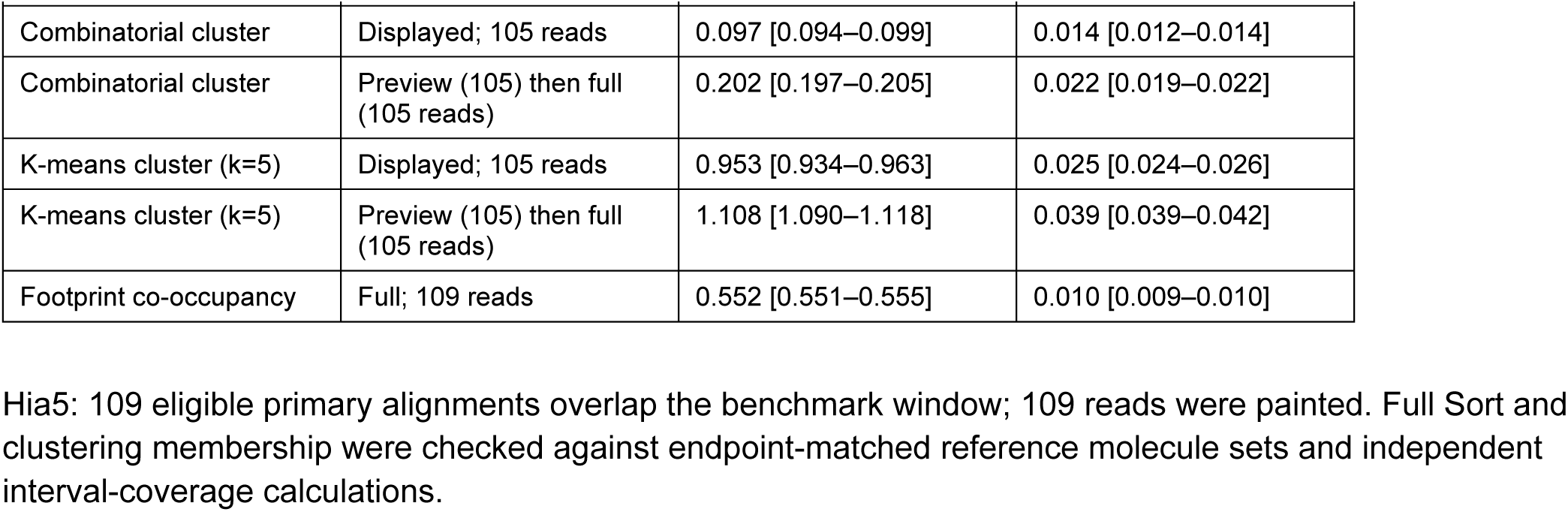
FiberBrowser analysis times for Hia5.

| **Analysis** | **Scope** | **First, median [IQR] s** | **Repeat, median [IQR] s** |
|---|---|---|---|
| Scan peaks | Displayed; 109 reads | 0.010 [0.010–0.011] | 0.009 [0.009–0.009] |
| Sort accessibility | Displayed; 109 reads | 0.106 [0.105–0.108] | 0.011 [0.011–0.011] |
| Sort accessibility | Preview (109) then full (109 reads) | 0.210 [0.208–0.210] | 0.016 [0.016–0.016] |
| Filter accessibility | Full; 109 reads | 0.216 [0.215–0.217] | 0.007 [0.007–0.007] |
| Combinatorial cluster | Displayed; 105 reads | 0.097 [0.094–0.099] | 0.014 [0.012–0.014] |
| Combinatorial cluster | Preview (105) then full (105 reads) | 0.202 [0.197–0.205] | 0.022 [0.019–0.022] |
| K-means cluster (k=5) | Displayed; 105 reads | 0.953 [0.934–0.963] | 0.025 [0.024–0.026] |
| K-means cluster (k=5) | Preview (105) then full (105 reads) | 1.108 [1.090–1.118] | 0.039 [0.039–0.042] |
| Footprint co-occupancy | Full; 109 reads | 0.552 [0.551–0.555] | 0.010 [0.009–0.010] |

Hia5: 109 eligible primary alignments overlap the benchmark window; 109 reads were painted. Full Sort and clustering membership were checked against endpoint-matched reference molecule sets and independent interval-coverage calculations.

**Supplementary Table 7.** FiberBrowser analysis times for SsDddA.

| Analysis | Scope | First, median [IQR] s | Repeat, median [IQR] s |
|---|---|---|---|
| Scan peaks | Displayed; 190 reads | 0.012 [0.011–0.012] | 0.008 [0.008–0.009] |
| Sort accessibility | Displayed; 190 reads | 0.107 [0.105–0.107] | 0.016 [0.015–0.016] |
| Sort accessibility | Preview (190) then full (2,976 reads) | 0.883 [0.881–0.887] | 0.030 [0.030–0.030] |
| Filter accessibility | Full; 2,976 reads | 1.341 [1.341–1.347] | 0.023 [0.022–0.026] |
| Combinatorial cluster | Displayed; 190 reads | 0.110 [0.110–0.115] | 0.019 [0.019–0.021] |
| Combinatorial cluster | Preview (190) then full (2,966 reads) | 0.939 [0.935–0.940] | 0.083 [0.081–0.084] |
| K-means cluster (k=5) | Displayed; 190 reads | 0.982 [0.970–0.988] | 0.028 [0.028–0.030] |
| K-means cluster (k=5) | Preview (190) then full (2,965 reads) | 2.030 [2.023–2.032] | 0.092 [0.089–0.093] |
| Footprint co-occupancy | Full; 2,975 reads | 1.430 [1.427–1.433] | 0.026 [0.024–0.029] |

SsDddA: 2,976 eligible primary alignments overlap the benchmark window; 190 reads were painted. Full Sort and clustering membership were checked against endpoint-matched reference molecule sets and independent interval-coverage calculations.

**Supplementary Table 8.**
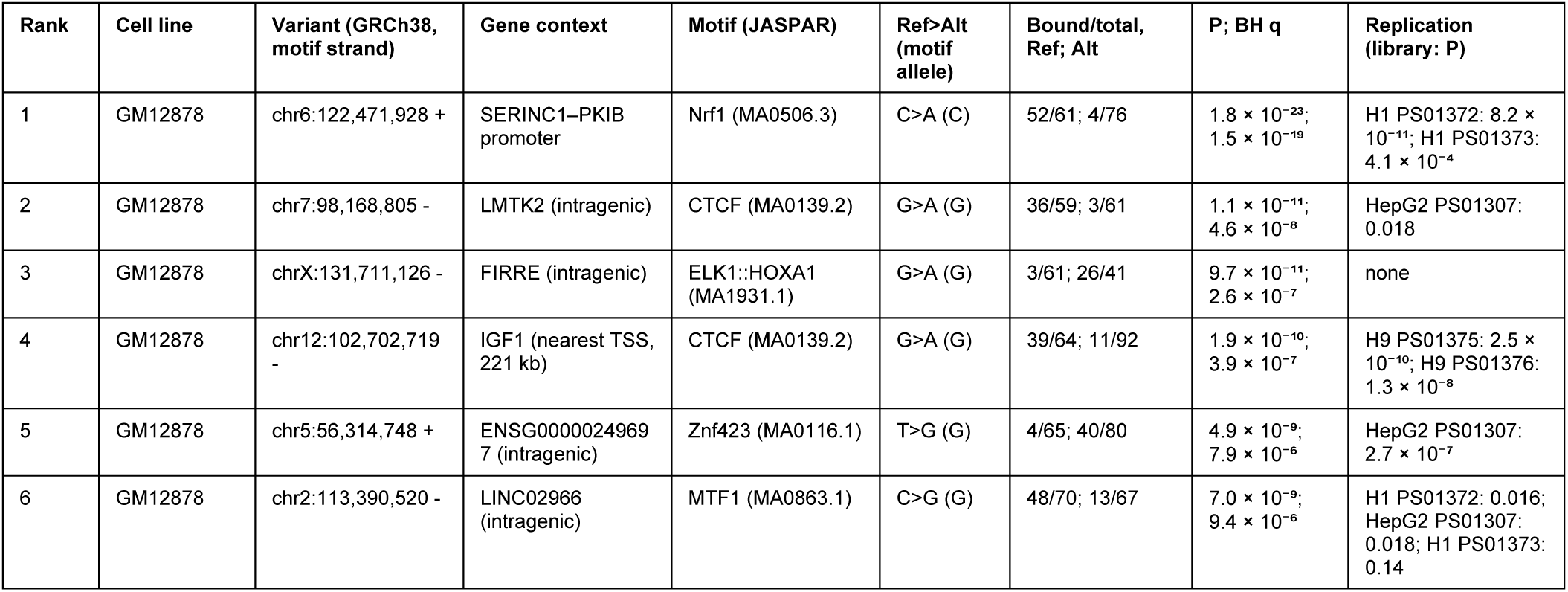
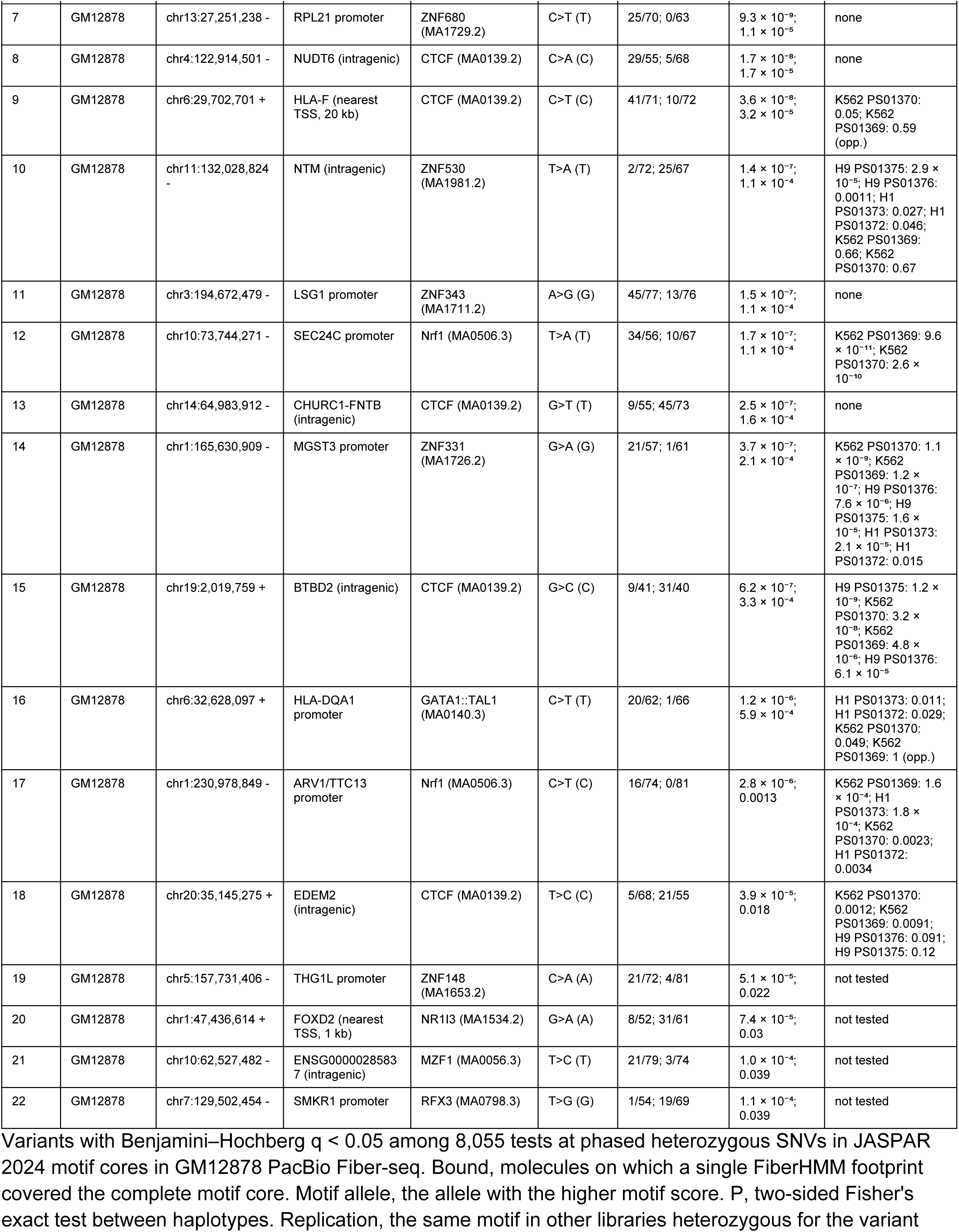
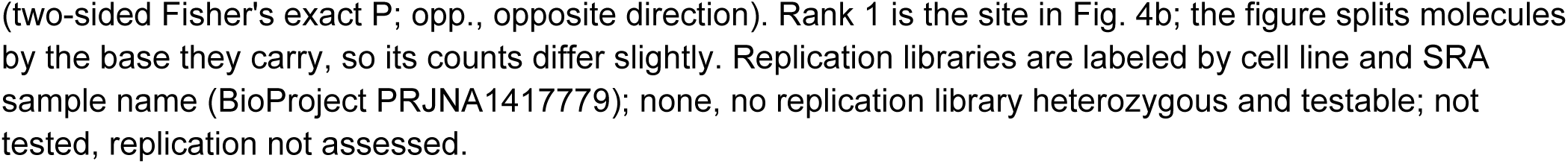
Allelic footprints at heterozygous motif variants.

**Supplementary Table 9.**
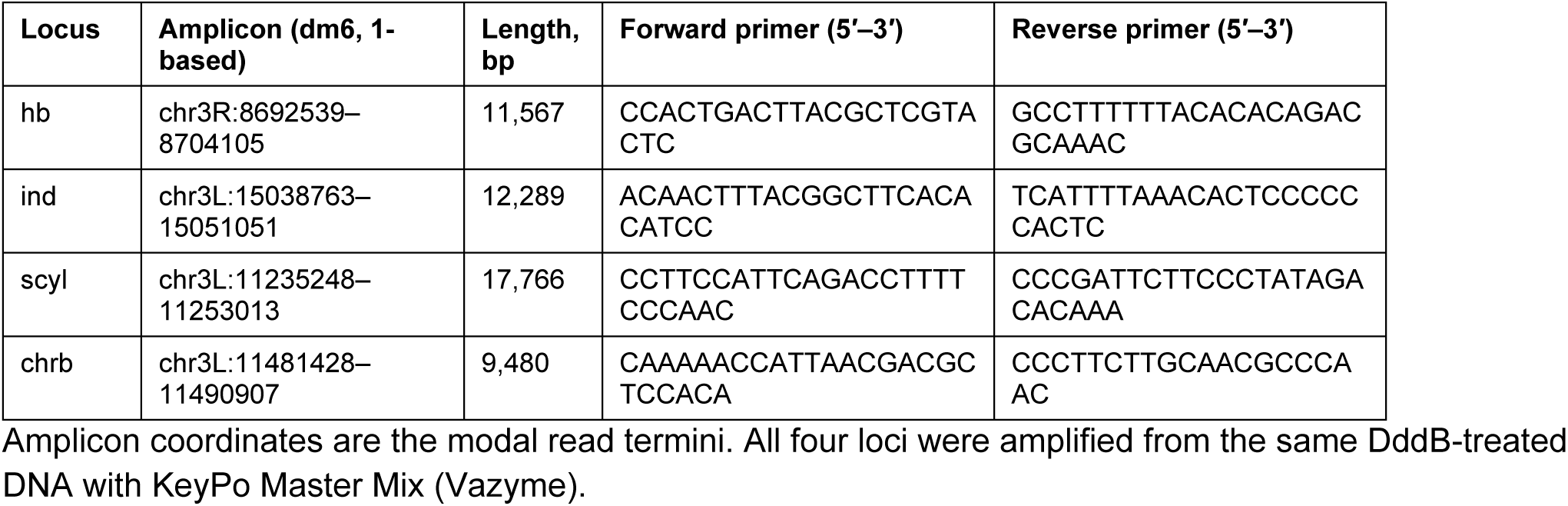
Amplicon primers.

## Supplementary Note 1. Motif footprint benchmark against ChIP-seq

Motif occupancy was compared between fiberhmm-footprint (FiberHMM 3.1.0) and fibertools footprint (ft footprint; fibertools-rs 0.13.0, commit 204f729) on fixed motif panels: 512 CTCF motifs in each of K562, GM12878, H1, H9 and HepG2, and 254–256 ChIP-blind accessible motifs each for NRF1 (JASPAR MA0506.3), REST (MA0138.3) and USF1 (MA0093.4) in K562, GM12878, H1 and HepG2 (H9 lacks ChIP-seq for these factors), selected with the CTCF rules. CTCF motifs used the literal 15-bp coordinates from the cohort catalog. ChIP labels were cell-matched ENCODE peak sets (Data availability), applied after calling. Both callers used the same FiberHMM MSPs in every cell line.

Both callers were scored on the same molecule–motif observations: molecules whose MSP spans the complete motif, the requirement ft footprint applies through its input MSPs (fiberhmm-footprint--msp-bam). ft footprint was run with one strand-oriented module per motif,--ml 125 and -F 3844; a positive was code 3, an unmodified complete module within an MSP. fiberhmm-footprint counted a molecule as bound when one FiberHMM footprint call at LLR ≥5 contained the motif within its outer edges (the widest extent consistent with the observations, bounded by the flanking m6A and stored with each call in the MA tag), no m6A (ML ≥125) fell within the motif and the conservative call width lay within the window. Site-level bound fractions, with numerators and denominators pooled across libraries, were ranked against ChIP labels by AUROC.

Without a width window this rule matched ft footprint (mean paired difference +0.008 for CTCF, +0.004 for NRF1 and 0.000 for REST and USF1, for which the two rules are equivalent; Supplementary Fig. 4d), so the width step provides the gain. Most remaining false positives at unbound motifs were broad 60–90-bp protections, which both callers count when the motif is unmethylated (Supplementary Fig. 4c,e). A label-free maximum width of 74 bp, the 95th percentile of conservative widths of FiberHMM footprint calls overlapping all benchmark motifs on MSP-eligible molecules (no labels used), raised FiberHMM above ft in every CTCF, NRF1 and REST cell line, with all paired intervals above zero (CTCF +0.012 to +0.030; NRF1 +0.030 to +0.058; REST +0.038 to +0.084). For USF1 the differences ranged from −0.005 to +0.014 and every interval included zero (Supplementary Fig. 4a and Supplementary Table 2).

ChIP calibration (fiberhmm-footprint calibrate--holdout cell) searched minimum widths of 0–80 bp and maximum widths of 10–150 bp in 5-bp steps, maximizing the mean of per-cell-line AUROCs; cell lines were not pooled because ChIP labels are cell specific. Each cell line was scored with the window learned on the other cell lines of the same factor. Held-out FiberHMM − ft differences averaged +0.031 for CTCF, +0.064 for NRF1 and +0.108 for REST, with all paired intervals above zero, and +0.014 for USF1, with every interval including zero (Fig. 1d, Supplementary Fig. 4a and Supplementary Table 2). Held-out windows were similar across cell lines (CTCF 20–60 bp, NRF1 15–55 bp, REST 0–45 bp and USF1 10–65 bp at their widest), and in-sample optimization overstated AUROC by a median of 0.010 (range 0.001–0.020). The maximum width carried the signal: AUROC peaked at maxima of 35–60 bp for CTCF, NRF1 and REST and about 65 bp for USF1, whereas minima above the motif length added little (Fig. 1e and Supplementary Fig. 4f). Windows in which FiberHMM positives made up <1% of eligible observations or occurred at <20% of motifs are shown in gray, because most motifs then score zero and AUROC reflects ties.

Confidence intervals are the 2.5th and 97.5th percentiles of 2,000 paired resamples of occupied 1-Mb genomic blocks (chromosome and floor(motif start/1,000,000)); NumPy default_rng (PCG64), seed 20260914, was reset per cell line, and the same draws were used for both callers. To test dependence on the footprint threshold, CTCF calls were re-decoded at each LLR from 2 to 8 (the decoders re-run, rather than filtering calls decoded at LLR 5 with--min-llr), with the LLR 5 eligible observations as a fixed denominator. With the 74-bp cap, AUROC varied by at most 0.012 across thresholds within each cell line (overall 0.833–0.889), and FiberHMM − ft ranged from +0.005 to +0.032 (Supplementary Fig. 4b and Supplementary Table 3).

## Supplementary Note 2. Browser display, navigation and analysis performance

### Display and navigation compared with IGV

FiberBrowser, IGV desktop 2.19.2 and igv.js 3.8.0 were measured on the same local *NAPA* inputs at 1,000, 10,000 and 50,000 molecules. Each tool, depth, sampling setting and evidence setting was tested in five serial fresh-application trials, with rotating tool order. The final comparison contains 180 planned trials; 170 completed and 10 ended in an application failure. Completed operations before a failure remain in the results. Unreached operations have no timing estimate.

Two endpoints are reported. The first visible view is the time until the first bounded viewport image was written. The all-row endpoint is the time until every eligible read had been captured across bounded image tiles and verified, which includes scrolling through every row. On a first visit to a region with all reads, FiberBrowser showed the first view sooner than IGV desktop at 1,000 molecules (0.30 versus 0.38 s, basic evidence). IGV desktop was faster at 10,000 and 50,000 molecules (1.8–2.6-fold basic, 2.5–4.8-fold rich), and the all-row endpoint was similar in the two tools (0.77–1.21-fold). On a same-process revisit, FiberBrowser showed the first view 2.5–6.6-fold sooner and completed the all-row endpoint 2.0–2.6-fold sooner. With default sampling, FiberBrowser revisits took 0.07–0.31 s regardless of depth, compared with 0.2–3.9 s for IGV desktop, whereas first visits were 1.7–3.9-fold slower than IGV desktop at 10,000–50,000 molecules (Supplementary Table 5, continued). The sampled tools drew different numbers of rows: FiberBrowser 526–572, IGV desktop up to 544 and igv.js 765–780.

Application failures are listed in Supplementary Table 4. Later operations in the same trial were not run; completed earlier operations remain in the source tables and five-repeat summaries.

### Display measurement methods

The depth-controlled BAMs were constructed from 2,971 primary mapped *NAPA* DAF-seq templates spanning the locus center. Balanced replication assigned unique names, preserved alignment positions and optional tags, and introduced no positional jitter. The tested window was chr19:47514150–47516150 (zero-based, half-open). Input hashes and the construction script accompany the source tables. These synthetic-depth inputs measure scaling of this amplicon workload.

Each application opened the BAM and local reference at an empty locus before timing. The first sequence visited the target, panned by half its width, zoomed to half its width, then zoomed to twice its width about the original center. The same sequence was repeated after visiting the empty locus in the same process. Filesystem caches remained warm. Untimed pilots populated shared compiled-code caches. Application startup is recorded separately and excluded from the display comparison.

Sampling-on trials used a 4,000-read working target in FiberBrowser, explicit IGV sampling at 100 alignment starts per 50-base window, and the igv.js default sampling depth of 1,000. Their sampling algorithms produce different cohorts; actual loaded, displayed and drawn counts are retained. The configured FiberBrowser working target does not cap its direct-read path: in the dense rich trials, this path loaded the full region before selecting displayed reads. All-read trials disabled sampling, minimum-footprint filters and terminal-gap filters. Eligible primary, mapped, nonduplicate records were counted independently from the BAM.

FiberBrowser and igv.js ran in headless Chromium 147.0.7727.15 through Playwright 1.59.1. Browser viewports were 1500×1000 logical pixels at device scale 1 with one-pixel rows. The IGV desktop frame was 1500×1000 and used its stock adaptive squished layout. All-read initial and revisit captures verified every eligible name over bounded tiles (browser axes at most 8192 pixels; IGV capture height at most 4096). Navigation captures verified the visible viewport. Stock draw calls, nonblank row pixels and PNGs provided paint evidence. Renderer implementations were unchanged.

The source tables separate target readiness, first visible PNG and completion of all-row capture with verification. Each process tree was sampled for memory every 100 ms. Peak values are summed resident-set sizes across tool and benchmark-script processes for the entire trial, including setup and both sequences. Shared pages can contribute to multiple processes; these values are not unique physical-memory allocations.

Trials requested a 16 GiB Java heap and a 16,384 MiB Chromium old-space limit. Effective heap limits are recorded separately because Chromium can impose a lower limit. Failures and subsequent unreached operations are tabulated without imputation. Medians and inclusive interquartile ranges are computed only for groups with all five repeats (Supplementary Table 5, both endpoints).

### Analyses at *NAPA*

Analysis timings used real, unreplicated *NAPA* Hia5 and SsDddA BAMs re-called with FiberHMM 3.0.0. The 340-base interval was chr19:47514980–47515320. Fixed NRF1 (47515206–47515222) and E-box (47515079–47515102) features were used for clustering, filtering and footprint co-occupancy. All analysis coordinates are zero-based and half-open. The timings characterize FiberBrowser analyses; these operations were not compared with IGV.

A fresh application was used for each assay, operation and repeat. Timing began after verified read display and two continuous seconds with no pending or new read/analysis requests; this setup wait was excluded from analysis latency. The display cap was 200 reads, row height was three pixels, and minimum-footprint, footprint-quality, spanning and terminal-gap filters were disabled. Each preview used the exact IDs verified on the canvas. Each full refinement retained the original full scope without an analysis or preparation cap. Default evidence requirements and feature-coverage rules determined eligibility.

API analyses were timed from browser request start through JSON decoding, excluding result-chart painting. Sort and clustering ran a preview followed by full refinement; their combined time is the per-trial sum of those two request durations, excluding benchmark-script overhead between requests. The full stage is therefore measured after preview. The repeated sequence ran in the same process with caches retained. Scan used only the displayed cohort.

Sort used accessibility in descending order. Filter selected reads with open accessibility over the fixed motif features. Combinatorial clustering used accessibility features with a threshold of 0.5. K-means (k=5) used TF footprint features with a 90-base subnucleosomal threshold. Footprint co-occupancy used the saved *NAPA* rules: TF footprints 10–89 bases long overlapping the NRF1 and E-box intervals, with motif discovery, sequences and profiles disabled. Exact requests are retained (Supplementary Tables 6 and 7).

All analysis trials passed validation. First/repeat API payloads and complete scientific results were compared with timing fields excluded; five-repeat stability was checked separately for each operation. Working molecule sets were retrieved only after timed requests. Full Sort included every native eligible name, full clustering included all names covering both selected features, and co-occupancy retained the full molecule set. Network timelines were checked for concurrent read or analysis requests. Raw responses and painted IDs accompany the source data.

### Reproducibility

The benchmark ran on an Apple M3 Max with macOS 26.7.1 (arm64), 16 physical CPU cores and 64 GiB RAM, with Python 3.10.13. Software was FiberBrowser 3.0.0 and FiberHMM 3.0.0. Exact source commits, Chromium, Playwright, Java and library versions, source hashes and exact input hashes are retained with the run metadata. The packaged Mac application uses its bundled Python runtime; these benchmarks ran from source.

Source tables include display measurements and outcomes for every planned operation, analysis measurements and stage timings, cohort validation, input manifests, memory traces, screenshots, exact API payloads and failure logs. Pilots are retained and excluded from formal summaries.

